# The MRi-Share database: brain imaging in a cross-sectional cohort of 1,870 university students

**DOI:** 10.1101/2020.06.17.154666

**Authors:** Ami Tsuchida, Alexandre Laurent, Fabrice Crivello, Laurent Petit, Marc Joliot, Antonietta Pepe, Naka Beguedou, Marie-Fateye Gueye, Violaine Verrecchia, Victor Nozais, Laure Zago, Nathalie Tzourio-Mazoyer, Emmanuel Mellet, Stephanie Debette, Christophe Tzourio, Bernard Mazoyer

**Affiliations:** Université de Bordeaux, Institut des Maladies Neurodégénératives, UMR5293, Groupe d’Imagerie Neurofonctionnelle, Bordeaux, France; CNRS, Institut des Maladies Neurodégénératives, UMR5293, Groupe d’Imagerie Neurofonctionnelle, Bordeaux, France; CEA, Institut des Maladies Neurodégénératives, UMR5293, Groupe d’Imagerie Neurofonctionnelle, Bordeaux, France; Fealinx and Université de Bordeaux, Ginesislab, Bordeaux, France; Université de Bordeaux, Inserm, Bordeaux Population Health Research Center, UMR1219, CHU Bordeaux, Bordeaux, France; Centre hospitalier universitaire Pellegrin, Bordeaux, France

**Keywords:** MRI, brain, student, cohort, cross-sectional, post-adolescence

## Abstract

We report on MRi-Share, a multi-modal brain MRI database acquired in a unique sample of 1,870 young healthy adults, aged 18 to 35 years, while undergoing university-level education. MRi-Share contains structural (T1 and FLAIR), diffusion (multispectral), susceptibility weighted (SWI), and resting-state functional imaging modalities. Here, we described the contents of these different neuroimaging datasets and the processing pipelines used to derive brain phenotypes, as well as how quality control was assessed. In addition, we present preliminary results on associations of some of these brain image-derived phenotypes at the whole brain level with both age and sex, in the subsample of 1,722 individuals aged less than 26 years. We demonstrate that the post-adolescence period is characterized by changes in both structural and microstructural brain phenotypes. Grey matter cortical thickness, surface area and volume were found to decrease with age, while white matter volume shows increase. Diffusivity, either radial or axial, was found to robustly decrease with age whereas fractional anisotropy only slightly increased. As for the neurite orientation dispersion and densities, both were found to increase with age. The isotropic volume fraction also showed a slight increase with age. These preliminary findings emphasize the complexity of changes in brain structure and function occurring in this critical period at the interface of late maturation and early aging.

## Introduction

There is mounting evidence indicating the importance of early life factors on cognitive status and neurological conditions later in life (Whalley et al., 2006). Genetic factors can shape early brain development as well as cognitive ageing processes through common molecular pathways (Kovacs et al., 2014), and early life conditions (e.g. pre- and postnatal environment, socioeconomic status, educational attainment) as well as lifestyle choices contribute to risk factors for cerebrovascular diseases and dementia later in life (e.g. Backhouse et al., 2017; Corley et al., 2018; Wajman et al., 2018). The vast majority of epidemiological studies investigating the risk factors for late-onset neurological conditions tend to focus either on the middle-to old-age population (e.g. Debette et al., 2011; Kivipelto et al., 2001; Whitmer et al., 2007) or on childhood (Backhouse et al., 2017; Field et al., 2016; Gluckman et al., 2008). Likewise, many large-scale neuroimaging cohort studies have charted morphological changes associated with healthy and pathological development and ageing, focusing on early and late childhood to early adulthood for development (e.g. PING, Jernigan et al., 2016; PNC, Satterthwaite et al., 2016; IMAGEN, Schumann et al., 2010; Generation R, White et al., 2013) or on middle-to late-life for ageing (Three-City, 3C Study Group, 2003; UK-Biobank (UKB), Alfaro-Almagro et al., 2018; 1000BRAINS, Caspers et al., 2014; LBC1936, Deary et al., 2007; Rotterdam, Ikram et al., 2017; EVA, Lemaître et al., 2005; LIFE, Loeffler et al., 2015; OASIS, Marcus et al., 2010; BILGIN, Mazoyer et al., 2016; SYS, Pausova et al., 2017; ADNI, Petersen et al., 2010; MAS, Sachdev et al., 2010, OATS, 2009; ASPS-Fam, Seiler et al., 2014; Framingham, Seshadri et al., 2004). Consequently, there is a relative paucity of epidemiological data and neuroimaging cohorts focusing on early adulthood, despite the significant life changes many undergo during this period, as they explore the world to attain independence and, for some, pursue higher education. While the most rapid brain changes occur during early development and the total brain size reaches adult levels by the end of childhood, both global and regional changes in brain structure and function continue throughout this period (Dumontheil, 2016). Yet, few studies investigate the impact of learning and social changes associated with higher education on maturational changes in the brain, and how it interacts with the personal traits, physical and mental health status to influence immediate as well as later-life events. Such data are crucial both for developing effective policies and interventions for promoting student health and well-being, and for gaining insight into the early life factors associated with vulnerabilities later in life.

The i-Share (for internet-based Student Health Research enterprise; www.i-share.fr) cohort project was conceived to fill this gap. It aims to evaluate important health aspects among 30,000 university students over the course of 10 years. Besides the evaluation of the frequency and impact of specific health conditions, i-Share will also allow for testing of biological mechanisms and preventive strategies for mental and physical health conditions in young adults. Launched in 2013, it has collected detailed information pertaining to personal characteristics and lifestyle habits, including risk-taking behaviours, physical activity, diet, sleep, and cognitive abilities, through web-based questionnaires, as well as medical and health status.

An important sub-component of the i-Share study, which was called “MRi-Share”, is a multi-modal brain magnetic resonance imaging (MRI) database collected in a subset of i-Share participants. The specific motivations behind MRi-Share were to 1) characterize late-maturational changes of post-adolescence brain; 2) investigate the impact of higher education on late maturational processes of the brain; 3) study the associations between brain phenotypes and neuropsychiatric conditions prevalent in young adults, such as migraine, depression and anxiety disorders, and substance abuse; and 4) establish the early occurrence of imaging biomarkers of late-life disorders, such as white matter hyperintensities (WMH) and enlarged perivascular space (ePVS). Nearly all MRi-Share participants enrolled in another closely associated i-Share sub-component, called “Bio-Share”, a biobank derived from analyses of blood samples, used to generate large scale multi-modal molecular biomarkers. Together, these two i-Share components permit the exploration of early neuroimaging-based biomarkers for late-onset neurocognitive conditions such as cerebrovascular disease and dementia.

The primary goal of the present manuscript is to describe the MRi-Share image acquisition protocol as well as the analysis pipelines used for deriving brain phenotypes from MRI. These image-derived phenotypes (IDPs) include 1) measures of both volume- and surface-based brain morphometry from structural MRI, 2) measures of the organization of the white matter microstructure organization based on diffusion MRI, and 3) measures of intrinsic functional connectivity derived from resting-state functional MRI. These phenotypes were obtained at both the brain, regional, and voxel levels. A special care has been taken to detail the quality control (QC) steps, since there is an increasing awareness of the impact of QC procedures and metrics on IDPs of morphometry (Ducharme et al., 2016; Madan, 2018; Reuter et al., 2015) and white matter properties (Roalf et al., 2016). The secondary goal is to present preliminary results on associations of some of these IDPs at the whole brain level with both age and sex, in this large cross-sectional cohort of post-adolescence individuals.

## Data Acquisition

### MRi-Share study protocol

The study protocol was approved by the local ethics committee (CPP2015-A00850-49). All participants were recruited through the larger i-Share cohort study. The i-Share participants recruited at the Bordeaux site were given the information regarding MRi-Share and Bio-Share substudies. Those interested in contributing received detailed information about both substudies, including a “virtual visit” to the MRI facility that gave the participants a better idea about what was involved, and were invited to make an appointment with one of the MD investigators. The MD investigator ensured that each participant received all the information pertinent to the participation in both studies, and also checked for the absence of any cause for exclusion. Exclusion criteria were: 1) age over 35 years; 2) pregnancy or nursing; 3) claustrophobia; and 4) contraindications for head MRI. Participants then signed an informed written consent form, and scheduled for the MRI session, after which they received compensation for their contribution.

Out of 2,000 individuals who met with the MD investigators between October 2015 and June 2017, 29 were either not willing to participate in the MRi-Share or found to be ineligible. Additional 46 withdrew from the study before scheduling the MRI session, and another 7 participants withdrew at the scheduled session, before the scanning took place. Forty-eight students had been eligible and willing to participate, but could not be scheduled before the acquisition terminated in November 2017, and therefore were never scanned. Out of the remaining 1,870 participants who underwent the scanning session, two individual datasets were removed, one because a participant withdrew from the study after being scanned, the other due to instrument failure during scanning. In addition, there were incidental findings requiring medical referral in 36 participants (see Incident findings section for details), and their imaging data were subsequently removed from further analyses, making the final sample size of 1,832.

### Demographic information

While the larger i-Share study collected a detailed socio-demographic, health and lifestyle-related information through web-based questionnaires (see Montagni et al., 2019 for examples of domains covered by the study), we report here a limited set of demographic variables collected specifically in association with the MRi-Share protocol. These include sex and age at the time of MRI acquisition, as summarized in Table 1. The table also contains the proportion of each sex for the entire i-Share cohort for comparison. It can be seen that the high proportion of females relative to males in the MRi-Share is a feature of the larger i-Share cohort itself. Actually, a higher proportion of women among University students is observed at the French national level (55%, source France Ministry for Higher Education, Research and Innovation 2019: https://cache.media.enseignementsup-recherche.gouv.fr/file/Brochures/32/8/parite2018_stats_A5_11d_908328.pdf). It is amplified in the i-Share cohort due to an over-recruitment of students coming from faculties in which even larger proportion of women are observed (medical and paramedical sciences, social sciences).

**Table 1.** Basic characteristics of the MRi-Share database. Age (mean ± SD and range, in years) of the MRi-Share participants are shown for entire sample as well as for those under 26 years old, for each sex separately and in the combined group. The proportion of each sex in the larger i-Share cohort is also shown for comparison.

|  |  | Females | Males | Total |
| --- | --- | --- | --- | --- |
| i-Share<br>(as of 11/2019) | N (%) | 13,064<br>(75.1%) | 4,323<br>(24.9%) | 17,414 |
| MRi-Share | N(%) | 1,320<br>(72.0%) | 512<br>(28.0%) | 1,832 |
|  | Age<br>(years) | 22.0 ± 2.3<br>[18.1, 34.9] | 22.3 ± 2.4<br>[18.1, 31.3] | 22.1 ± 2.3<br>[18.1, 34.9] |
| MRi-Share<br>(< 26 years old) | N (%) | 1,252<br>(72.7%) | 470<br>(27.3%) | 1,722 |
|  | Age<br>(years) | 21.7 ± 1.7<br>[18.1, 26.0] | 21.9 ± 1.8<br>[18.1, 25.9] | 21.7 ± 1.8<br>[18.1, 26.0] |

While the study protocol allowed enrolment of students up to 35 years of age, almost 95% of our sample was under 26 years old (95th percentile values for age in the entire sample = 26.3 years old; see Figure 1). We thus focused on the subsample under 26 years of age when describing the association of the MRI image derived phenotypes (IDPs) with age and sex in the sections below. Table 1 provides the demographic summary of this sub-sample as well. Males and females in our sample had a small but significant difference in their age (difference in mean age < 4 months, *p* = 0.007, *t*-test) in the entire cohort, but the difference was marginal in the sub-sample of under 26 year-olds (2 months difference in age, *p* = 0.068, *t*-test).

**Figure 1.**
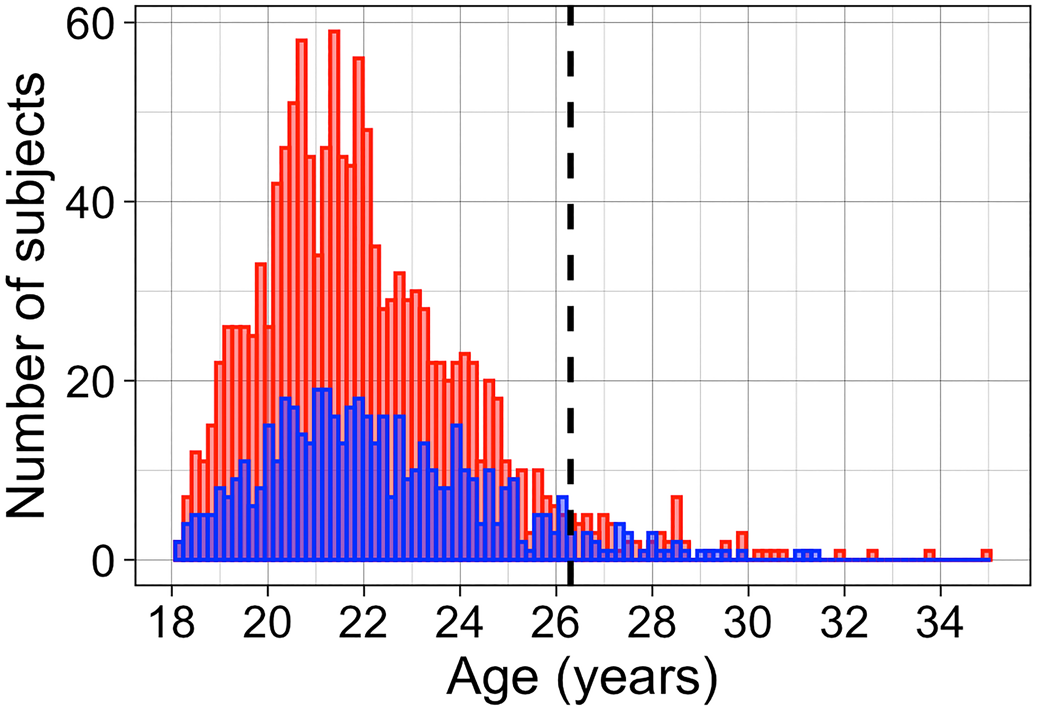
Age distribution histogram of the entire MRi-Share database. The age distribution histogram is shown for male (blue) and female (red). The dotted line indicates the 95th percentile of the distribution.

### MRI acquisition

The MRI acquisition protocol for the MRi-Share database was designed to closely emulate that of the UKB MR brain imaging study (Alfaro-Almagro et al., 2018), in terms of both modalities and scanning parameters for each. We emulated the UKB brain MRI protocol so that it would allow the combined analysis of the two databases in the future, as the early adulthood target period of MRi-Share is not covered by the UKB design which includes individuals aged over 45 years old.

There were, still, some differences between the MRi-Share and UKB neuroimaging protocols. First, we did not include task-related fMRI runs and used the time gained to extend the resting-state fMRI (rs-fMRI) acquisition duration which lasted for ~15 min (instead of 6 min in the UKB) resulting in 1,054 volumes for MRi-Share (to be compared with 490 in the UKB rs-fMRI). Another minor difference in the scanning protocol was in diffusion weighted imaging (DWI): we acquired 8, 32, and 64 directions each for *b* values 300, 1000, and 2000 s/mm^2^, respectively, while the UKB did not acquire *b* value of 300 s/mm^2^ and instead acquired 50 directions each for *b* values 1000, and 2000 s/mm^2^. We also had slightly more sets of *b* = 0 images acquired in Anterior-Posterior (AP) and the reverse PA phase encoding (8 pairs of AP and PA) than in UKB (3 pairs).

All neuroimaging data were acquired on the same Siemens 3T Prisma scanner with a 64-channels head coil (gradients: 80 mT/m - 200 T/m/s), in the 2-year period between November 2015 and November 2017.Table 2 summarizes the key acquisition parameters for each imaging modality. The whole acquisition session lasted ~45 min. Prior to the rs-fMRI acquisition, participants were instructed to “keep their eyes closed, to relax, to refrain from moving, to stay awake, and to let their thoughts come and go”, for the duration of the 15 min-long run. Compliance to the instruction was checked by the questions included in the self-report questionnaire, performed at the end of the scanning session.

**Table 2.** Summary of acquisition parameters. For each of the five modalities, the key acquisition parameters are listed. T1: T1-weighted imaging, MPRAGE: magnetization-prepared rapid acquisition with gradient echo, TR: repetition time, TE: echo time, TI: inversion time, FLAIR: Fluid-attenuated inversion recovery imaging, SPACE: sampling perfection with application-optimized contrasts using different flip angle evolutions, R: in-plane acceleration factor, PF: partial Fourier, SWI: susceptibility-weighted imaging, DWI: diffusion weighted imaging, MB: multiband factor, AP/PA: anterior-posterior/posterior-anterior; b-values are in s/mm2 and the number of directions is given in parentheses. rs-fMRI: resting-state functional magnetic resonance imaging; EPI: EchoPlanar Imaging.

| Modality | Duration | Voxel (matrix) size | Key parameters | # vol | N |
| --- | --- | --- | --- | --- | --- |
| T1 | 4:54 | 1.0x1.0x1.0 mm<sup>3</sup><br>(192x256x256) | 3D MPRAGE, sagittal, R=2,<br>TR/TE/TI=2000/2.0/880 ms | 1 | 1,832 |
| FLAIR | 5:50 | 1.0x1.0x1.0 mm<sup>3</sup><br>(192x256x256) | 3D SPACE, sagittal, R=2, PF=7/8,<br>TR/TE/TI=5000/394.0/1800 ms | 1 | 1,832 |
| SWI | 2:15 | 0.8x0.8x3.0 mm<sup>3</sup><br>(252x288x48) | 2D axial, R=2, PF=7/8, TR/TE1=24.0/9.42 ms | 1 | 1,830 |
| DWI | 9:45 | 1.75x1.75x1.75 mm<sup>3</sup><br>(118x118x84) | 2D axial, MB=3, R=1, PF=6/8,<br>TR/TE=3540/75.0 ms, fat sat,<br>b=0 s/mm<sup>2</sup> (8PA +8AP), b=300 s/mm<sup>2</sup> (8),<br>b=1000 s/mm<sup>2</sup> (32), b=2000 s/mm<sup>2</sup> (60) | 116 | 1,823 |
| rs-fMRI | 14:58 | 2.4x2.4x2.4 mm<sup>3</sup><br>(88x88x66) | 2D axial, EPI, MB=6,<br>TR/TE=850/35.0 ms, flip angle=56°, fat sat | 1,058 | 1,814 |

While all 1,832 participants completed the structural scans (T1 and FLAIR), 17 did not complete the whole scanning session (2 with missing SWI, DWI, and rs-fMRI, 15 with missing or incomplete DWI and/ or rs-fMRI) either due to anxiety attack or technical problems during scanning. The number of total images acquired for each scan (excluding the incidentals) is also listed in Table 2. For the description of each modality, we refer readers to the overview provided by Alfaro-Almagro et al. (2018).

#### Incident findings

Within days following the acquisition, T1 and FLAIR images were visually checked for quality by one of the three trained MD investigators (B.M, E.M). In the course of this quality control (QC) procedure, presence of any incident finding was recorded and paid special attention to when performing modality-specific individual QC (described in the Quality control section). In addition, any potentially harmful incident finding was sent to another neuroradiologist for a second opinion. Upon confirmation by the neuroradiologist, discovery of the incidental finding in a participant was notified to a neurologist investigator of the i-Share study (S.D) who took care of the participant information and follow-up. A total of 36 participants (1.9%) were identified as having incidental requiring medical referral and their data were excluded from analyses presented in the present manuscript. A detailed description of these incidental findings will be published in a separate report.

### Automated image analysis pipelines

The acquired data were managed and processed with the Automated Brain Anatomy for Cohort Imaging platform (ABACI, IDDN.FR.001.410013.000.S .P.2016.000.31235) which integrates processing pipelines built with a *nipype* interface (Gorgolewski et al., 2011) with the Extensible Neuroimaging Archive Toolkit (XNAT; http://www.xnat.org) database management system. Through this platform, all image processing, except rs-fMRI data, was performed using a dedicated computing cluster system composed of CPU servers, in a python 3.6.3 environment, with *ny-pipe* version 1.0.2. For running functions from the Statistical Parametric Mapping (SPM12: https://www.fil.ion.ucl.ac.uk/spm/), Matlab compiler runtime (MCR) was used (R2010A, v713, The Mathworks, Natick, MA). Processing of the rs-fMRI data was computed on CURTA, a shared computing cluster provided by the Mésocentre de Calcul Intensif Aquitain (MCIA) dedicated to intensive parallel computation. The python (3.6.5) and *nipype* (1.0.2) environment, and the resting-state pipeline contained in the ABACI platform were packaged on a single “resting state” singularity container (Kurtzer et al., 2017), together with all other tools and softwares the pipeline depended on. It used the same MCR v713 when running any functions from SPM12, but used MCR R2018a v94 when running other custom functions in Matlab. The main ABACI image analysis pipelines are briefly described below and detailed in the supplementary material section.

#### Image anonymization

In order to protect the anonymity of the participants, the high-resolution anatomical images (T1 and FLAIR) were processed with a defacing pipeline that masked out voxels in the facial region. This pipeline is a Nipype implementation of defacing protocol used by the UKB (Alfaro-Almagro et al., 2018), and uses the same face mask in MNI space available at the official code repository of the UKB (under templates folder in https://git.fmrib.ox.ac.uk/falmagro/UK_bio-bank_pipeline_v_1/tree/master).

#### T1 and T2-FLAIR structural pipeline

Our structural pipeline processed T1 and FLAIR images for multi-channel volume- and surface-based morphometry, primarily with SPM12 (https://www.fil.ion.ucl.ac.uk/spm/) and Freesurfer v6.0 (http://surfer.nmr.mgh.harvard.edu/). It also produced bias-field corrected and ‘cropped’ T1 images with reduced amount of non-brain tissues to be used as a reference image for all other modalities. Details of the proposed dual structural pipelines are described in the Supplementary Material, with the schematic representation of the pipeline in Supplemental Figure 1. Of note, we had initially used default parameters for SPM12-based unified tissue-segmentation and normalization of the structural images, using the cohort-specific template; however, the early visual QC of the tissue segmentation outputs revealed wrong cortical ribbon extraction leading to a underestimation of grey matter in the vast majority of participants (Supplemental Figure 2 and 3). We therefore modified the SPM12 default settings to overcome this grey matter underestimation, as described in detail in the Supplementary Material.

Global IDPs derived with this pipeline were: total grey matter volume (GM) and white matter volume (WM), mean cortical thickness (CT), total cortical inner surface defined by grey-white matter interface (inner CSA), and total cortical pial surface area (pial CSA). Although only explicitly discussed in a few surface-based morphometric studies (e.g. Hogstrom et al., 2013; Storsve et al., 2014; Tamnes et al., 2017), CSA is usually defined at the GM/WM boundary, representing inner, white surface, since the pial surface defined by the GM/cerebrospinal fluid (CSF) boundary in theory may be more sensitive to changes in CT (Winkler et al., 2012). Here, to thoroughly describe the age-related variation in GM morphometry, we also computed the total pial surface area. Distributions of these metrics in the entire sample (*N* = 1,832) are shown in Supplemental Figure 15. Overall, the IDPs from this pipeline include:

- 3 global Freesurfer surface-based IDPs (mean CT, total inner CSA, total pial CSAl)
- 1,390 regional Freesurfer surface-based IDPs (CT, inner CSA, pial CSA, number of vertices, and cortical volume for right and left cortical regions as defined by three cortical atlases included with Freesurfer package: Desikan-Killiany (34 regions: Desikan et al., 2006), DKT (31 regions: Klein and Tourville, 2012), and Destrieux (74 regions: Destrieux et al., 2010) cortical atlases)
- 31 global Freesurfer volume-based IDPs (17 global measures of volume contained in ‘aseg. stats’ table, 14 of which are described in https://surfer.nmr.mgh.harvard.edu/fswiki/MorphometryStats, plus 17 additionally calculated global measures of volume based on ‘aseg’ segmentation, as described in Supplementary Material)
- 34 regional Freesurfer volume-based IDPs (volumes for each region in ‘aseg’ labels, but excluding 7 ventricular regions that are aggregated in one of the global measures)
- 5 Freesurfer brainstem substructure volume IDPs (medulla oblongata, pons, superior cerebellar peduncle, and whole brainstem: Iglesias et al., 2015b)
- 26 Freesurfer hippocampal subfield volume IDPs (left/right parasubiculum, presubiculum, subiculum, cornu ammonis (CA) 1, CA2/3, CA4, granule cell and molecular layer of the dentate gyrus (GC-ML-DG), molecular layer, HATA, fimbria, hippocampal tail, hippocampal fissure, and whole hippocampus: Iglesias et al., 2015a)
- 3 tissue volume IDPs from SPM (total GM, WM, and CSF volumes)
- 138 regional GM volume IDPs based on SPM (total GM volume within regions defined by Harvard-Oxford cortical and subcortical atlases (Desikan et al., 2006; Frazier et al., 2005; Goldstein et al., 2007; left/right 48 cortical and 7 subcortical regions, and brainstem: Makris et al., 2006), and by Diedrichsen probabilistic cerebellar atlas (27 regions: Diedrichsen et al., 2009)

#### Fieldmap generation pipeline

As in the UKB (Alfaro-Almagro et al., 2018), we estimated the fieldmap images from the b=0 images with opposing AP-PA phase-encoding directions from DWI scans, rather than from “traditional” fieldmaps based on dual echo-time gradient-echo images. The Supplementary Material section describes the details of this pipeline, with the schematic representation of the pipeline in Supplemental Figure 5. The main outputs are: 1) the fieldmap phase and magnitude images in native T1 structural space that are used for EPI unwarping in the rs-fMRI pipeline; 2) the acquisition parameters, the field coefficient image and top-up movement parameters, and the brain mask generated from the average distortion-corrected b0 maps that are used for eddy-current and top-up distortion corrections in the DWI and rs-FMRI pipelines. This pipeline was built primarily using tools available from FMRIB Software Library (FSL, v5.0.10: https://fsl.fmrib.ox.ac.uk).

#### Diffusion MRI pipeline

The preprocessing steps of DWI are described in the Supplementary Material section, and the Supplemental Figure 6 shows the schematic representation of the pipeline. Briefly, following the eddy current and top-up distortion correction and denoising of the DWI data, the resulting image was then used to fit 1) DTI (Diffusion-Tensor Imaging; Basser et al., 1994) modelling and 2) microstructural model fitting with NODDI (Neurite Orientation Dispersion and Density Imaging; Zhang et al., 2012). The preprocessing and DTI fitting was performed using tools from FSL and the *dipy* package (0.12.0, https://dipy.org; Garyfallidis et al., 2014), while the AMICO (Accelerated Microstructure Imaging via Convex Optimization) tool (Daducci et al., 2015) was used for NODDI fitting. Here we report global DTI/NODDI statistics measured for each individual within a cerebral WM mask defined using the Freesurfer-segmented WM further refined using the SPM12-based WM probability map with a 0.5 lower threshold. This masking procedure ensured that the mean DTI/NODDI values were computed within cerebral WM regions with limited partial volume effects.

The global IDPs from this pipeline that are reported in this paper are mean DTI and NODDI metrics within the cerebral WM mask. Specifically, these metrics are: fractional anisotropy (FA), mean, axial, and radial diffusivity (MD, AD, and RD), based on DTI modeling, neurite density index (NDI), orientation dispersion index (ODI), and isotropic volume fraction (IsoVF), derived from NODDI modeling. Distributions of these metrics in the entire sample with complete DWI data (*N* = 1,823) are shown in Supplemental Figure 16. Figure 3 shows the group average maps of these DTI and NODDI metrics in standard MNI space. The following list summarizes these and other IDPs from this pipeline, and the Supplementary Material describes how they were generated:

- 77 DTI/NODDI regional WM IDPs based on subject-specific masks generated using Freesurfer aseg labels (mean FA, MD, AD, RD, NDI, ODI, IsoVF within total and left/right cerebral, cerebellar, and ventral diencephalon WM, and within corpus callosum and brainstem)
- 28 DTI/NODDI regional GM IDPs based on subject-specific masks generated using Freesurfer aseg labels (mean DTI/NODDI metrics within cortical, hippocampal, subcortical (excluding hippocampi), and cerebellar GM)
- 525 DTI/NODDI regional WM IDPs based on subject-specific masks generated using Freesurfer wmparc labels (mean DTI/NODDI metrics within 75 WM parcellated regions of the wmparc atlas (5 corpus callosal subregions and 35 left/right WM regions: Salat et al., 2009)
- 336 DTI/NODDI regional WM IDPs based on spatially normalized DTI/NODDI maps and JHU ICBM-DTI-81 white matter labels atlas (6 bilateral and 21 left/right WM tracts: Mori et al., 2008; Oishi et al., 2008)
- 36 DTI global WM skeleton IDPs based on spatially normalized WM skeleton created using TBSS (Tract-Based Spatial Statistics, Smith et al., 2006), part of FSL package (Smith et al., 2004) (mean, standard deviation, and peak-width 90 of the skeleton, or the 95th to 5th percentile value over the WM skeleton, as described in Baykara et al. (2016) for the 4 DTI metrics, as well as the same metrics computed over the left and right WM skeletons, as described in Beaudet et al. (2020))

#### Resting-state fMRI pipeline

Processing of rs-fMRI is described in the Supplementary Material section, with the Supplemental Figure 7 showing the schematic representation of the preprocessing steps. Briefly, the distortion-corrected rs-fMRI data were aligned to the T1-weighted individual reference space, band-pass filtered to a frequency window of 0.01 - 0.1 Hz, then warped into the stereotaxic space at a voxel sampling size of 2×2×2 mm^3^. These preprocessing steps were primarily performed using tools from FSL (v5.0.10: https://fsl.fmrib.ox.ac.uk) and AFNI (v18.0.05: https://afni.nimh.nih.gov, Cox, 1996). The following IDPs were then generated (details in the Supplemental Material) from the preprocessed data:

- 2 regional intrinsic connectivity (IC) matrices using the 384 regions of the AICHA atlas (Atlas of Intrinsic Connectivity of Homotopic Areas; Joliot et al., 2015), either with or without global signal removal;
- A ReHo map (Zang et al., 2004) that measures the homogeneity of local connectivity;
- An ALFF map that depicts the local amplitude of low frequency fluctuations (Yang et al., 2007)
- A map of fALFF (Zou et al., 2008), a normalized value of ALFF that improves sensitivity and specificity in detecting intrinsic brain activities;
- subject-specific IC components and their time series based on subject-level independent component analysis (ICA; Beckmann and Smith 2004).

### Quality control

#### Overview of the QC strategy

The quality of acquired MR images and phenotypes obtained from them was ensured by a 5-stage quality control processes (Figure 2) : 1) Aborting or postponing the MRI scan during acquisition if the MR technician observes significant amount of artefacts due to motion or any technical problems; 2) Reviewing of acquired structural scans (T1 and FLAIR) by experts to check for any significant artefact or presence of anomalies (see the section on Incidentals and anomalies above, and see Supplemental Figure 8 for examples of the flagged artefact); 3) Generation and reviewing of modality- and processing-specific QC images for each participant, organized as static web-pages (Supplementary Material for description and examples from each pipeline); 4) Examination of the distributions of both QC metrics and IDPss for detection of outliers using interactive web-based figures with links to subject-specific QC web-pages (Supplementary Material for the description and distributions of IDPs and quantitative QC metrics for each pipeline); and finally 5) Detailed review of relevant raw and processed data when the images were flagged as having visible artefact or benign anomalies (i.e. not classified as incidentals) in step 2 or when any metrics computed in step 4 indicated potential problems in specific processing pipeline (e.g. review of tissue segmentation images for individuals with visible artefacts, known structural anomaly, or outlier values in specific morphometric values).

**Figure 2.**
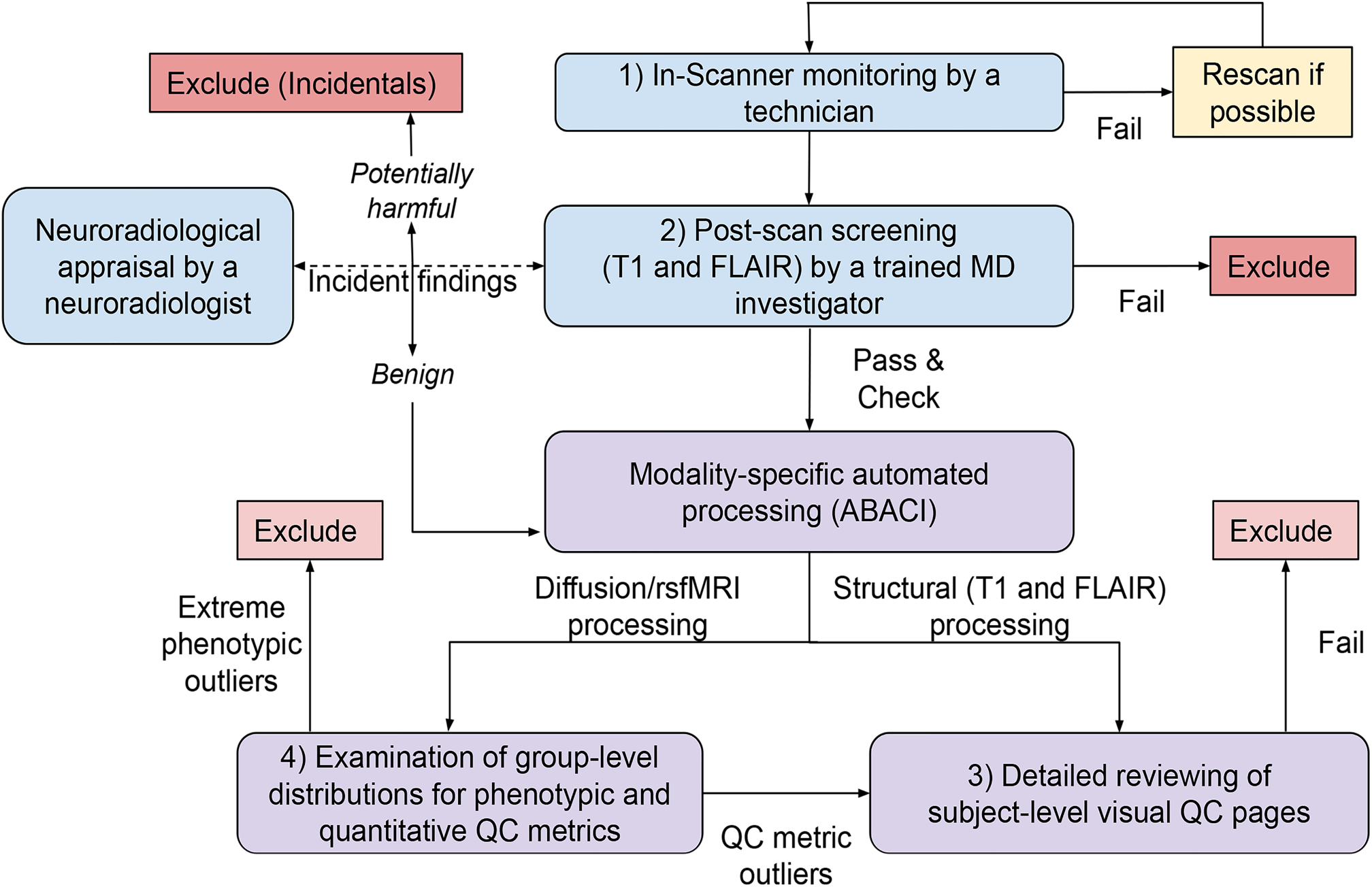
Overview of the QC workflow. Boxes in light blue represent QC steps that deal with raw data during and after acquisition, and those in light purple represent QC steps that use processed images via automated processing pipelines. While all MRI data from subjects with incidental findings were excluded (dark pink), those with processing-specific problems were excluded only from relevant analyses (light pink). Since the structural images, in particular T1 scans, served as a reference in other processing pipelines, those with severe artefacts in raw T1 images were removed from all analyses. The flowchart is loosely based on Backhausen et al. (2016), who described a recommended QC workflow for structural (T1) processing.

Since the structural scans, in particular T1 image, serve as a reference for all other modalities, we paid additional attention at checking the quality of their acquisition and processing, subjecting them to more individual individual-level QC review at step 2, in addition to the thorough review of raw acquired data at step 1. This is also because it is relatively easier to spot processing errors more objectively in the structural processing (e.g. segmentation failures) in individual-level QC images than in the DWI and rs-fMRI pipelines, where the accuracy of final output images cannot be easily assessed visually. However, whenever the group-level QC metric distributions indicated potential problems in the specific processing pipeline, individual-level QC images in the DWI and rs-fMRI pipelines were reviewed to check for any obvious problems in image quality and/or processing error. No manual editing or subject-specific parameter adjustment was performed for any of the pipelines, in order to ensure the repeatability of our results. In this paper, we primarily use the distribution of phenotypic variables of interest to define the outliers to be excluded in the analyses. Specifically, we checked for extreme outliers in each phenotypic variables using interquartile range (IQR), or Tukey fence method (Tukey, 1977), defining those with values below 3*IQR from the first quartile or above 3*IQR from the third quartile as the “far out” outliers. The distributions of all the quantitative QC metrics are reported in the Supplementary Material.

### Summary of modality-specific QC criteria and excluded participants

#### Structural pipeline

A total of 19 participants were flagged as having visible artefacts in raw T1 and/or FLAIR images by one of the three MD experts who reviewed raw images soon after the acquisition (step 2). However, none of the images had artefacts severe enough to exclude participants immediately (see Supplementary Material for the rating of artefact severity), partly because the first step in our QC protocol prevented unusable data from entering the database. We followed the recommendations and procedures proposed by (Backhausen et al., 2016), to carefully follow the impact of these flagged artefacts at each step of the structural processing, and found that all of the flagged images could be processed without any major issues. For structural pipeline, an independent trained rater (N.B) also reviewed subject-specific QC images generated from both SPM12 and Freesurfer streams for all participants, regardless of the artefact flagging, and found all 1,832 participants to have acceptable processing of T1 and FLAIR images, at least with regard to global morphometry reported here. The Supplemental Material Figure 15 shows the distributions of all the global morphometric variables we report in the current paper (CT, inner CSA, pial CSA, GM and WM volumes). None of these variables had extreme outliers in the entire sample (*N* = 1,832), nor in those under 26 years old (*N* = 1,722), who were used to examine the age and sex effects in the current paper. While most of the QC for the structural pipeline was performed by visual inspection of various images, we also computed some quantitative QC metrics and report their distributions in the Supplemental Figure 11.

#### Diffusion pipeline

For the diffusion pipeline, 9 participants had either missing or incomplete DWI scans due to technical problems or anxiety attacks, resulting in a total of 1,823 DWI data that could be processed. A number of quantitative QC metrics were reviewed for any outliers, and subject-specific visual QC images were reviewed by experts (L.P and A.T) for those outliers to detect any obvious problems with the quality of the acquired DWI or with any specific processing steps (see Supplementary Material for details for the distributions of all the quantitative QC metrics), but none was detected. There were also no extreme outliers in any of the mean DTI and NODDI metrics within the cerebral WM in the entire sample (*N* = 1,823), nor in those under 26 years old (*N* = 1,714).

#### Resting-state fMRI pipeline

The rs-fMRI scan, which was the last of the MRI session, was either missing or incomplete in 17 participants. One additional participant was found to have a wrong parameter for TR, and subsequently removed from the analysis. As a result, there was a total of 1,814 data that could be processed in the pipeline. Similarly to the diffusion pipeline, the QC for the resting-state fMRI focused on the identification of outliers on a number of quantitative QC metrics automatically generated by the pipeline, followed by the reviewing of subject-specific QC images by experts (M.J and M-F.G) to determine the acceptability of the data (see Supplementary Material for details for the distributions of all the quantitative QC metrics).

## Statistical Analysis

### MRi-Share study protocol

While the primary goal of the present manuscript is to describe the MRi-Share image acquisition protocol as well as the analysis pipelines used for deriving brain phenotypes from MRI, a secondary goal is to present preliminary results on associations of some of these brain image-derived phenotypes (IDP) at the whole brain level with both age and sex in yound adults.

As mentioned earlier, the vast majority of MRi-Share participants are aged between 18 and 26 years, with only about 5% being aged between 26 and 35 years. Rather than making an inference about age-related variations for the age-range of the entire sample that would rely on a small number of participants for the later age range, we focus here on the earlier age range of 18 to 26 years old where we have good coverage of the age span. Given the narrow target age range, we expected most of the age-related variations that exist in the IDPs to be captured by a linear age model. Indeed, preliminary comparison of models that included quadratic age effect to capture any non-linear trend did not improve model fit relative to linear age effect models in any of the IDPs examined, as judged by the Bayesian Information Criterion (BIC). Also using BIC, we examined linear age models with or without including the estimate of intracranial volume (eTIV) from Freesurfer as a covariate for the structural morphometric analyses, to check for the significant contribution of the eTIV in explaining the variance of the IDPs. For the mean DTI and NODDI metrics within cerebral WM, instead of eTIV, we examined the contribution of the volume of the cerebral WM mask as the latter has been shown to modulate the estimated DTI mean values by affecting the amount of partial volume effects (PVE) within the mask (Vos et al., 2011). We certainly minimised the PVE by restricting the mean metric computation in the Freesurfe WM mask voxels having a WM tissue probability larger than 0.5. Nevertheless, including PVE as a covariate in the statistical analysis allowed us to verify that any residual PVE did not affect the estimates for the age effect.

Given the large disparity in the sample size of each sex, we studied age effects first separately for men and women first, then as acombined group to test for the main effect of sex as well as any interaction between sex and age. Thus, for sex-specific analyses, we examined two models as described in the following equations:

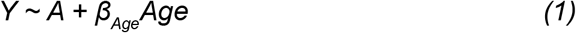

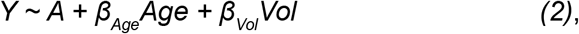

where *Vol* represents eTIV for the structural morphometry IDPs and the cerebral WM mask volume for the DTI and NODDI IDPs. Comparing these two models, we primarily report the model showing optimal fit, i.e. having the lowest BIC. For the combined group analyses, the following two models were considered:

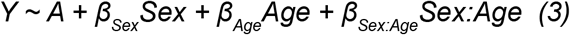

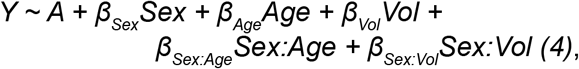

and we report the model parameter estimates with (3) or without eTIV or WM mask volume *(4)* when the best fit models for two sexes agreed (which should be chosen if we were to use BIC on the combined group models), and the results with *Vol (4)* when eTIV or the WM mask volume explained a significant amount of variance in at least one of the sexes. This was to avoid biasing the model selection for the combined group towards the larger sample size female group, when the selected model differs between the two sexes. Similarly, in the model *(4)*, we included a *Sex* by *Vol* interaction in the model to allow for a possible difference between the groups in the slope of the IDPs versus Vol, again avoiding biasing the fit towards the larger female group (Nordenskjöld et al., 2015). The interaction term with *Sex* and *Age* was included in both models to test for any sex differences in the age-related trajectory.

Both *Age* and *Vol* were mean-centered in respective groups (i.e. using female- or male-specific means in sex-specific analyses and combined mean in combined group analyses). Our aim in presenting these analyses is to describe the overall age trajectory of these global metrics inferred from the age-related variations in our dataset, rather than to test any specific hypotheses. Thus, we present the raw model fits with unadjusted *p*-values. Although we primarily report the results of the model selected by the procedure described above, we also provide the alternative model results in the Supplementary Material, given the inconsistencies in how global brain size is corrected (or not corrected) in prior studies investigating brain morphometric changes during development and maturation (e.g. Table 2 in a recent review by Vijayakumar et al., 2018). As for the DTI and NODDI IDPs, the effects of the mask volume during development are largely unknown.

All model fits were performed using *Im* function as implemented in *stats* library included with R version 3.4.4 (R Core Team, 2018), and the effect size estimates for each covariate in the model were computed as partial *η^2^* using *‘etasq’* function in the R *heplots* library (Fox et al., 2018). BIC was computed using the *‘extractAIC* function in the R stats package, with the weight of the equivalent degrees of freedom (*k*) set to log(*n*), where n represents the sample size in the mode. In the combined-sex analyses, we report the estimates of *β’s* for sex and sex interaction terms using the default contrast setting in R that treats one of the groups (female in our case) as a reference, so that the *β* represents the difference between male and female groups. However, to present the age effect across groups, we report *β* for age effect by setting the contrast to ‘contr.sum’, that gives orthogonal contrasts where the effect estimates for non-categorical variables (i.e. age in our case) represent those for the overall group mean.

## Results

Figure 3 and 4 illustrate the spatially normalized average maps of various morphometry and white matter property IDPs across the entire MRi-Share sample (*N* = 1,832 for Figure 3, *N* = 1,823 for Figure 4). Figure 3A and B show the sample average images of spatially normalized T1 and FLAIR. Figures 3C, D, and E depict the vertex-level sample average of CT, inner CSA and pial CSA computed in the *fsaverage* surface-based template space. Figures 3E and 3F show the sample average GM and WM tissue probability maps and Figure 4 the sample average DTI and NODDI maps.

**Figure 3.**
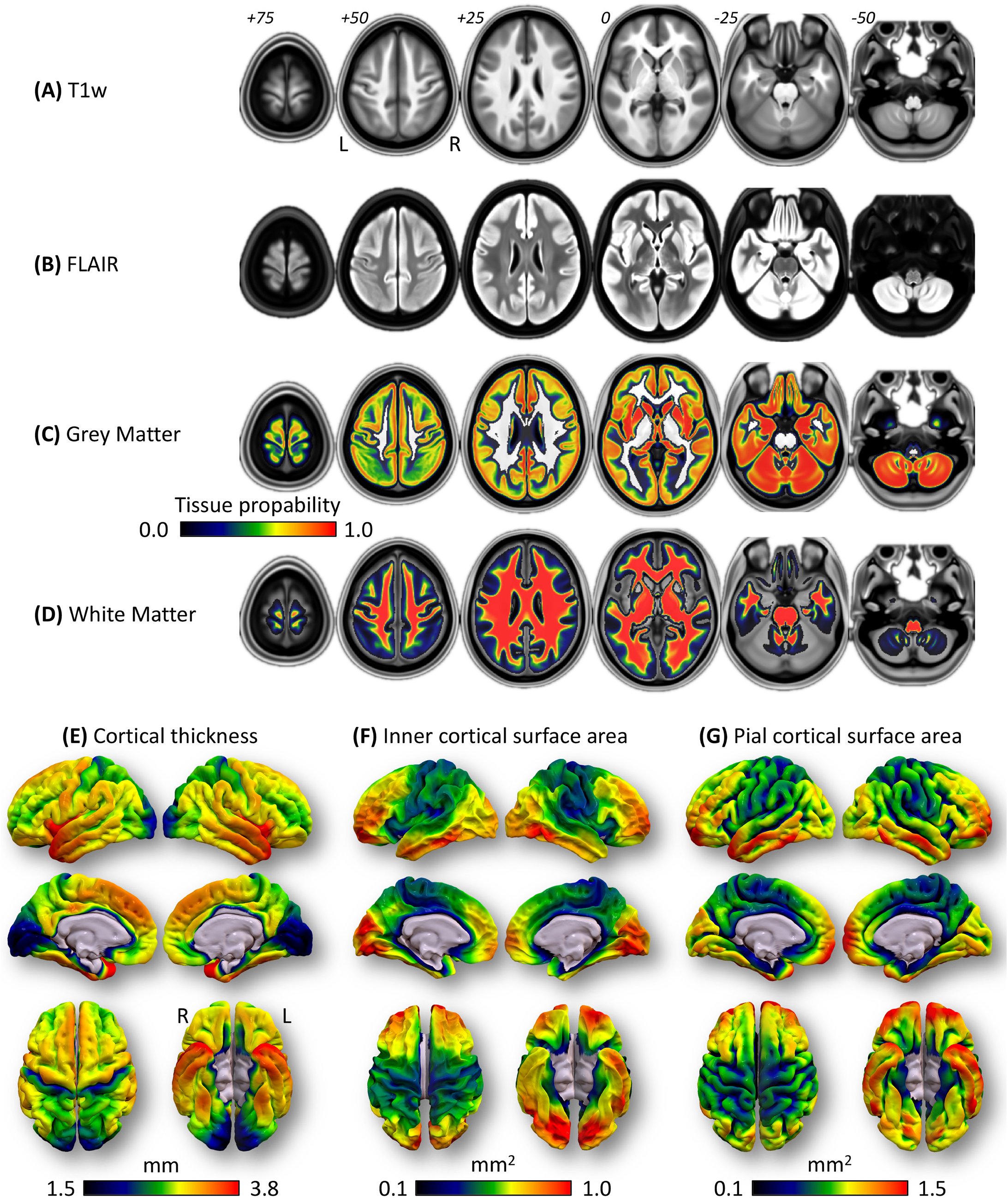
Group average maps for structural images for 1,832 MRi-Share subjects. Average maps across subjects are shown for (A) T1, (B) FLAIR, (C) mean CT, (D) total inner CSA, (E) total pial CSA, (F) GM tissue map, and (G) WM tissue map. Volumetric images (A, B, F, G) are spatially normalized and in standard MNI space. The tissue probability maps (F and G) are overlaid on the average T1 image, and the color bar at the bottom indicates the group average tissue probability. All volumetric maps were with the MRIcron (v1.0.20190902; https://people.cas.sc.edu/rorden/mricron/). Surface-based metrics (C to E) are projected onto *fsaverage* template space, with (C) and (E) projected onto pial surface, and (D) onto white surface of the template, visualized using the Suf Ice (v1.0.20190902; https://www.nitrc.org/projects/surfice/).

**Figure 4.**
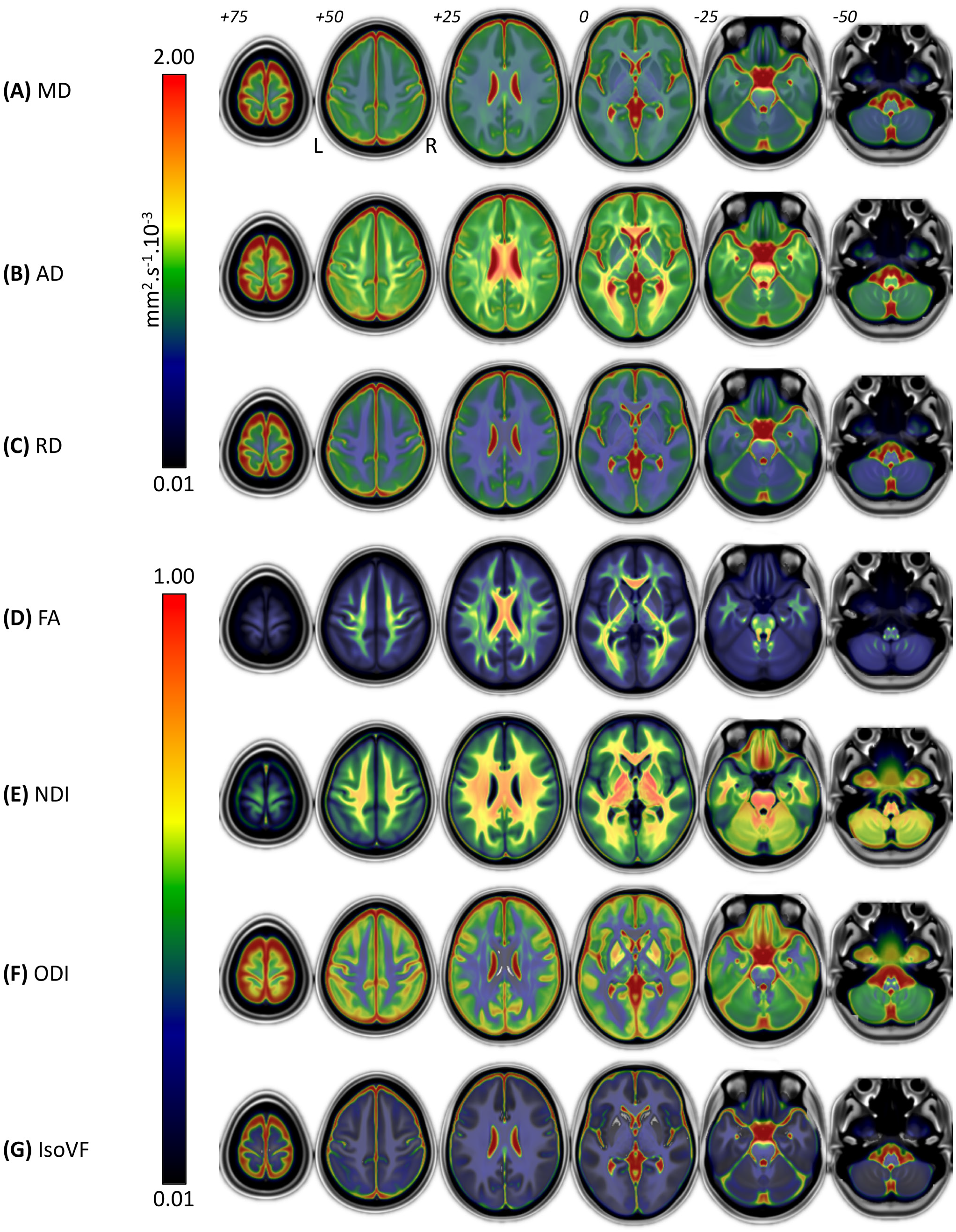
Group average maps for DTI and NODDI metrics in 1,823 MRiShare subjects. Average maps of (A) FA, (B) MD, (C) RD, and (D) AD from the DTI modeling, and (E) NDI, (F) ODI, and (G) IsoVF from the NODDI modeling across subjects are shown in the standard MNI space. Colorbar for the FA, NDI, ODI, and IsoVF is shown at the left-bottom, and that for the MD, RD, and AD is shown at the right-bottom of the figure. All maps were visualized with the MRIcron (v1.0.20190902; https://people.cas.sc.edu/rorden/mricron/).

### Global grey matter morphometry

Table 3 summarizes the descriptive statistics and the age and any sex differences in the three global cortical GM morphometry: Freesurfer-based mean CT, total inner CSA, total pial CSA, and SPM-based total GM volume, the latter being calculated from jacobian modulated and warped GM maps. The age and sex effect estimate on these metrics are also reported in Table 3 and plotted on Figure 5 for surface-based metrics, and Figure 6A for GM volume.

**Table 3.** Summary of grey matter morphometry. Descriptive statistics and the results of linear model fit for mean CT, total inner CSA, and total pial CSA from Freesurfer v6.0, as well as total GM volume from SPM12, are shown. Parameter estimates (*β*) for age and sex are shown, together with 95% confidence intervals (CI) for each *β*, as well as uncorrected *p* values and partial eta squared as the effect sizes for each variable. The Model column indicates the selected model (see text) with the corresponding model number as described in Methods section. The interaction between sex and age was tested in the combined group but was not significant in any of the metrics listed here, and thus is not included in the table. The variance explained by eTIV for models that included it is discussed in the text. Overall model fit is indicated as adjusted squared R values.

| | | Mean ± SD [min, max] | | Age | | | Sex (M>F) | | | |
| --- | --- | --- | --- | --- | --- | --- | --- | --- | --- | --- |
| | *N* | | Model | $\beta$ [95% CI] | $p$ | Partial $\eta^2$ | $\beta$ [95% CI] | $p$ | partial $\eta^2$ | adj-R<sup>2</sup> |
| **CT** | | (mm) | | ($\times 10^{-3}$ mm/yr) | | | ($\times 10^{-3}$ mm) | | | |
| Males | 470 | 2.86 ± 0.08 [2.60, 3.10] | (1) | -2.49 [-6.47, 1.49] | 0.22 | 0.003 | -- | -- | -- | 0.001 |
| Females | 1252 | 2.84 ± 0.07 [2.61, 3.11] | (1) | -4.58 [-6.96, -2.19] | 1.7$\times 10^{-4}$ | 0.011 | -- | -- | -- | 0.010 |
| Combined | 1722 | 2.84 ± 0.08 [2.60, 3.11] | (3) | -3.53 [-5.77, -1.29] | 0.0020 | 0.008 | 23.7 [15.6, 31.8] | 1.0$\times 10^{-8}$ | 0.019 | 0.025 |
| **inner CSA** | | ($\times 10^5$ mm<sup>2</sup>) | | ($\times 10^2$ mm<sup>2</sup>/yr) | | | ($\times 10^2$ mm<sup>2</sup>) | | | |
| Males | 470 | 1.86 ± 0.14 [1.48, 2.30] | (2) | -4.01 [-7.53, -0.11] | 0.026 | 0.011 | -- | -- | -- | 0.738 |
| Females | 1252 | 1.68 ± 0.12 [1.31, 2.22] | (2) | -1.63 [-3.53, 0.27] | 0.093 | 0.002 | -- | -- | -- | 0.767 |
| Combined | 1722 | 1.73 ± 0.15 [1.31, 2.30] | (4) | -2.82 [-4.67, -0.97] | 0.0029 | 0.004 | -12.9 [-22.6, -3.2] | 0.0086 | 0.006 | 0.826 |
| **pial CSA** | | ($\times 10^5$ mm<sup>2</sup>) | | ($\times 10^2$ mm<sup>2</sup>/yr) | | | ($\times 10^2$ mm<sup>2</sup>) | | | |
| Males | 470 | 2.27 ± 0.16 [1.84, 2.73] | (2) | -7.23 [-11.2, -3.25] | 3.9$\times 10^{-4}$ | 0.027 | -- | -- | -- | 0.746 |
| Females | 1252 | 2.05 ± 0.14 [1.60, 2.62] | (2) | -4.33 [-6.55, -2.11] | 1.3$\times 10^{-4}$ | 0.012 | -- | -- | -- | 0.768 |
| Combined | 1722 | 2.11 ± 0.18 [1.60, 2.73] | (4) | -5.78 [-7.92, -3.65] | 1.2$\times 10^{-7}$ | 0.016 | -5.5 [-16.6, 5.6] | 0.33 | 0.002 | 0.832 |
| **GM** |  | (cc) |  | (cc/yr) |  |  | (cc) |  |  |  |
| Males | 470 | 730.6 ± 54.7 [598.9, 912.8] | (2) | -2.03 [-3.22, -0.84] | 9.0$\times 10^{-4}$ | 0.023 | -- | -- | -- | 0.808 |
| Females | 1252 | 659.0 ± 48.9 [495.4, 824.1] | (2) | -2.38 [-3.10, -1.65] | 2.1$\times 10^{-10}$ | 0.032 | -- | -- | -- | 0.785 |
| Combined | 1722 | 678.5 ± 59.8 [495.4, 912.8] | (4) | -2.20 [-2.88, -1.52] | 2.9$\times 10^{-10}$ | 0.029 | -7.0 [-10.6, -3.5] | 1.0$\times 10^{-4}$ | 0.009 | 0.851 |

**Figure 5.**
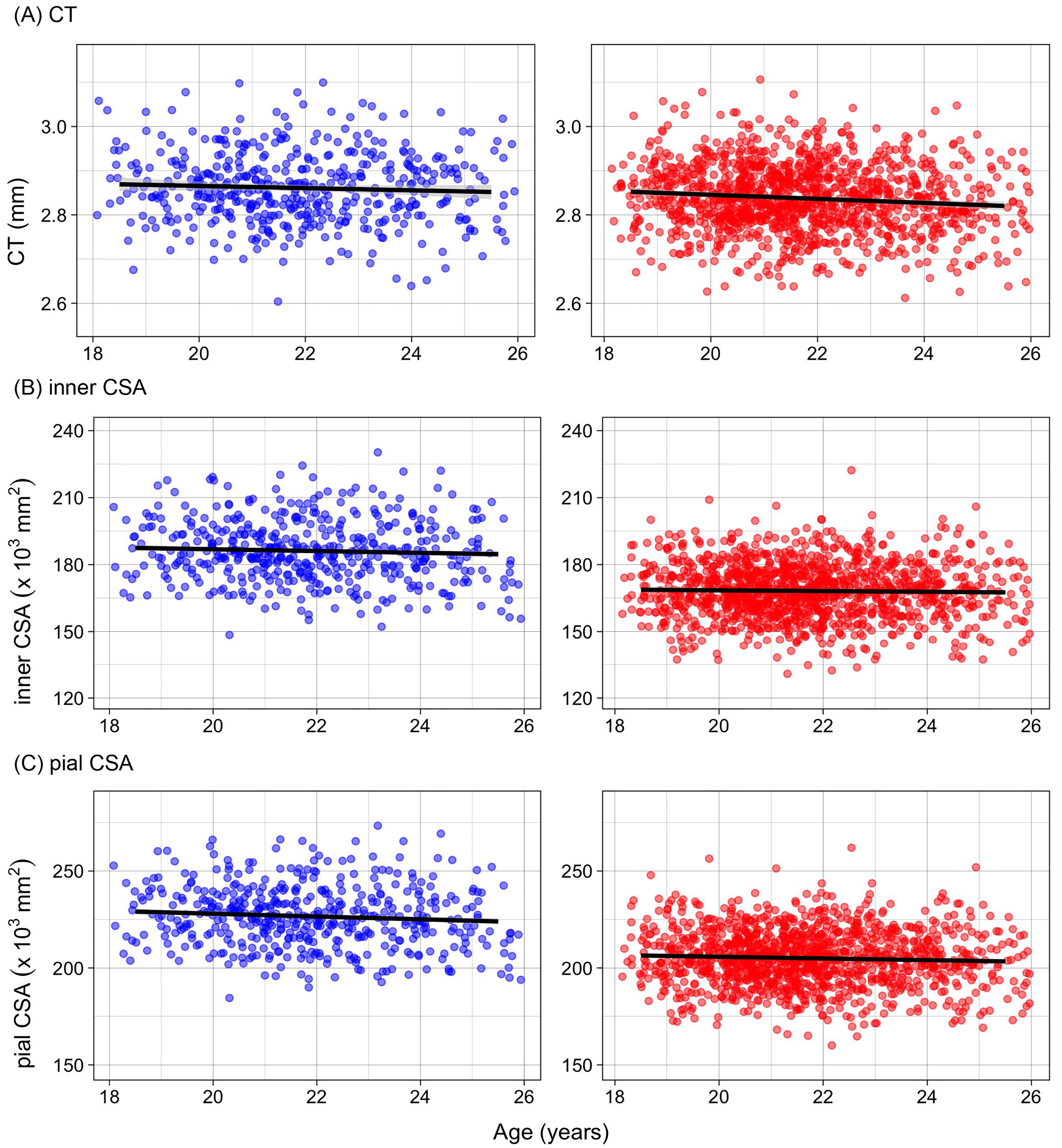
Age-related variations in surface-based morphometry. (A) The mean CT, (B) total inner CSA, and (C) total pial CSA are plotted against age, with individual subjects represented as scatter points (left panel: male, right panel: female). For each sex, predicted age trajectories, based on the model selected by BIC (see text) in the sex-specific analyses are shown. The plots in the right panel show predicted trajectories for hypothetical male or female subjects with identical, global mean eTIV, based on combined group analyses. The 95% confidence intervals of the predictions are represented as shades around the line.

**Figure 6.**
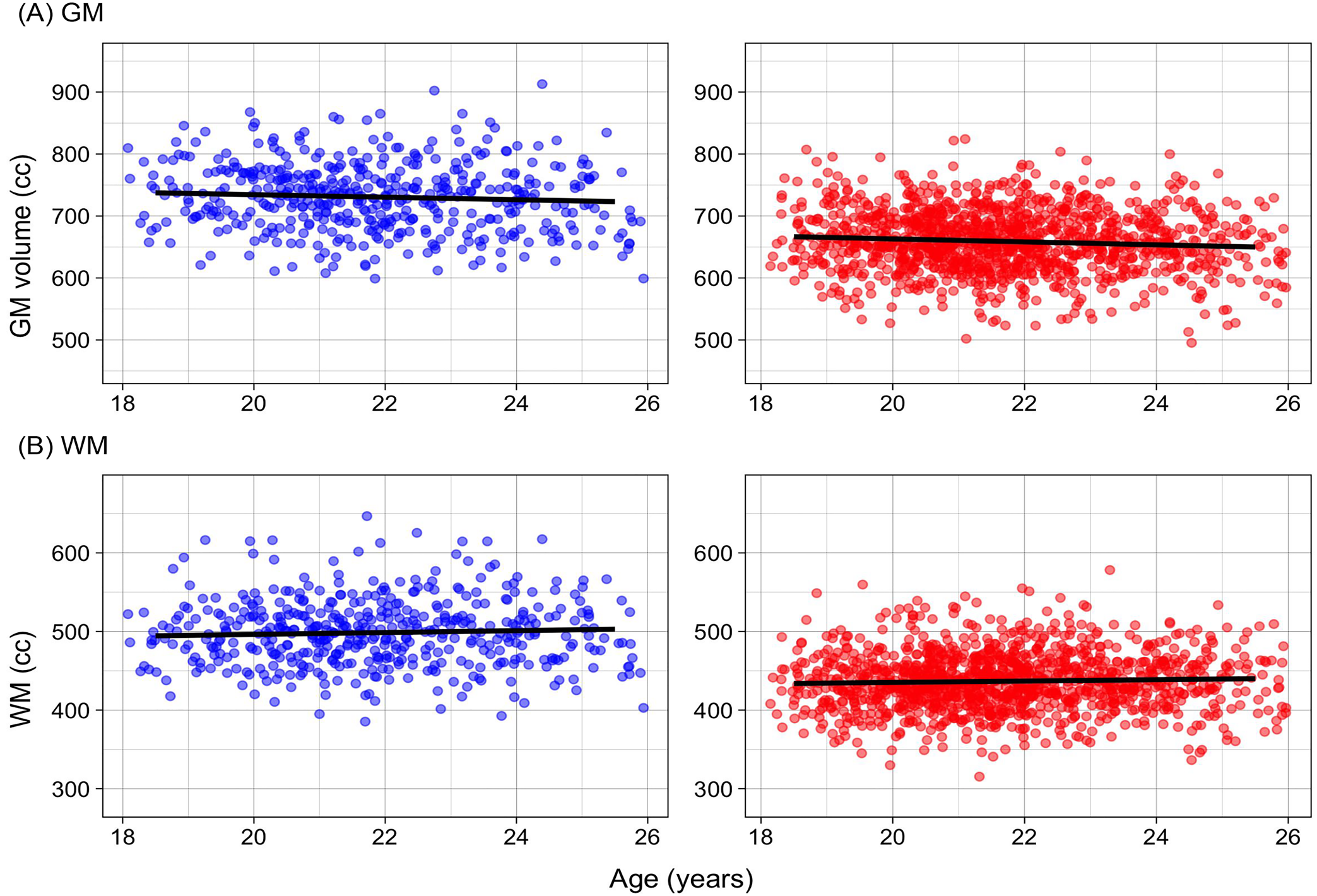
Age-related variations in gray and white matter volumes. (A) The total GM volume, and (B) total WM volume are plotted against age, with individual subjects represented as scatter points (left panel: male, middle panel: female). For each sex, predicted age trajectories, based on the model selected by BIC (see text) in the sex-specific analyses are shown. The plots in the right panel show predicted trajectories for hypothetical male or female subjects with identical, global mean eTIV, based on combined group analyses. The 95% confidence intervals of the predictions are represented as shades around the line.

The best model for the mean CT was the model without eTIV in both sexes, which explained 0.1 and 1.0% of total variance in male and female data, respectively. In the combined group analysis, age, sex, and their interaction together explained 2.5% of the variance. There was a significant main effect of sex, with males showing slightly thicker CT than females (approximately 0.8% difference in CT).

A significant decrease in mean CT with age was observed in females and in the combined group. In males, mean CT also decreased with age but not significantly, but there was no significant interaction between the age and sex on mean CT (Table 3, Figure 5A and Supplementary Table 1). Strictly speaking, the observed age effects are age-associated variations in this cross-sectional cohort, but assuming they represent the age-related changes at this age range, the observed change represents annual percent change (APC) of roughly −0.16% in females and −0.12% in the combined group.

Contrary to its absence of effect on mean CT, eTIV significantly impacted the other GM morphometrics (inner CSA, pial CSA, and GM volume, all *p*’s < 2 x 10^−16^) and explained a large amount of the variances of these data (over 76% for all, based on partial *η^2^* values for eTIV in combined group analyses). None of these metrics showed any significant interaction between sex and eTIV (all *p*’s > 0.36), indicating similar effects of eTIV on these measures across sexes. Thus, below we report age and sex effects for these metrics based on the model including eTIV (model (2) for sex-specific analyses and (4) for combined analyses).

Proportions of age-related variance in inner CSA were similar to those of CT, explaining only 1.1, 0.2, and 0.4% in males, females, and combined groups, respectively. In contrast to CT, the age effect on inner CSA was significant in males (*p* = 0.026) and in the combined group (*p* = 0.0029) but did not reach significance in females (*p* = 0.093).

As for CT, the estimated effects were consistently negative, representing an APC of about −0.2% in males and the combined group, and there was no detectable difference in the age trajectory between males and females (*p* = 0.21). Although raw values of inner CSA were larger in males compared to females, the difference was actually slightly but significantly reversed in the model with eTIV (0.7% larger in females than in males, *p* = 0.008).

Unsurprisingly, inner CSA and pial CSA were highly correlated in our dataset (pearson correlation *r* = 0.986, Supplemental Figure 16), and the total variance of pial CSA data explained by the models with eTIV was similar to those of inner CSA (over 74% in all cases). Yet, observed age-related variances for pial CSA were consistently higher than for inner CSA or CT data (2.7, 1.2, and 1.6% in males, females, and combined data). There were significant age effects in both sex-specific and combined analyses, with the estimated effects representing APC of about −0.2 to −0.3%. In the analysis that combined both sexes, no interaction between sex and age was detected. Similar to what was observed for inner CSA, the pial CSA was larger for males compared to females, but the difference was reversed non-significantly when eTIV was included in the model (0.2% larger in females than in males; *p* = 0.33; Table 3 and Supplemental Table 1).

The total variance of GM volume explained by age, eTIV, and sex in the case of sex-combined analysis was slightly higher than for CSA data (over 78% in all cases), and the age-related variance in GM volume tended to be higher than CT or CSA data individually (2.3, 3.2, and 2.9% in male, female, and combined data). The total GM volume significantly decreased with age in all comparisons (*p*’s < 9.0×10^−4^), representing an APC of approximately −0.3%. Again there was no detectable difference in age-related reduction between males and females (*p* = 0.61). Not surprisingly, males had larger total GM than females when not taking into account the overall head size. When eTIV was taken into account, females showed about 1% larger volume than males (*p* = 1.0×10^−4^; Table 3).

### Global white matter morphometry and DTI parameters

Table 4 provides the descriptive statistics of the following metrics of white matter: total WM volume and mean values of DTI metrics. The table also contains a summary of age and sex effects on these metrics, which are also visually presented in Figure 6B for WM volume, and Figure 7 for mean DTI metrics.

**Table 4.**
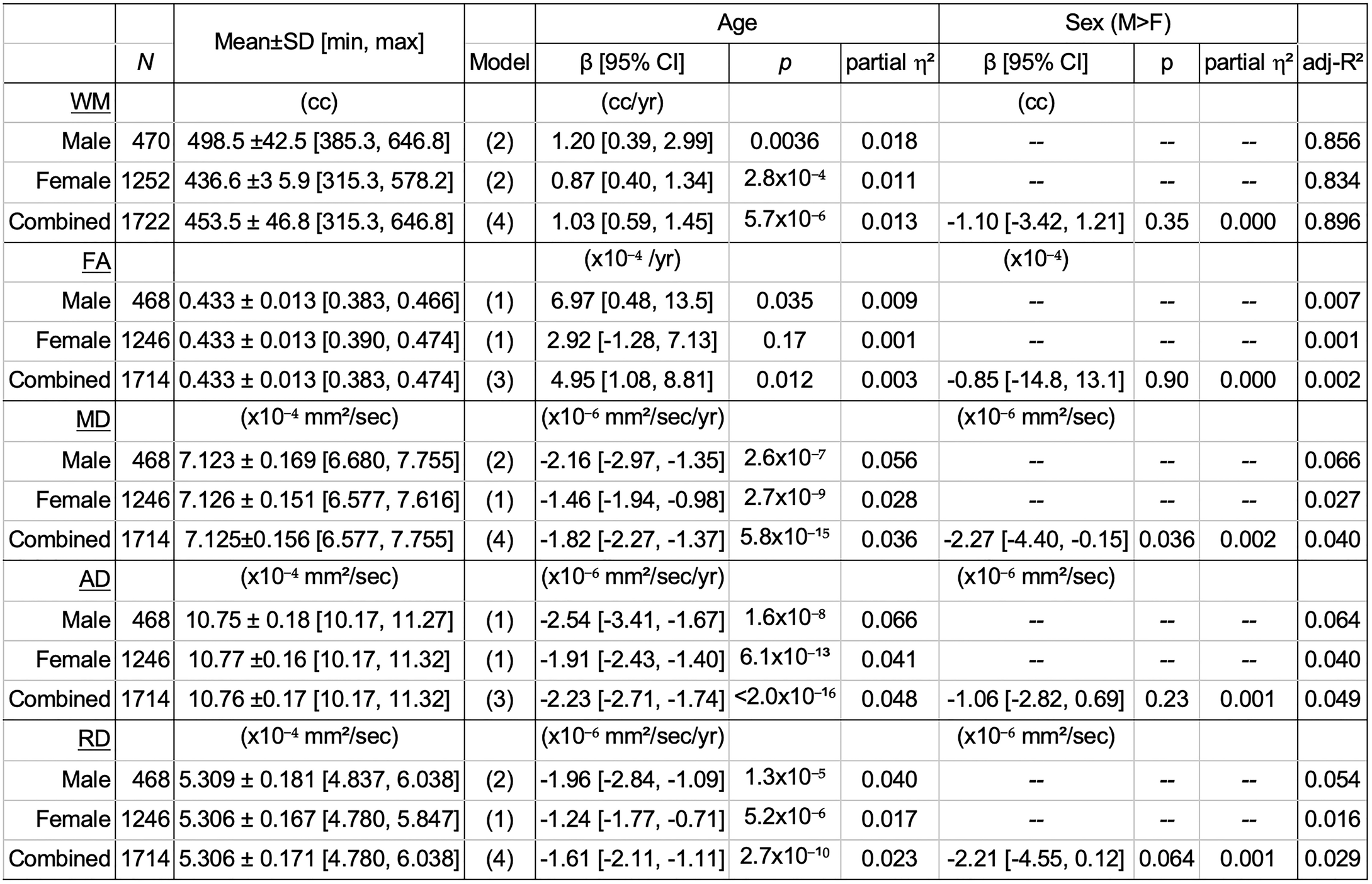
Summary of white matter volume and mean DTI metrics within cerebral white matter. Descriptive statistics and the results of linear model fit for total white matter volume (WM) from SPM12, as well as mean DTI metrics within subject-specific cerebral WM masks are shown. Parameter estimates (β) for age and sex are shown, together with 95% confidence intervals (CI) for each β, as well as uncorrected p values and partial eta squared as the effect sizes for each variable. The Model column indicates the chosen model (see text for details on the model selection), with the corresponding model number as described in Methods section. The interaction between sex and age was tested in the combined group but was not significant in any of the metrics listed here, and thus is not included in the table. The variance explained by eTIV or mask volume for models that included them is discussed in the text. Overall model fit is indicated as adjusted squared R values.

**Figure 7.**
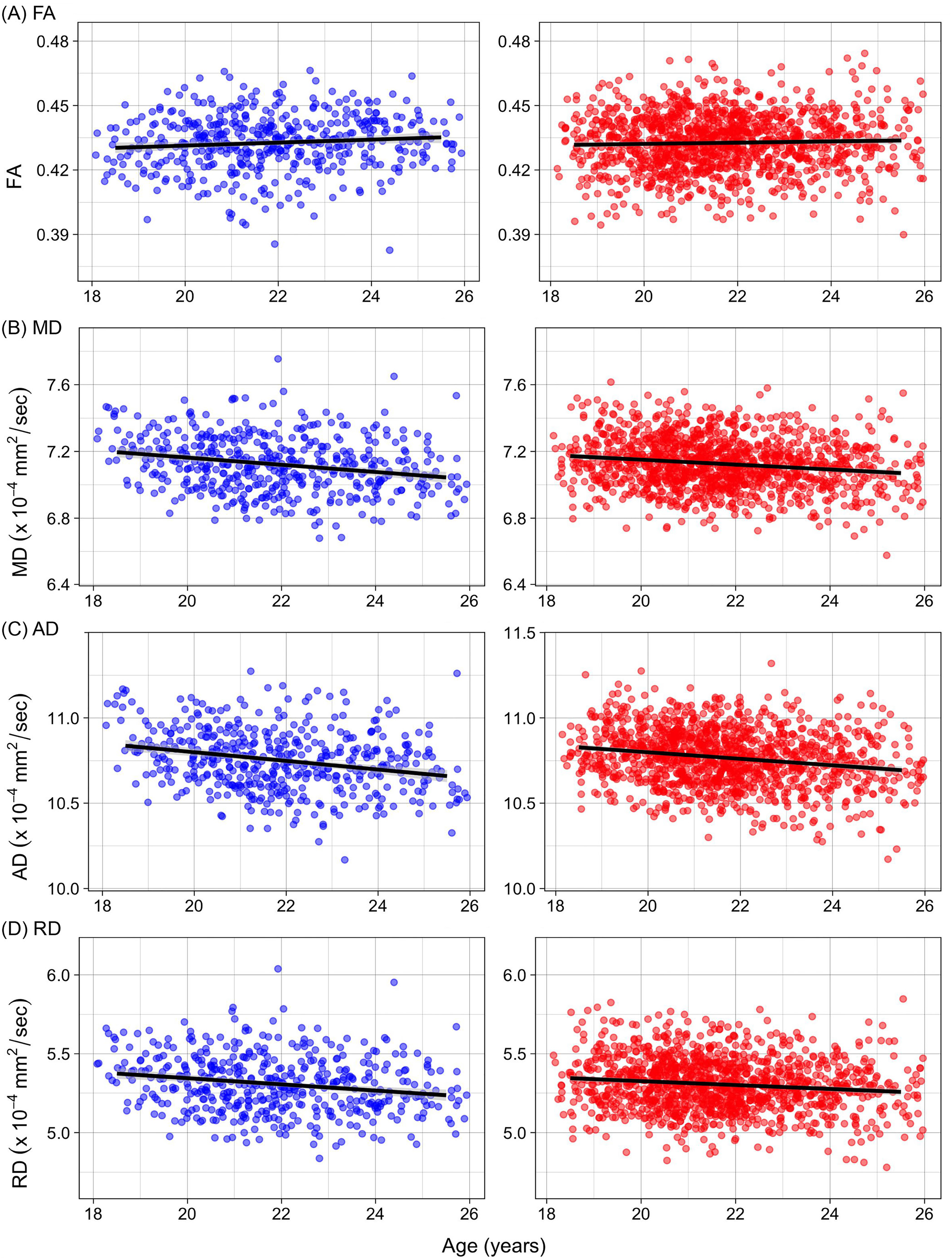
Age-related variations in mean DTI values within the cerebral WM mask. The mean values of (A) FA, (B) MD, (C) AD, and (D) RD within subject-specific cerebral WM mask are plotted against age, with individual subjects represented as scatter points (left panel: male, middle panel: female). For each sex, predicted age trajectories, based on the model selected by BIC (see text) in the sex-specific analyses are shown. The plots in the right panel show predicted trajectories for hypothetical male or female subjects with identical, global mean cerebral WM volume, based on combined group analyses. The 95% confidence intervals of the predictions are represented as shades around the line.

Like for CSA and GM volume data, eTIV explained a large portion of variance in the WM volume data (over 83% in all cases). Of note, unlike in GM morphometric data, there was a significant interaction between sex and eTIV (*p* = 4.1×10^−4^), with males showing steeper slope than females for the effect of eTIV on WM volume. There was a positive and significant linear effect of age on the WM volume in both sexes (Table 4, Figure 6B and Supplemental Table 2), with an APC of about 0.2%. There was no detectable sex differences on age trajectory, and age accounted for 1 to 2% of total variance in the WM volume data. As in GM morphometric data, the sex difference in WM volume was non-significant with the inclusion of eTIV in the model (*p* = 0.35).

Like for CSA and GM volume data, eTIV explained a large portion of variance in the WM volume data (over 83% in all cases). Of note, unlike in GM morphometric data, there was a significant interaction between sex and eTIV (*p* = 4.1×10^−4^), with males showing steeper slope than females for the effect of eTIV on WM volume. There was a positive and significant linear effect of age on the WM volume in both sexes (Table 4, Figure 6B and Supplemental Table 2), with an APC of about 0.2%. There was no detectable sex differences on age trajectory, and age accounted for 1 to 2% of total variance in the WM volume data. As in GM morphometric data, the sex difference in WM volume was non-significant with the inclusion of eTIV in the model (*p* = 0.35).

Overall, the cerebral WM mask volume used to compute DTI mean values did not influence the estimate of age effects on these mean values, and the BIC index indicated that the mask volume contributed little to the overall fit in most comparisons. The only exceptions were the mean MD and RD values in males, where diffusivity values were positively correlated with the mask volume (with partial *η^2^* of 0.017 and 0.021 for MD and RD, respectively, representing less than 0.01% increase in diffusivity per cc; *p*’s < 0.005). This, however, did not impact the estimates of age effects, and only slightly affected the sex effect estimates in the combined group analyses. Below we report the results of the selected models (i.e. models with mask volume for MD and RD in males and in combined group, and models without mask volume for the rest), but provide the alternative model results in the Supplemental Table 2.

All the diffusivity measures (MD, AD, and RD) showed robust significant decrease with APC of about −0.2% for AD and −0.3% for RD, with MD somewhere in between over the age range of our dataset (all *p*’s ≤ 1.3×10^−5^; Table 4, Figure 7). FA, by contrast, only showed a trend for increase at this age range that reached significance in the males (*p* = 0.035) and sex-combined group (*p* = 0.012), but not in females (*p* = 0.17). Age accounted for 4.0 to 6.6% of the total variance of the diffusivity metrics (most in AD, least in RD) in males, and 1.7 to 4.1% in females, and about 1% or less in mean FA values for both sexes. The combined group analyses did not show any evidence for a sex difference in the age-related changes in any of the DTI metrics. There was a trend for a main effect of sex only for MD (greater diffusivity in females than in males at mean age), which reached uncorrected *p* < 0.05, but only in the model with the mask volume (*p* = 0.036; Table 4 and Supplemental Table 2). Of note, AD also showed the similar trend for the main effect of sex in the model with the mask volume, which was not selected based on BIC (Supplemental Table 2).

Table 5 and Figure 8 summarize the results of similar analyses conducted on mean NODDI metrics over the same subject-specific cerebral WM mask used to compute mean DTI metrics. Similarly to the DTI metrics, the effects of mask volume on the overall model fit as well as on observed age and sex effects were tested. The mask volume did not impact the estimates of age effects on NODDI metrics, but did show a small but highly significant relationship with IsoVF, with larger mask volume associated with higher IsoVF (0.03 to 0.04% increase per cc mask volume with *p*’s < 2 x 10^−8^ in both sexes, explaining 6.6 and 3.3% of total variance for males and females, respectively). It modified the magnitude of sex effect on mean IsoVF, but did not significantly impact that of mean NDI or ODI values (Table 5 and Supplemental Table 3). Below we present the results of models without mask volume for NDI and ODI, and the model with mask volume for IsoVF, but the alternative model results can be found in the Supplemental Table 3.

**Table 5.**
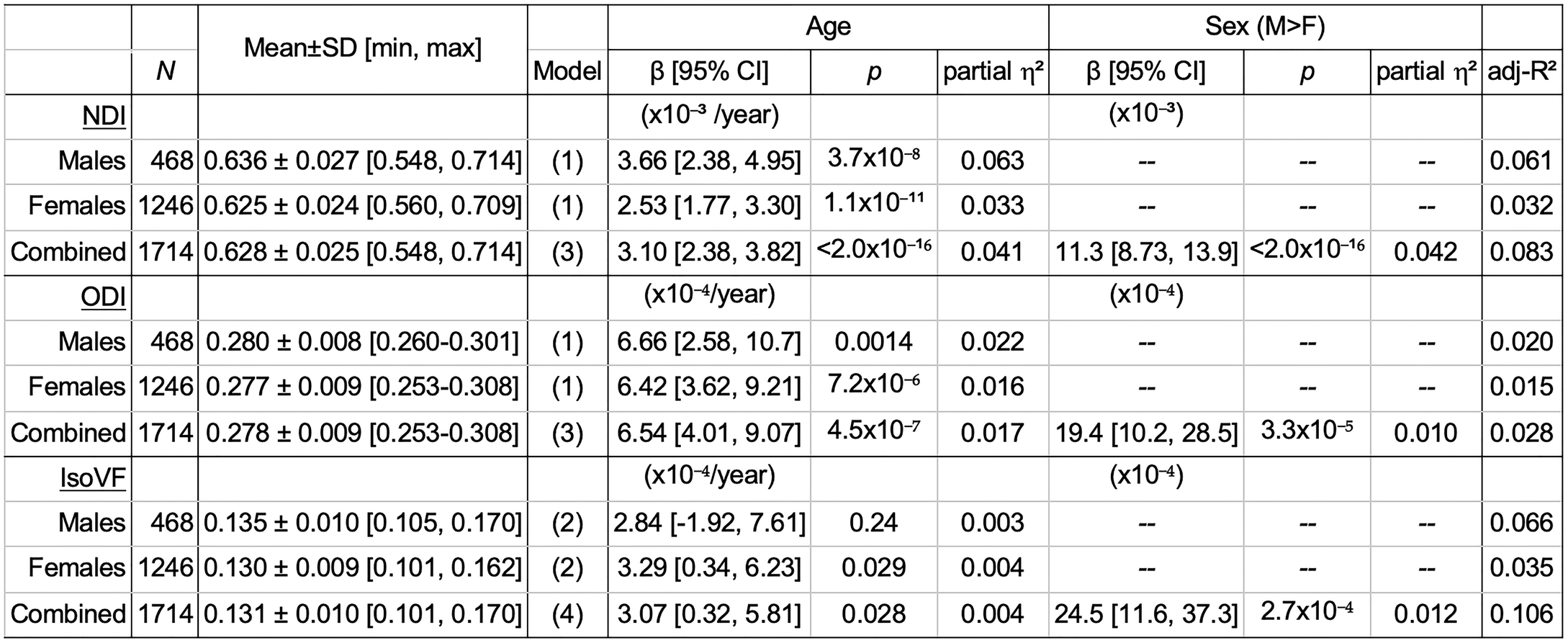
Summary of mean NODDI metrics within cerebral white matter. Descriptive statistics and the results of linear model fit for mean NODDI metrics within subject-specific cerebral WM masks are shown. Parameter estimates (β) for age and sex are shown, together with 95% confidence intervals (CI) for each β, as well as uncorrected p values and partial eta squared as the effect sizes for each variable. The Model column indicates the selected model (see text) with the corresponding model number as described in Methods section. The interaction between sex and age was tested in the combined group but was not significant in any of the metrics listed here, and thus is not included in the table. The variance explained by mask volume for models that included it is discussed in the text. Overall model fit is indicated as adjusted squared R values.

| | | Mean±SD [min, max] | | Age | | | Sex (M>F) | | | |
| --- | --- | --- | --- | --- | --- | --- | --- | --- | --- | --- |
| | $N$ | | Model | $\beta$ [95% CI] | $p$ | partial $\eta^2$ | $\beta$ [95% CI] | $p$ | partial $\eta^2$ | adj-$R^2$ |
| **NDI** | | | | ($\times 10^{-3}$ /year) | | | ($\times 10^{-3}$) | | | |
| Males | 468 | 0.636 ± 0.027 [0.548, 0.714] | (1) | 3.66 [2.38, 4.95] | 3.7$\times 10^{-8}$ | 0.063 | -- | -- | -- | 0.061 |
| Females | 1246 | 0.625 ± 0.024 [0.560, 0.709] | (1) | 2.53 [1.77, 3.30] | 1.1$\times 10^{-11}$ | 0.033 | -- | -- | -- | 0.032 |
| Combined | 1714 | 0.628 ± 0.025 [0.548, 0.714] | (3) | 3.10 [2.38, 3.82] | <2.0$\times 10^{-16}$ | 0.041 | 11.3 [8.73, 13.9] | <2.0$\times 10^{-16}$ | 0.042 | 0.083 |
| **ODI** | | | | ($\times 10^{-4}$/year) | | | ($\times 10^{-4}$) | | | |
| Males | 468 | 0.280 ± 0.008 [0.260-0.301] | (1) | 6.66 [2.58, 10.7] | 0.0014 | 0.022 | -- | -- | -- | 0.020 |
| Females | 1246 | 0.277 ± 0.009 [0.253-0.308] | (1) | 6.42 [3.62, 9.21] | 7.2$\times 10^{-6}$ | 0.016 | -- | -- | -- | 0.015 |
| Combined | 1714 | 0.278 ± 0.009 [0.253-0.308] | (3) | 6.54 [4.01, 9.07] | 4.5$\times 10^{-7}$ | 0.017 | 19.4 [10.2, 28.5] | 3.3$\times 10^{-5}$ | 0.010 | 0.028 |
| **IsoVF** | | | | ($\times 10^{-4}$/year) | | | ($\times 10^{-4}$) | | | |
| Males | 468 | 0.135 ± 0.010 [0.105, 0.170] | (2) | 2.84 [-1.92, 7.61] | 0.24 | 0.003 | -- | -- | -- | 0.066 |
| Females | 1246 | 0.130 ± 0.009 [0.101, 0.162] | (2) | 3.29 [0.34, 6.23] | 0.029 | 0.004 | -- | -- | -- | 0.035 |
| Combined | 1714 | 0.131 ± 0.010 [0.101, 0.170] | (4) | 3.07 [0.32, 5.81] | 0.028 | 0.004 | 24.5 [11.6, 37.3] | 2.7$\times 10^{-4}$ | 0.012 | 0.106 |

**Figure 8.**
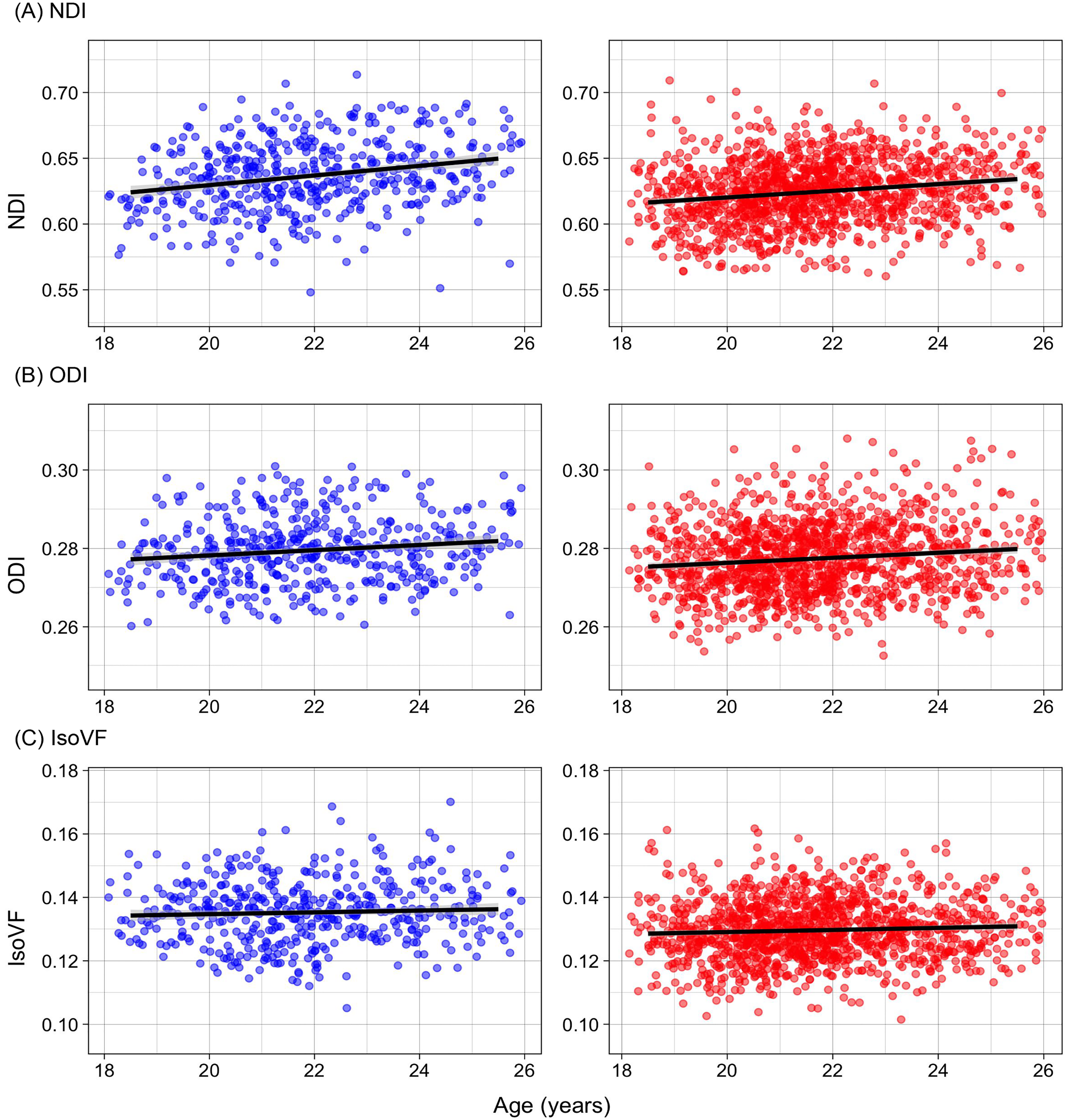
Age-related variations in mean NODDI values within the cerebral WM mask. The mean values of (A) NDI, (B) ODI, and (C) IsoVF within subject-specific cerebral WM mask are plotted against age, with individual subjects represented as scatter points (left panel: male, middle panel: female). For each sex, predicted age trajectories, based on the model selected by BIC (see text) in the sex-specific analyses are shown. The plots in the right panel show predicted trajectories for hypothetical male or female subjects with identical, global mean cerebral WM volume, based on combined group analyses. The 95% confidence intervals of the predictions are represented as shades around the line.

Both NDI and ODI showed a small but robust increase with age at this age range in both the male and female groups with APC of about 0.6% and 0.2% for NDI and ODI in males, and 0.4% and 0.2%, in females. The age effect on IsoVF only reached the uncorrected *p* < 0.05 in females (*p* = 0.029) but not in males (*p* = 0.24), in the positive direction (APC of about 0.25%). The combined group analyses indicated robust main effects of sex for all the NODDI metrics, with males showing higher values for all three metrics, representing 1.8, 0.7, and 1.9% larger values in males relative to femalesfor NDI, ODI, and IsoVF, respectively. However, none of the NODDI metrics showed evidence for the sex difference in the age-related changes. Partial *η^2^* indicated that the amount of variance explained by age for NODDI metrics in the combined group to be about 4, 2, and 0.4% for NDI, ODI, and IsoVF, respectively, while main effect of sex accounted for about 4, 1, and 1% of the variance.

### Resting-state fMRI

Figure 9A shows the average regional IC matrix computed on the 384 regions of the AICHA homotopic functional atlas. As the definition of the region is based on the functional signal, some regions (such as the rolandic region) cross the anatomical borders because they exhibit very high correlations. The regional IC matrices computed with and without global signal regression are shown side by side. Figure 9B, C, and D show the group average maps of ReHo, ALFF, and fALFF, respectively.

**Figure 9.**
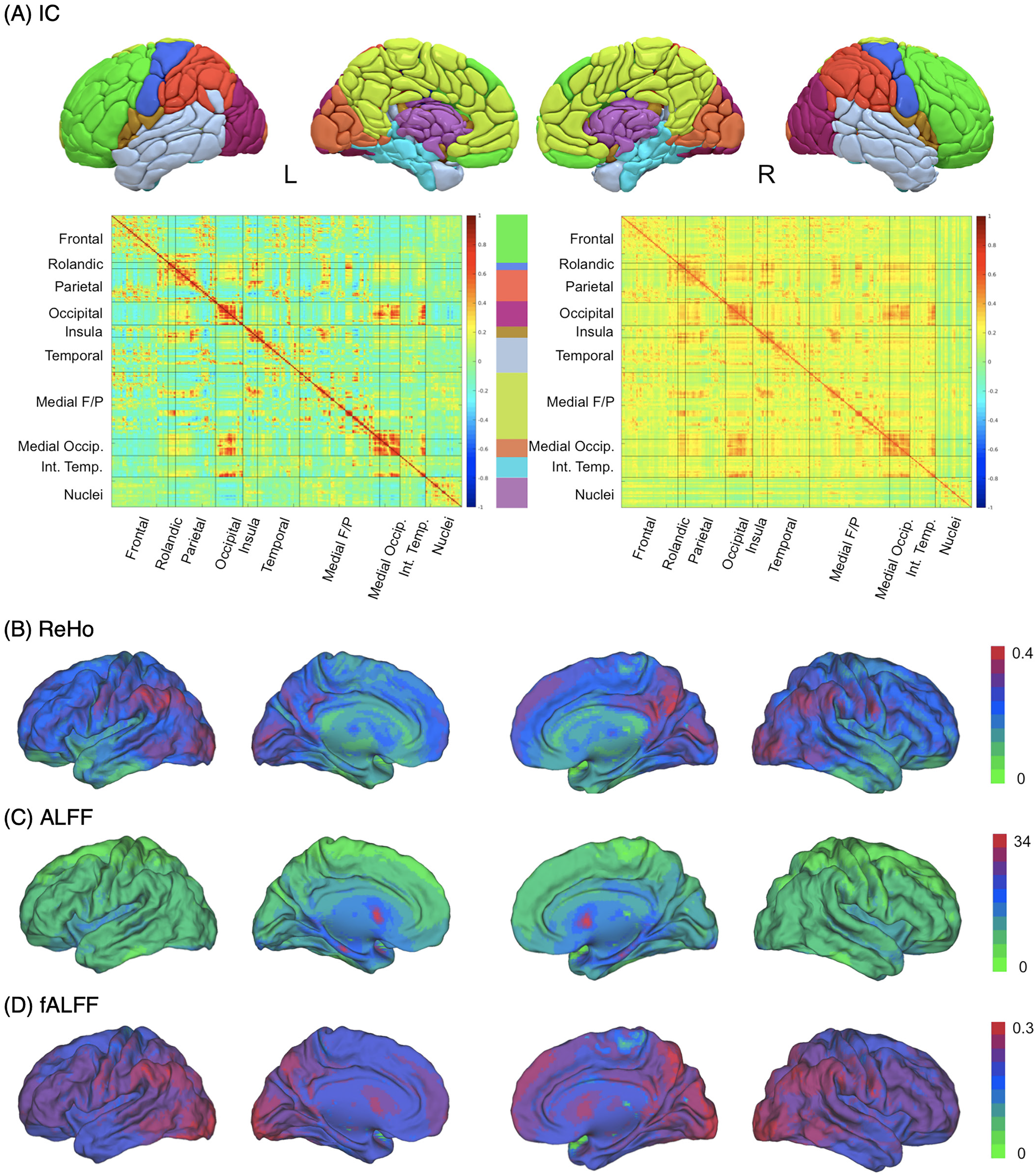
Group average IC matrices and three rs-fMRI-based metrics in 1,814 MRi-Share subjects. (A) AICHA atlas-based regional IC matrices computed from rs-fMRI signals with (left) and without (right) global signal regression. The top panel shows the 10 anatomical partitions of 384 AICHA regions, with the color corresponding to the color block between the two IC matrices (Medial F/P: medial fronto-parietal; Medial Occip.: medial occipital; Int.Temp.: internal temporal), displayed with Surf Ice (https://www.nitrc.org/projects/surfice). Voxel-level group average maps of (B) ReHo, (C) ALFF, and (D) fALFF in the spatially normalized space are superimposed on fiducial surface of PALS template using Caret 5.65 (http:brainvis.wustle.edu).

## Summary of the Results

In a sample of students aged between 18 and 26 years, several significant age-associated variations were observed. Cortical GM thickness, surface area, and total GM volume were decreasing with age, whereas WM total volume showed increase. Diffusivity, either radial or axial (and therefore mean) was found to robustly decrease with age but fractional anisotropy showed only a tendency for a slight increase. Both NDI and ODI robustly increased with age during this period, accompanied with a trend for increasing IsoVF as well. There were no sex differences in age-related trajectory in any of the global metrics we reported here. For GM morphometry, we observed thicker CT in males than in females, while inner CSA and GM volume were larger in females than in males when eTIV was taken into account. The WM DTI metrics did not differ between the two sexes. By contrast, all three NODDI metrics showed a strong sex effects, with higher values in males than in females.

## Discussion

We will first address the impact that some specific features of the MRi-Share study design may have on the interpretation of these findings. We then discuss our results with respect to the existing literature on the maturational/ageing changes as well as sex differences in global brain phenotypes. Lastly, we will describe the future directions and perspectives on the MRi-Share database specifically, and on the even richer resources available when combining this database with the Bio-Share and i-Share study data.

### Specific features of the MRi-Share study design

The MRi-Share sample of about 2,000 students, which has been drawn from the i-Share participants from Bordeaux University, is not a representative sample of the population of healthy young adults and was actually never intended to be so. It must indeed be reminded that the primary objective of the i-Share cohort study is to investigate the student population health. As a consequence, our study sample concerns only 25% of the population of the same age range, and its structure differs from the 75% others particularly in terms of socio-demographics, lifestyle, and level of education. Some of these sampling biases can be accounted for through stratification as we did for example for sex. Some others, such as level of education or lifestyle for example, will require access to additional sources of data to be accounted for.

Another source of concern is the non-uniformity of the sample age distribution as the number of students reduces with the number of years attended at the University. This led us to analyse only a subsample of the participants with a shorter age range (18 to 26 years), thereby avoiding claiming findings to be valid for those 110 participants aged between 26 to 35 years. Additional data are clearly needed to study how the brain changes during this later period of early adulthood. Several potential confounding factors in interpreting our preliminary were not accounted for, including past or current history of mental illness, alcohol intake, smoking habits, and/or use of any recreational drugs and psychotropic medications, … Effects of these factors on structural/functional brain phenotypes in the MRi-Share cohort papers will be reported in other forthcoming publications.

The MRi-Share study design is also cross-sectional in nature, which makes one take the results of the present study with caution. Numerous reports have indeed pointed out the caveats of cross-sectional design for assessing effects of age and demonstrated how such design may lead to spurious findings when compared to those obtained with longitudinal data (Fjell et al., 2010; Pfefferbaum and Sullivan, 2015). In the present work, we tried to minimize these pitfalls by reducing the age range to 8 years only and by selecting a simple linear model to examine age-related changes. We note that several results of the present study are indeed compatible with those of previously published longitudinal studies (see discussion below) in adolescents and adults. However, such precautions do not fully eradicate some intrinsic limits of our cross-sectional study, and particularly the impossibility of identifying the dynamics of age trajectories for the different brain phenotypes as has been done in other studies (Raznahan et al. 2011). Such limitations could be alleviated by having MRi-Share participants undergo a follow-up MRI examination.

### Age-related patterns of cortical thickness, surface area’ and GM volume

We observed varying degrees of age-related variations in all GM morphometric measures we examined in the students aged between 18 and 26 years old. All metrics showed apparent reduction with age, suggesting that CT and both inner and pial CSA are shrinking, resulting in the reduction of total GM volume, even at the narrow age-range of late adolescence and young adulthood represented by the MRi-Share cohort. We did not find any sex differences in the expected age-related trajectory of these metrics. The observed age-related trajectories are in line with the large body of work documenting the age-related variations and changes in brain structural morphometry over the lifespan, which has revealed that while both GM and WM volumes increase rapidly during infancy, GM volume starts to decrease during childhood, when WM volume continues to grow well into adulthood, before declining at middle-to late-adulthood (reviewed in Fjell and Walhovd, 2010; Lebel and Deoni, 2018; Vijayakumar et al., 2018). The GM volume reduction is accompanied by the reduction in both CT and CSA post-childhood, although the age-related trajectories of CT and CSA are largely independent (Potvin et al., 2017; Wierenga et al., 2014), and regionally the reduction in CT and CSA may even have negative relationships (Hogstrom et al., 2013; Storsve et al., 2014).

The estimates of APC observed in our cross-sectional data (CT: −0.12%, inner CSA: −0.16%, GM: −0.32%, based on the combined-sex analyses) were less than those observed in longitudinal cohorts during development (CT: −0.8 to −1.4%, inner CSA: −0.4 to −0.7%, cortical GM: −1.1 to −1.9% between 7 to 29 years of age, Tamnes et al., 2017), and very close to those observed during adult life (CT: −0.16, −0.35%, inner CSA: −0.22, −0.19%, cortical GM: −0.41, −0.51% reported in cohorts between 18 to 87 years of age in Lemaitre et al. 2012, and between 23 to 87 years of age in Storsve et al. 2014, respectively). The slightly lower estimate for GM volume APC in our data may be partially explained by the fact that we report total GM volume, rather than cortical GM volume only, which has been shown to decrease more with age compared to subcortical GM volumes (Walhovd et al., 2011, 2005). Alternatively, the lower GM volume, as well as the slightly lower CT and CSA APC may reflect the slowed volume change at the end of maturation and before the onset or the early phase of the ageing-related reduction in volume. We also demonstrated that pial CSA is more sensitive to age-related changes (estimated APC in the combined group at −0.27%) than both the inner CSA computed based on GM/WM boundary and the CT. This was despite the tight correlation between the inner and pial CSA. It may be explained by the notion that the pial surface is likely to be affected by age-related variations in CT more than in the white surface (Winkler et al., 2012), resulting in the slightly greater age-related reduction than either inner CSA or CT.

### Age-related patterns of global WM volume and DTI/NODDI metrics

In contrast with age effects on GM morphometric measures, the eTIV-adjusted WM volume was found to be increasing between the ages of 18 and 26 years, at the rate of 0.23% annually, whereas the raw WM volume was only slightly but not significantly affected by age positively (APC = 0.08%, Supplementary Table 2). These findings are consistent with the literature that indicates the inverted U-shape age trajectory of WM volume, which increases largely linearly up to the fourth or fifth decade of life, before declining in later life (Hasan et al., 2007; Lebel et al., 2012; Walhovd et al., 2011; Westlye et al., 2010). The estimates of APC depend on the precise age range examined as well as the reference age, which are not always reported in the literature, as well as on whether or not adjustment for intracranial volume was performed. For instance, Walhovd et al. (2011) reported percent change per decade for cerebral WM to be 3.9%, calculated based on mean raw WM volumes of those aged 18 to 29 years and those aged 30 to 39 years, which would mean roughly 0.4% APC, assuming a linear and constant increase during the decade. Although the percent change per decade in this study was based on raw WM volume not corrected for intracranial volume (ICV), another study investigating multiple younger cohorts reported APC (estimated at each age) of 0 to 1% for the age range of 20 to 30 years when correcting for ICV either by dividing the WM by the ICV or by including ICV as a covariate in the model, similar to what we did here (Figure S4 in Mills et al., 2016), more in line with our estimated APC.

The observed age-related variations of WM microstructure IDPs are in consistent with the past studies that revealed the U-shaped trajectories for WM water diffusivity measures (MD, AD, and RD), which all keep decreasing throughout development and reach their minimum values between second and fourth decade of life, before increasing with age, and FA showing the opposite pattern of trajectory that increases during development and decreases in ageing (Beaudet et al., 2020; Hasan et al., 2010, 2007; Lebel et al., 2012; Slater et al., 2019; Westlye et al., 2010). More limited number of studies using NODDI showed the continuous increase of NDI through development (Genc et al., 2017; Mah et al., 2017) and adulthood (Chang et al., 2015; but see Kodiweera et al., 2016), peaking around the fourth and fifth decade of life (Slater et al., 2019) before declining at a later age (Cox et al., 2016; Merluzzi et al., 2016). ODI, on the other hand, has not been reported to change noticeably during development but starts to increase during young adulthood (Chang et al., 2015; Genc et al., 2017; Mah et al., 2017), peaking between the fourth and sixth decade of life and declining thereafter (Cox et al., 2016; Slater et al., 2019).

Although both DTI and NODDI metrics are often described in terms of biological properties, in reality they are based on mathematical models that are fit to explain DWI data that are collected at the resolution many times coarser than the structures these models aim to probe (axonal fibers, intracellular and extracellular compartments). Thus, it is not always straightforward to interpret apparent changes or differences in these metrics, in particular for DTI metrics, which could be affected by many different microstructural features (number, size, and orientations of axonal fibers, membrane permeability, myelination etc) (Jones et al., 2013). Nonetheless, it could be speculated that the apparent age-related elevation in NDI and ODI is consistent with the interpretation that continuing growth of neurites that steadily increase intracellular (axonal) diffusion in the cerebral WM is coupled with increased overall fiber complexity at this age range (ref needed). Such increase in overall fiber complexity, coupled with reduced extra-axon diffusion as a result of denser packing of axons (Suzuki et al., 2003), can account for the overall reduction in the rate of diffusion in the simple tensor model estimated at the voxel-level.

### Sex effects on GM and WM global phenotypes

Beyond the well-known difference in the overall head and brain size, the findings with regard to sex differences in global and regional GM morphometry and in age-related trajectory on of these morphometric measures are often mixed (Fjell et al., 2015; e.g. Gennatas et al., 2017; Hasan et al., 2007; Herron et al., 2015; Jahanshad and Thompson, 2017; Kaczkurkin et al., 2019; Lemaître et al., 2005; McKay et al., 2014; Mutlu et al., 2013; Raznahan et al., 2010; Ritchie et al., 2018; Sowell et al., 2007), and are known to depend on if and how the effect of head or brain size is taken into account (Herron et al., 2015; Mills et al., 2016). Generally, CSA and GM volumes, but not CT, are strongly correlated with the overall brain size (Potvin et al., 2017), and unsurprisingly, males show larger absolute values in these metrics compared to females, as was the case in our data. When corrected for eTIV, we saw a slight but significantly greater GM volume in females than in males and a similar trend for CSA in our data, although the difference was very small relative to the total variance of data (sex effects accounting for less than 1% of variance for both, based on partial *η^2^* values). For CT, prior large-scale studies (sample size > 500) have reported either no sex difference in global mean CT during development and over lifespan (Ducharme et al., 2016; Fjell et al., 2015; Gennatas et al., 2017), or globally greater CT in females than in males in mid-to late-adulthood (Ritchie et al., 2018), though there are also some reports of greater mean CT in males than in females throughout development (Raznahan et al., 2011) and over lifespan (Kochunov et al., 2011). We found significantly thicker mean CT in males than in females in our data, which was not affected by eTIV correction, that accounted for nearly 2% of variance in the CT data. Since sex differences in CT are also reported to be regionally heterogeneous and age-dependent (Mutlu et al., 2013; Raznahan et al., 2010; Sotiras et al., 2017; Sowell et al., 2007), future investigations will examine the regional patterns in the CT difference and clarify whether what we observe is consistent with what would be expected at this age range or somehow unique to our sample due to some specific aspects of demographic characteristics (e.g. higher education) in our data.

We did not find any sex differences in eTIV-adjusted WM volume or its age-related variations, consistent with prior studies reporting similar normalized WM volumes and age-related trajectory during much of the adulthood (Coupé et al., 2017; Fjell et al., 2009; Lebel et al., 2012), although there are a few high-powered studies reporting larger WM volume in males than in females even after correcting for the total brain volume, albeit with much reduced effect sizes (Ritchie et al., 2018; Wierenga et al., 2014).

As for the sex differences in the age trajectories of DTI metrics, most studies report no or minimal differences (Hasan et al., 2010; Hsu et al., 2010, 2008; Lebel et al., 2012; Pohl et al., 2016; Wang et al., 2012), except earlier in development (Simmonds et al., 2014). In our recent multi-cohort study that investigated mean DTI metrics across the WM skeleton in a large sample (total N >20,000) that included the MRi-Share and 9 other cohorts that collectively spanned the entire adult lifespan, we also failed to detect any sex effects on the age-associated trajectory of the DTI metrics at this or any other age ranges examined during adulthood (Beaudet et al., 2020). In the same study, we also found significantly greater AD and MD in females than males when controlling for eTIV, which affected the mean AD, MD, and FA values significantly. While it has been reported that volume over which mean DTI metrics are computed can affect mean values due to PVEs (Vos et al., 2011), the effects of tract or mask volumes, or the overall head and brain size when DTI metrics are compared in spatially normalized space, are not often investigated. Further studies are needed to clarify how it would impact the apparent sex differences or lack of them when comparing DTI metrics.

Very little is known about the sex differences in NODDI metrics. The few studies that report sex effects seem to indicate little or no sex differences in the age trajectory (Cox et al., 2016; Slater et al., 2019), but are inconsistent with regard to the main effects of sex on NODDI metrics: one developmental (Mah et al., 2017) and another lifespan (Slater et al., 2019) study reported no main effects of sex, while one study in young to middle-aged adults reported greater NDI for males compared to females in most WM tracts examined as well as greater ODI in some tracts (Kodiweera et al., 2016), and yet another study with high power (*N* >3,500) reported consistently higher ODI in females compared to males across all the WM tracts examined in older adults (Cox et al., 2016). While the direction of the sex differences we observed is similar to those reported by Kodiweera et al. (2016), the magnitude of the difference was much smaller in our study (2.1 and 0.7% for NDI and ODI in our data, compared to about 7% for both NDI and ODI reported by Kodiweera et al. (2016)). Further studies are needed to investigate whether the sex differences in these microstructural measures manifest themselves in specific time window of early adulthood, or whether they may be modulated by other demographic variables.

### Future directions and perspectives

The MRi-Share database represents a unique, multi-modal neuroimaging and cognitive and genetic dataset of a large, cross-sectional cohort of young adults undergoing university-level education. Its design and the sample size will allow detailed characterization of age-related changes in brain structure and function in the post-adolescence period, as well as the investigation of lifestyle and sociodemographic factors that may modulate these changes.

In addition, the MRi-Share database enriches the relatively scarce corpus of data available in young adults, thereby allowing investigations of structural and functional brain changes across the complete adult lifespan. A proof of concept of such added-value of the MRi-Share sample was recently provided through a multi-cohort study across the adult life span of age-related changes of PSMD (Peak width Skeletonized Mean Diffusivity), a novel imaging marker of small vessel disease (Baykara et al., 2016): in this study, thanks to the MRi-Share sample, PSMD was demonstrated to be the only DTI-derived phenotype that increased in the immediate post-adolescence period, indicating that it could serve as an early marker of aging (Beaudet et al., 2020). MRi-Share also provides the opportunity to assess whether imaging markers of late-life disorders can be detected earlier in adult life. With this in mind, we will release in the near future a number of additional IDPs, including white matter hyperintensities (WMH) assessed on T2-FLAIR images, enlarged perivascular spaces (ePVS) assessed on T1w images, and cerebral microbleeds (CMB) assessed on SWI images,all being well-established imaging-markers of small vessel disease (Wardlaw et al., 2013).

## Supporting information

MRi-Share_supplemental

## Acknowledgements

The i-Share cohort has been funded by a grant ANR-10-COHO-05-01 as part of the Programme pour les Investissements d’Avenir). Supplementary funding was received from the Conseil Régional of Nouvelle-Aquitaine, reference 4370420. The MRi-Share cohort and the ABACI software development have been supported by ANR-10-LABX-57 (TRAIL) and ANR-16-LCV2-0006 (GINESISLAB for the software) grants. The bio-Share cohort and some regulatory and ethical aspects of MRi-Share have been supported by the European Research Council (ERC) under the European Union’s Horizon 2020 research and innovation programme under grant agreement No 640643 and the FHU SMART. Acquisition of a MRI scanner dedicated to MRi-Share was made possible thanks to a grant from the Conseil Régional d’Aquitaine. Ami Tsuchida, Naka Beguedou, and Alexandre Laurent have been supported by a grant from the Fondation pour la Recherche Médicale (DIC202161236446), Marie-Fateye Gueye, Violaine Verrecchia, and Victor Nozais by a grant ANR-16-LCV2-0006 (GINESISLAB), and Antonietta Pepe by a grant ANR-15-HBPR-0001-03 (as part of the EU FLAG-ERA MULTI-LATERAL consortium). The authors express their deep gratitude to Loïc Labache for his help in preparing the manuscript. They are also indebted to the following individuals for their invaluable contribution to the MRi-Share project: Serge Anandra, Amandine André, Gregory Beaudet, Christophe Bernard, Bruno Brochet, Aurore Capelli, Claire Cardona, Arnaud Chaussé, Christophe Delalande, Vincent Durand, Louise Knafo, Morgane Lachaize, Hugues Loiseau, Elena Milesi, Marie Mougin, Maylis Melin, Guy Perchey, Clothilde Pollet, Thomas Tourdias, Cécile Marchal, Guillaume Penchet, Cécile Dulau, Igor Sibon, Sabrina Debruxelle, Sophie Auriacombe, Caroline Roussillon, Nicolas Vinuesa, and the i-Share “relay” students. The authors are also indebted to Paul Matthews (Imperial College, London, UK) and to the personnel of the UK Biobank imaging center at Stockport (UK) for their help while designing the Mri-Share image acquisition protocol, and to Maxime Descoteaux (Sherbrook University, Canada) for his help in impementing the DWI processing and QC pipelines.Finally, the authors would like to express their gratitude to the 1,870 students of the Bordeaux University who gave their consent to participate in MRi-Share.

## Data Availability

i-Share is open to collaborations and partnerships and supports local, national, and international collaborations from the public or private sector. Requests to access the database should be sent at.

## Notes

### Competing Interest Statement

The authors have declared no competing interest.

## References

3C Study Group, 2003. Vascular factors and risk of dementia: design of the Three-City Study and baseline characteristics of the study population. Neuroepidemiology 22, 316–325. doi:10.1159/000072920

Alfaro-Almagro, F., Jenkinson, M., Bangerter, N.K., Andersson, J.L.R., Griffanti, L., Douaud, G., Sotiropoulos, S.N., Jbabdi, S., Hernandez-Fernandez, M., Vallee, E., Vidaurre, D., Webster, M., McCarthy, P., Rorden, C., Daducci, A., Alexander, D.C., Zhang, H., Dragonu, I., Matthews, P.M., Miller, K.L., Smith, S.M., 2018. Image processing and Quality Control for the first 10,000 brain imaging datasets from UK Biobank. Neuroimage 166, 400–424. doi:10.1016/j.neuroimage.2017.10.034

Backhausen, L.L., Herting, M.M., Buse, J., Roessner, V., Smolka, M.N., Vetter, N.C., 2016. Quality Control of Structural MRI Images Applied Using FreeSurfer-A Hands-On Workflow to Rate Motion Artifacts. Front. Neurosci. 10, 558. doi:10.3389/fnins.2016.00558

Backhouse, E.V., McHutchison, C.A., Cvoro, V., Shenkin, S.D., Wardlaw, J.M., 2017. Early life risk factors for cerebrovascular disease: A systematic review and meta-analysis. Neurology 88, 976–984. doi:10.1212/WNL.0000000000003687

Basser, P.J., Mattiello, J., LeBihan, D., 1994. MR diffusion tensor spectroscopy and imaging. Biophys. J. 66, 259–267. doi:10.1016/S0006-3495(94)80775-1

Baykara, E., Gesierich, B., Adam, R., Tuladhar, A.M., Biesbroek, J.M., Koek, H.L., Ropele, S., Jouvent, E., Alzheimer’s Disease Neuroimaging Initiative, Chabriat, H., Ertl-Wagner, B., Ewers, M., Schmidt, R., de Leeuw, F.-E., Biessels, G.J., Dichgans, M., Duering, M., 2016. A novel imaging marker for small vessel disease based on skeletonization of white matter tracts and diffusion histograms. Ann. Neurol. 80, 581–592. doi:10.1002/ana.24758

Beaudet, G., Tsuchida, A., Petit, L., Tzourio, C., Caspers, S., Schreiber, J., Pausova, Z., Patel, Y., Paus, T., Schmidt, R., Pirpamer, L., Sachdev, P.S., Brodaty, H., Kochan, N., Trollor, J., Wen, W., Armstrong, N.J., Deary, I.J., Bastin, M.E., Wardlaw, J.M., Mazoyer, B., 2020. Age-Related Changes of Peak Width Skeletonized Mean Diffusivity (PSMD) Across the Adult Lifespan: A Multi-Cohort Study. Front. Psychiatry 11, 342. doi:10.3389/fpsyt.2020.00342

Caspers, S., Moebus, S., Lux, S., Pundt, N., Schütz, H., Mühleisen, T.W., Gras, V., Eickhoff, S.B., Romanzetti, S., Stöcker, T., Stirnberg, R., Kirlangic, M.E., Minnerop, M., Pieperhoff, P., Mödder, U., Das, S., Evans, A.C., Jöckel, K.-H., Erbel, R., Cichon, S., Amunts, K., 2014. Studying variability in human brain aging in a population-based German cohort-rationale and design of 1000BRAINS. Front. Aging Neurosci. 6, 149. doi:10.3389/fnagi.2014.00149

Chang, Y.S., Owen, J.P., Pojman, N.J., Thieu, T., Bukshpun, P., Wakahiro, M.L.J., Berman, J.I., Roberts, T.P.L., Nagarajan, S.S., Sherr, E.H., Mukherjee, P., 2015. White Matter Changes of Neurite Density and Fiber Orientation Dispersion during Human Brain Maturation. PLoS ONE 10, e0123656. doi:10.1371/journal.pone.0123656

Corley, J., Cox, S.R., Deary, I.J., 2018. Healthy cognitive ageing in the Lothian Birth Cohort studies: marginal gains not magic bullet. Psychol. Med. 48, 187–207. doi:10.1017/S0033291717001489

Coupé, P., Catheline, G., Lanuza, E., Manjón, J.V., Alzheimer’s Disease Neuroimaging Initiative, 2017. Towards a unified analysis of brain maturation and aging across the entire lifespan: A MRI analysis. Hum. Brain Mapp. 38, 5501–5518. doi:10.1002/hbm.23743

Cox, R.W., 1996. AFNI: software for analysis and visualization of functional magnetic resonance neuroimages. Comput. Biomed. Res. 29, 162–173. doi:10.1006/cbmr.1996.0014

Cox, S.R., Ritchie, S.J., Tucker-Drob, E.M., Liewald, D.C., Hagenaars, S.P., Davies, G., Wardlaw, J.M., Gale, C.R., Bastin, M.E., Deary, I.J., 2016. Ageing and brain white matter structure in 3,513 UK Biobank participants. Nat. Commun. 7, 13629. doi:10.1038/ncomms13629

Daducci, A., Canales-Rodríguez, E.J., Zhang, H., Dyrby, T.B., Alexander, D.C., Thiran, J.-P., 2015. Accelerated Microstructure Imaging via Convex Optimization (AMICO) from diffusion MRI data. Neuroimage 105, 32–44. doi:10.1016/j.neuroimage.2014.10.026

Deary, I.J., Gow, A.J., Taylor, M.D., Corley, J., Brett, C., Wilson, V., Campbell, H., Whalley, L.J., Visscher, P.M., Porteous, D.J., Starr, J.M., 2007. The Lothian Birth Cohort 1936: a study to examine influences on cognitive ageing from age 11 to age 70 and beyond. BMC Geriatr. 7, 28. doi:10.1186/1471-2318-7-28

Debette, S., Seshadri, S., Beiser, A., Au, R., Himali, J.J., Palumbo, C., Wolf, P.A., DeCarli, C., 2011. Midlife vascular risk factor exposure accelerates structural brain aging and cognitive decline. Neurology 77, 461–468. doi:10.1212/WNL.0b013e318227b227

Desikan, R.S., Ségonne, F., Fischl, B., Quinn, B.T., Dickerson, B.C., Blacker, D., Buckner, R.L., Dale, A.M., Maguire, R.P., Hyman, B.T., Albert, M.S., Killiany, R.J., 2006. An automated labeling system for subdividing the human cerebral cortex on MRI scans into gyral based regions of interest. Neuroimage 31, 968–980. doi:10.1016/j.neuroimage.2006.01.021

Destrieux, C., Fischl, B., Dale, A., Halgren, E., 2010. Automatic parcellation of human cortical gyri and sulci using standard anatomical nomenclature. Neuroimage 53, 1–15. doi:10.1016/j.neuroimage.2010.06.010

Diedrichsen, J., Balsters, J.H., Flavell, J., Cussans, E., Ramnani, N., 2009. A probabilistic MR atlas of the human cerebellum. Neuroimage 46, 39–46. doi:10.1016/j.neuroimage.2009.01.045

Ducharme, S., Albaugh, M.D., Nguyen, T.-V., Hudziak, J.J., Mateos-Pérez, J.M., Labbe, A., Evans, A.C., Karama, S., Brain Development Cooperative Group, 2016. Trajectories of cortical thickness maturation in normal brain development--The importance of quality control procedures. Neuroimage 125, 267–279. doi:10.1016/j.neuroimage.2015.10.010

Dumontheil, I., 2016. Adolescent brain development. Curr. Opin. Behav. Sci. 10, 39–44. doi:10.1016/j.cobeha.2016.04.012

Field, T.S., Doubal, F.N., Johnson, W., Backhouse, E., McHutchison, C., Cox, S., Corley, J., Pattie, A., Gow, A.J., Shenkin, S., Cvoro, V., Morris, Z., Staals, J., Bastin, M., Deary, I.J., Wardlaw, J.M., 2016. Early life characteristics and late life burden of cerebral small vessel disease in the Lothian Birth Cohort 1936. Aging (Albany NY) 8, 2039–2061. doi:10.18632/aging.101043

Fjell, A.M., Grydeland, H., Krogsrud, S.K., Amlien, I., Rohani, D.A., Ferschmann, L., Storsve, A.B., Tamnes, C.K., Sala-Llonch, R., Due-Tønnessen, P., Bjørnerud, A., Sølsnes, A.E., Håberg, A.K., Skranes, J., Bartsch, H., Chen, C.-H., Thompson, W.K., Panizzon, M.S., Kremen, W.S., Dale, A.M., Walhovd, K.B., 2015. Development and aging of cortical thickness correspond to genetic organization patterns. Proc Natl Acad Sci USA 112, 15462–15467. doi:10.1073/pnas.1508831112

Fjell, A.M., Walhovd, K.B., 2010. Structural brain changes in aging: courses, causes and cognitive consequences. Rev. Neurosci. 21, 187–221. doi:10.1515/REVNEURO.2010.21.3.187

Fjell, A.M., Walhovd, K.B., Westlye, L.T., Østby, Y., Tamnes, C.K., Jernigan, T.L., Gamst, A., Dale, A.M., 2010. When does brain aging accelerate? Dangers of quadratic fits in cross-sectional studies. Neuroimage 50, 1376–1383. doi:10.1016/j.neuroimage.2010.01.061

Fjell, A.M., Westlye, L.T., Amlien, I., Espeseth, T., Reinvang, I., Raz, N., Agartz, I., Salat, D.H., Greve, D.N., Fischl, B., Dale, A.M., Walhovd, K.B., 2009. Minute effects of sex on the aging brain: a multisample magnetic resonance imaging study of healthy aging and Alzheimer’s disease. J. Neurosci. 29, 8774–8783. doi:10.1523/JNEUROSCI.0115-09.2009

Fox, J., Friendly, M., Monette, G., 2018. heplots: Visualizing Tests in Multivariate Linear Models. R package version 1.3–5.

Frazier, J.A., Chiu, S., Breeze, J.L., Makris, N., Lange, N., Kennedy, D.N., Herbert, M.R., Bent, E.K., Koneru, V.K., Dieterich, M.E., Hodge, S.M., Rauch, S.L., Grant, P.E., Cohen, B.M., Seidman, L.J., Caviness, V.S., Biederman, J., 2005. Structural brain magnetic resonance imaging of limbic and thalamic volumes in pediatric bipolar disorder. Am. J. Psychiatry 162, 1256–1265. doi:10.1176/appi.ajp.162.7.1256

Garyfallidis, E., Brett, M., Amirbekian, B., Rokem, A., van der Walt, S., Descoteaux, M., Nimmo-Smith, I., Dipy Contributors, 2014. Dipy, a library for the analysis of diffusion MRI data. Front. Neuroinformatics 8, 8. doi:10.3389/fninf.2014.00008

Genc, S., Malpas, C.B., Holland, S.K., Beare, R., Silk, T.J., 2017. Neurite density index is sensitive to age related differences in the developing brain. Neuroimage 148, 373–380. doi:10.1016/j.neuroimage.2017.01.023

Gennatas, E.D., Avants, B.B., Wolf, D.H., Satterthwaite, T.D., Ruparel, K., Ciric, R., Hakonarson, H., Gur, R.E., Gur, R.C., 2017. Age-Related Effects and Sex Differences in Gray Matter Density, Volume, Mass, and Cortical Thickness from Childhood to Young Adulthood. J. Neurosci. 37, 5065–5073. doi:10.1523/JNEUROSCI.3550-16.2017

Gluckman, P.D., Hanson, M.A., Cooper, C., Thornburg, K.L., 2008. Effect of in utero and early-life conditions on adult health and disease. N. Engl. J. Med. 359, 61–73. doi:10.1056/NEJMra0708473

Goldstein, J.M., Seidman, L.J., Makris, N., Ahern, T., O’Brien, L.M., Caviness, V.S., Kennedy, D.N., Faraone, S.V., Tsuang, M.T., 2007. Hypothalamic abnormalities in schizophrenia: sex effects and genetic vulnerability. Biol. Psychiatry 61, 935–945. doi: 10.1016/j.biopsych.2006.06.027

Gorgolewski, K., Burns, C.D., Madison, C., Clark, D., Halchenko, Y.O., Waskom, M.L., Ghosh, S.S., 2011. Nipype: a flexible, lightweight and extensible neuroimaging data processing framework in python. Front. Neuroinformatics 5, 13. doi:10.3389/fninf.2011.00013

Hasan, K.M., Kamali, A., Abid, H., Kramer, L.A., Fletcher, J.M., Ewing-Cobbs, L., 2010. Quantification of the spatiotemporal microstructural organization of the human brain association, projection and commissural pathways across the lifespan using diffusion tensor tractography. Brain Struct. Funct. 214, 361–373. doi:10.1007/s00429-009-0238-0

Hasan, K.M., Sankar, A., Halphen, C., Kramer, L.A., Brandt, M.E., Juranek, J., Cirino, P.T., Fletcher, J.M., Papanicolaou, A.C., Ewing-Cobbs, L., 2007. Development and organization of the human brain tissue compartments across the lifespan using diffusion tensor imaging. Neuroreport 18, 1735–1739. doi: 10.1097/WNR.0b013e3282f0d40c

Herron, T.J., Kang, X., Woods, D.L., 2015. Sex differences in cortical and subcortical human brain anatomy [version 1; peer review: 1 approved, 1 approved with reservations]. F1000Res. 4. doi:10.12688/f1000research.6210.1

Hogstrom, L.J., Westlye, L.T., Walhovd, K.B., Fjell, A.M., 2013. The structure of the cerebral cortex across adult life: age-related patterns of surface area, thickness, and gyrification. Cereb. Cortex 23, 2521–2530. doi:10.1093/cercor/bhs231

Hsu, J.-L., Leemans, A., Bai, C.-H., Lee, C.-H., Tsai, Y.-F., Chiu, H.-C., Chen, W.-H., 2008. Gender differences and age-related white matter changes of the human brain: a diffusion tensor imaging study. Neuroimage 39, 566–577. doi:10.1016/j.neuroimage.2007.09.017

Hsu, J.-L., Van Hecke, W., Bai, C.-H., Lee, C.-H., Tsai, Y.-F., Chiu, H.-C., Jaw, F.-S., Hsu, C.-Y., Leu, J.-G., Chen, W.-H., Leemans, A., 2010. Microstructural white matter changes in normal aging: a diffusion tensor imaging study with higher-order polynomial regression models. Neuroimage 49, 32–43. doi:10.1016/j.neuroimage.2009.08.031

Iglesias, J.E., Augustinack, J.C., Nguyen, K., Player, C. M., Player, A., Wright, M., Roy, N., Frosch, M. P., McKee, A.C., Wald, L.L., Fischl, B., Van Leemput, K., Alzheimer’s Disease Neuroimaging Initiative, 2015a. A computational atlas of the hippocampal formation using ex vivo, ultra-high resolution MRI: Application to adaptive segmentation of in vivo MRI. Neuroimage 115, 117–137. doi:10.1016/j.neuroimage.2015.04.042

Iglesias, J.E., Van Leemput, K., Bhatt, P., Casillas, C., Dutt, S., Schuff, N., Truran-Sacrey, D., Boxer, A., Fischl, B., Alzheimer’s Disease Neuroimaging Initiative, 2015b. Bayesian segmentation of brainstem structures in MRI. Neuroimage 113, 184–195. doi:10.1016/j.neuroimage.2015.02.065

Ikram, M.A., Brusselle, G.G.O., Murad, S.D., van Duijn, C.M., Franco, O.H., Goedegebure, A., Klaver, C. C.W., Nijsten, T.E.C., Peeters, R.P., Stricker, B.H., Tiemeier, H., Uitterlinden, A.G., Vernooij, M.W., Hofman, A., 2017. The Rotterdam Study: 2018 update on objectives, design and main results. Eur. J. Epidemiol. 32, 807–850. doi:10.1007/s10654-017-0321-4

Jahanshad, N., Thompson, P.M., 2017. Multimodal neuroimaging of male and female brain structure in health and disease across the life span. J. Neurosci. Res. 95, 371–379. doi:10.1002/jnr.23919

Jernigan, T.L., Brown, T.T., Hagler, D.J., Akshoomoff, N., Bartsch, H., Newman, E., Thompson, W.K., Bloss, C.S., Murray, S.S., Schork, N., Kennedy, D. N., Kuperman, J.M., McCabe, C., Chung, Y., Libiger, O., Maddox, M., Casey, B.J., Chang, L., Ernst, T.M., Frazier, J.A., Pediatric Imaging, Neurocognition and Genetics Study, 2016. The pediatric imaging, neurocognition, and genetics (PING) data repository. Neuroimage 124, 1149–1154. doi:10.1016/j.neuroimage.2015.04.057

Joliot, M., Jobard, G., Naveau, M., Delcroix, N., Petit, L., Zago, L., Crivello, F., Mellet, E., Mazoyer, B., Tzourio-Mazoyer, N., 2015. AICHA: An atlas of intrinsic connectivity of homotopic areas. J. Neurosci. Methods 254, 46–59. doi:10.1016/j.jneumeth.2015.07.013

Jones, D.K., Knösche, T.R., Turner, R., 2013. White matter integrity, fiber count, and other fallacies: the do’s and don’ts of diffusion MRI. Neuroimage 73, 239–254. doi:10.1016/j.neuroimage.2012.06.081

Kaczkurkin, A.N., Raznahan, A., Satterthwaite, T.D., 2019. Sex differences in the developing brain: insights from multimodal neuroimaging. Neuropsychopharmacology 44, 71–85. doi:10.1038/s41386-018-0111-z

Kivipelto, M., Helkala, E.L., Hänninen, T., Laakso, M.P., Hallikainen, M., Alhainen, K., Soininen, H., Tuomilehto, J., Nissinen, A., 2001. Midlife vascular risk factors and late-life mild cognitive impairment: A population-based study. Neurology 56, 1683–1689. doi:10.1212/wnl.56.12.1683

Klein, A., Tourville, J., 2012. 101 labeled brain images and a consistent human cortical labeling protocol. Front. Neurosci. 6, 171. doi:10.3389/fnins.2012.00171

Kochunov, P., Glahn, D.C., Lancaster, J., Thompson, P.M., Kochunov, V., Rogers, B., Fox, P., Blangero, \ J., Williamson, D.E., 2011. Fractional anisotropy of cerebral white matter and thickness of cortical gray matter across the lifespan. Neuroimage 58, 41–49. doi:10.1016/j.neuroimage.2011.05.050

Kodiweera, C., Alexander, A.L., Harezlak, J., McAllister, T.W., Wu, Y.-C., 2016. Age effects and sex differences in human brain white matter of young to middle-aged adults: A DTI, NODDI, and q-space study. Neuroimage 128, 180–192. doi:10.1016/j.neuroimage.2015.12.033

Kovacs, G.G., Adle-Biassette, H., Milenkovic, I., Cipriani, S., van Scheppingen, J., Aronica, E., 2014. Linking pathways in the developing and aging brain with neurodegeneration. Neuroscience 269, 152–172. doi:10.1016/j.neuroscience.2014.03.045

Kurtzer, G.M., Sochat, V., Bauer, M.W., 2017. Singularity: Scientific containers for mobility of compute. PLoS ONE 12, e0177459. doi:10.1371/journal.pone.0177459

Lebel, C., Deoni, S., 2018. The development of brain white matter microstructure. Neuroimage 182, 207–218. doi:10.1016/j.neuroimage.2017.12.097

Lebel, C., Gee, M., Camicioli, R., Wieler, M., Martin, W., Beaulieu, C., 2012. Diffusion tensor imaging of white matter tract evolution over the lifespan. Neuroimage 60, 340–352. doi:10.1016/j.neuroimage.2011.11.094

Lemaître, H., Crivello, F., Grassiot, B., Alpérovitch, A., Tzourio, C., Mazoyer, B., 2005. Age- and sex-related effects on the neuroanatomy of healthy elderly. Neuroimage 26, 900–911. doi:10.1016/j.neuroimage.2005.02.042

Loeffler, M., Engel, C., Ahnert, P., Alfermann, D., Arelin, K., Baber, R., Beutner, F., Binder, H., Brähler, E., Burkhardt, R., Ceglarek, U., Enzenbach, C., Fuchs, M., Glaesmer, H., Girlich, F., Hagendorff, A., Häntzsch, M., Hegerl, U., Henger, S., Hensch, T., Thiery, J., 2015. The LIFE-Adult-Study: objectives and design of a population-based cohort study with 10,000 deeply phenotyped adults in Germany. BMC Public Health 15, 691. doi:10.1186/s12889-015-1983-z

Madan, C.R., 2018. Age differences in head motion and estimates of cortical morphology. PeerJ 6, e5176. doi:10.7717/peerj.5176

Mah, A., Geeraert, B., Lebel, C., 2017. Detailing neuroanatomical development in late childhood and early adolescence using NODDI. PLoS ONE 12, e0182340. doi:10.1371/journal.pone.0182340

Makris, N., Goldstein, J.M., Kennedy, D., Hodge, S.M., Caviness, V.S., Faraone, S.V., Tsuang, M. T., Seidman, L.J., 2006. Decreased volume of left and total anterior insular lobule in schizophrenia. Schizophr. Res. 83, 155–171. doi:10.1016/j.schres.2005.11.020

Marcus, D.S., Fotenos, A.F., Csernansky, J.G., Morris, J.C., Buckner, R.L., 2010. Open access series of imaging studies: longitudinal MRI data in nondemented and demented older adults. J. Cogn. Neurosci. 22, 2677–2684. doi: 10.1162/jocn.2009.21407

Mazoyer, B., Mellet, E., Perchey, G., Zago, L., Crivello, F., Jobard, G., Delcroix, N., Vigneau, M., Leroux, G., Petit, L., Joliot, M., Tzourio-Mazoyer, N., 2016. BIL&GIN: A neuroimaging, cognitive, behavioral, and genetic database for the study of human brain lateralization. Neuroimage 124, 1225–1231. doi:10.1016/j.neuroimage.2015.02.071

McKay, D.R., Knowles, E.E.M., Winkler, A.A.M., Sprooten, E., Kochunov, P., Olvera, R.L., Curran, J.E., Kent, J.W., Carless, M.A., Göring, H.H.H., Dyer, T.D., Duggirala, R., Almasy, L., Fox, P.T., Blangero, J., Glahn, D.C., 2014. Influence of age, sex and genetic factors on the human brain. Brain Imaging Behav. 8, 143–152. doi:10.1007/s11682-013-9277-5

Merluzzi, A.P., Dean, D.C., Adluru, N., Suryawanshi, G.S., Okonkwo, O.C., Oh, J.M., Hermann, B.P., Sager, M.A., Asthana, S., Zhang, H., Johnson, S.C., Alexander, A.L., Bendlin, B.B., 2016. Age-dependent differences in brain tissue microstructure assessed with neurite orientation dispersion and density imaging. Neurobiol. Aging 43, 79–88. doi:10.1016/j.neurobiolaging.2016.03.026

Mills, K.L., Goddings, A.-L., Herting, M.M., Meuwese, R., Blakemore, S.-J., Crone, E.A., Dahl, R.E., Güroğlu, B., Raznahan, A., Sowell, E.R., Tamnes, C.K., 2016. Structural brain development between childhood and adulthood: Convergence across four longitudinal samples. Neuroimage 141, 273–281. doi:10.1016/j.neuroimage.2016.07.044

Montagni, I., Cariou, T., Tzourio, C., González-Caballero, J.-L., 2019. “I don’t know”, “I’m not sure”, “I don’t want to answer”: a latent class analysis explaining the informative value of nonresponse options in an online survey on youth health. Int. J. Soc. Res. Methodol. 1–17. doi:10.1080/13645579.2019.1632026

Mori, S., Oishi, K., Jiang, H., Jiang, L., Li, X., Akhter, K., Hua, K., Faria, A.V., Mahmood, A., Woods, R., Toga, A.W., Pike, G.B., Neto, P.R., Evans, A., Zhang, J., Huang, H., Miller, M.I., van Zijl, P., Mazziotta, J., 2008. Stereotaxic white matter atlas based on diffusion tensor imaging in an ICBM template. Neuroimage 40, 570–582. doi:10.1016/j.neuroimage.2007.12.035

Mutlu, A.K., Schneider, M., Debbané, M., Badoud, D., Eliez, S., Schaer, M., 2013. Sex differences in thickness, and folding developments throughout the cortex. Neuroimage 82, 200–207. doi:10.1016/j.neuroimage.2013.05.076

Nordenskjöld, R., Malmberg, F., Larsson, E.-M., Simmons, A., Ahlström, H., Johansson, L., Kullberg, J., 2015. Intracranial volume normalization methods: considerations when investigating gender differences in regional brain volume. Psychiatry Res. 231, 227–235. doi:10.1016/j.pscychresns.2014.11.011

Oishi, K., Zilles, K., Amunts, K., Faria, A., Jiang, H., Li, X., Akhter, K., Hua, K., Woods, R., Toga, A.W., Pike, G.B., Rosa-Neto, P., Evans, A., Zhang, J., Huang, H., Miller, M.I., van Zijl, P.C.M., Mazziotta, J., Mori, S., 2008. Human brain white matter atlas: identification and assignment of common anatomical structures in superficial white matter. Neuroimage 43, 447–457. doi:10.1016/j.neuroimage.2008.07.009

Pausova, Z., Paus, T., Abrahamowicz, M., Bernard, M., Gaudet, D., Leonard, G., Peron, M., Pike, G.B., Richer, L., Séguin, J.R., Veillette, S., 2017. Cohort profile: the saguenay youth study (SYS). Int. J. Epidemiol. 46, e19. doi:10.1093/ije/dyw023

Petersen, R.C., Aisen, P.S., Beckett, L.A., Donohue, M.C., Gamst, A.C., Harvey, D.J., Jack, C.R., Jagust, W.J., Shaw, L.M., Toga, A.W., Trojanowski, J.Q., Weiner, M.W., 2010. Alzheimer’s Disease Neuroimaging Initiative (ADNI): clinical characterization. Neurology 74, 201–209. doi:10.1212/WNL.0b013e3181cb3e25

Pfefferbaum, A., Sullivan, E.V., 2015. Cross-sectional versus longitudinal estimates of age-related changes in the adult brain: overlaps and discrepancies. Neurobiol. Aging 36, 2563–2567. doi:10.1016/j.neurobiolaging.2015.05.005

Pohl, K.M., Sullivan, E.V., Rohlfing, T., Chu, W., Kwon, D., Nichols, B.N., Zhang, Y., Brown, S. A., Tapert, S.F., Cummins, K., Thompson, W.K., Brumback, T., Colrain, I.M., Baker, F.C., Prouty, D., De Bellis, M.D., Voyvodic, J.T., Clark, D.B., Schirda, C., Nagel, B.J., Pfefferbaum, A., 2016. Harmonizing DTI measurements across scanners to examine the development of white matter microstructure in 803 adolescents of the NCANDA study. Neuroimage 130, 194–213. doi:10.1016/j.neuroimage.2016.01.061

Potvin, O., Dieumegarde, L., Duchesne, S., Alzheimer’s Disease Neuroimaging Initiative, 2017. Normative morphometric data for cerebral cortical areas over the lifetime of the adult human brain. Neuroimage 156, 315–339. doi:10.1016/j.neuroimage.2017.05.019

Raznahan, A., Lee, Y., Stidd, R., Long, R., Greenstein, D., Clasen, L., Addington, A., Gogtay, N., Rapoport, J.L., Giedd, J.N., 2010. Longitudinally mapping the influence of sex and androgen signaling on the dynamics of human cortical maturation in adolescence. Proc Natl Acad Sci USA 107, 16988–16993. doi:10.1073/pnas.1006025107

Raznahan, A., Shaw, P., Lalonde, F., Stockman, M., Wallace, G.L., Greenstein, D., Clasen, L., Gogtay, N., Giedd, J.N., 2011. How does your cortex grow? J. Neurosci. 31, 7174–7177. doi:10.1523/JNEUROSCI.0054-11.2011

Reuter, M., Tisdall, M.D., Qureshi, A., Buckner, R.L., van der Kouwe, A.J.W., Fischl, B., 2015. Head motion during MRI acquisition reduces gray matter volume and thickness estimates. Neuroimage 107, 107–115. doi:10.1016/j.neuroimage.2014.12.006

Ritchie, S.J., Cox, S.R., Shen, X., Lombardo, M.V., Reus, L.M., Alloza, C., Harris, M.A., Alderson, H.L., Hunter, S., Neilson, E., Liewald, D.C.M., Auyeung, B., Whalley, H.C., Lawrie, S.M., Gale, C.R., Bastin, M.E., McIntosh, A.M., Deary, I.J., 2018. Sex Differences in the Adult Human Brain: Evidence from 5216 UK Biobank Participants. Cereb. Cortex 28, 2959–2975. doi:10.1093/cercor/bhy109

Roalf, D.R., Quarmley, M., Elliott, M.A., Satterthwaite, T. D., Vandekar, S.N., Ruparel, K., Gennatas, E. D., Calkins, M.E., Moore, T.M., Hopson, R., Prabhakaran, K., Jackson, C.T., Verma, R., Hakonarson, H., Gur, R.C., Gur, R.E., 2016. The impact of quality assurance assessment on diffusion tensor imaging outcomes in a large-scale population-based cohort. Neuroimage 125, 903–919. doi:10.1016/j.neuroimage.2015.10.068

R Core Team, 2018. R: A Language and Environment for Statistical Computing.

Sachdev, P.S., Brodaty, H., Reppermund, S., Kochan, N.A., Trollor, J.N., Draper, B., Slavin, M.J., Crawford, J., Kang, K., Broe, G.A., Mather, K.A., Lux, O., Memory and Ageing Study Team, 2010. The Sydney Memory and Ageing Study (MAS): methodology and baseline medical and neuropsychiatric characteristics of an elderly epidemiological non-demented cohort of Australians aged 70-90 years. Int. Psychogeriatr. 22, 1248–1264. doi:10.1017/S1041610210001067

Sachdev, P.S., Lammel, A., Trollor, J.N., Lee, T., Wright, M.J., Ames, D., Wen, W., Martin, N.G., Brodaty, H., Schofield, P.R., OATS research team, 2009. A comprehensive neuropsychiatric study of elderly twins: the Older Australian Twins Study. Twin Res. Hum. Genet. 12, 573–582. doi:10.1375/twin.12.6.573

Salat, D.H., Greve, D.N., Pacheco, J.L., Quinn, B.T., Helmer, K.G., Buckner, R.L., Fischl, B., 2009. Regional white matter volume differences in nondemented aging and Alzheimer’s disease. Neuroimage 44, 1247–1258. doi:10.1016/j.neuroimage.2008.10.030

Satterthwaite, T.D., Connolly, J.J., Ruparel, K., Calkins, M.E., Jackson, C., Elliott, M.A., Roalf, D.R., Hopson, R., Prabhakaran, K., Behr, M., Qiu, H., Mentch, F.D., Chiavacci, R., Sleiman, P.M.A., Gur, R.C., Hakonarson, H., Gur, R.E., 2016. The Philadelphia Neurodevelopmental Cohort: A publicly available resource for the study of normal and abnormal brain development in youth. Neuroimage 124, 1115–1119. doi:10.1016/j.neuroimage.2015.03.056

Schumann, G., Loth, E., Banaschewski, T., Barbot, A., Barker, G., Büchel, C., Conrod, P.J., Dalley, J. W., Flor, H., Gallinat, J., Garavan, H., Heinz, A., Itterman, B., Lathrop, M., Mallik, C., Mann, K., Martinot, J.L., Paus, T., Poline, J.B., Robbins, T.W., IMAGEN consortium, 2010. The IMAGEN study: reinforcement-related behaviour in normal brain function and psychopathology. Mol. Psychiatry 15, 1128–1139. doi:10.1038/mp.2010.4

Seiler, S., Pirpamer, L., Hofer, E., Duering, M., Jouvent, E., Fazekas, F., Mangin, J.-F., Chabriat, H., Dichgans, M., Ropele, S., Schmidt, R., 2014. Magnetization transfer ratio relates to cognitive impairment in normal elderly. Front. Aging Neurosci. 6, 263. doi:10.3389/fnagi.2014.00263

Seshadri, S., Wolf, P.A., Beiser, A., Elias, M.F., Au, R., Kase, C.S., D’Agostino, R.B., DeCarli, C., 2004. Stroke risk profile, brain volume, and cognitive function: the Framingham Offspring Study. Neurology 63, 1591–1599. doi:10.1212/01.wnl.0000142968.22691.70

Simmonds, D.J., Hallquist, M.N., Asato, M., Luna, B., 2014. Developmental stages and sex differences of white matter and behavioral development through adolescence: a longitudinal diffusion tensor imaging (DTI) study. Neuroimage 92, 356–368. doi:10.1016/j.neuroimage.2013.12.044

Slater, D.A., Melie-Garcia, L., Preisig, M., Kherif, F., Lutti, A., Draganski, B., 2019. Evolution of white matter tract microstructure across the life span. Hum. Brain Mapp. 40, 2252–2268. doi:10.1002/hbm.24522

Smith, S.M., Jenkinson, M., Johansen-Berg, H., Rueckert, D., Nichols, T.E., Mackay, C.E., Watkins, K.E., Ciccarelli, O., Cader, M.Z., Matthews, P.M., Behrens, T.E.J., 2006. Tract-based spatial statistics: voxelwise analysis of multi-subject diffusion data. Neuroimage 31, 1487–1505. doi:10.1016/j.neuroimage.2006.02.024

Smith, S.M., Jenkinson, M., Woolrich, M.W., Beckmann, C.F., Behrens, T.E.J., Johansen-Berg, H., Bannister, P.R., De Luca, M., Drobnjak, I., Flitney, D.E., Niazy, R.K., Saunders, J., Vickers, J., Zhang, Y., De Stefano, N., Brady, J.M., Matthews, P.M., 2004. Advances in functional and structural MR image analysis and implementation as FSL. Neuroimage 23 Suppl 1, S208–19. doi:10.1016/j.neuroimage.2004.07.051

Sotiras, A., Toledo, J.B., Gur, R.E., Gur, R.C., Satterthwaite, T.D., Davatzikos, C., 2017. Patterns of coordinated cortical remodeling during adolescence and their associations with functional specialization and evolutionary expansion. Proc Natl Acad Sci USA 114, 3527–3532. doi:10.1073/pnas.1620928114

Sowell, E.R., Peterson, B.S., Kan, E., Woods, R.P., Yoshii, J., Bansal, R., Xu, D., Zhu, H., Thompson, P.M., Toga, A.W., 2007. Sex differences in cortical thickness mapped in 176 healthy individuals between 7 and 87 years of age. Cereb. Cortex 17, 1550–1560. doi:10.1093/cercor/bhl066

Storsve, A.B., Fjell, A.M., Tamnes, C.K., Westlye, L.T., Overbye, K., Aasland, H.W., Walhovd, K.B., 2014. Differential longitudinal changes in cortical thickness, surface area and volume across the adult life span: regions of accelerating and decelerating change. J. Neurosci. 34, 8488–8498. doi:10.1523/JNEUROSCI.0391-14.2014

Suzuki, Y., Matsuzawa, H., Kwee, I.L., Nakada, T., 2003. Absolute eigenvalue diffusion tensor analysis for human brain maturation. NMR Biomed. 16, 257–260. doi:10.1002/nbm.848

Tamnes, C.K., Herting, M.M., Goddings, A.-L., Meuwese, R., Blakemore, S.-J., Dahl, R.E., Güroğlu, B., Raznahan, A., Sowell, E.R., Crone, E.A., Mills, K.L., 2017. Development of the Cerebral Cortex across Adolescence: A Multisample Study of Inter-Related Longitudinal Changes in Cortical Volume, Surface Area, and Thickness. J. Neurosci. 37, 3402–3412. doi:10.1523/JNEUROSCI.3302-16.2017

Tukey, J.W., 1977. Exploratory Data Analysis, 1st ed. Pearson, Reading, Mass.

Vijayakumar, N., Mills, K.L., Alexander-Bloch, A., Tamnes, C.K., Whittle, S., 2018. Structural brain development: A review of methodological approaches and best practices. Dev. Cogn. Neurosci. 33, 129–148. doi:10.1016/j.dcn.2017.11.008

Vos, S.B., Jones, D.K., Viergever, M.A., Leemans, A., 2011. Partial volume effect as a hidden covariate in DTI analyses. Neuroimage 55, 1566–1576. doi:10.1016/j.neuroimage.2011.01.048

Wajman, J.R., Mansur, L.L., Yassuda, M.S., 2018. Lifestyle Patterns as a Modifiable Risk Factor for Late-life Cognitive Decline: A Narrative Review Regarding Dementia Prevention. Curr. Aging Sci. 11, 90–99. doi:10.2174/1874609811666181003160225

Walhovd, K.B., Fjell, A.M., Reinvang, I., Lundervold, A., Dale, A.M., Eilertsen, D.E., Quinn, B.T., Salat, D., Makris, N., Fischl, B., 2005. Effects of age on volumes of cortex, white matter and subcortical structures. Neurobiol. Aging 26, 1261–70; discussion 1275. doi:10.1016/j.neurobiolaging.2005.05.020

Walhovd, K.B., Westlye, L.T., Amlien, I., Espeseth, T., Reinvang, I., Raz, N., Agartz, I., Salat, D.H., Greve, D.N., Fischl, B., Dale, A.M., Fjell, A.M., 2011. Consistent neuroanatomical age-related volume differences across multiple samples. Neurobiol. Aging 32, 916–932. doi:10.1016/j.neurobiolaging.2009.05.013

Wang, Y., Adamson, C., Yuan, W., Altaye, M., Rajagopal, A., Byars, A.W., Holland, S.K., 2012. Sex differences in white matter development during adolescence: a DTI study. Brain Res. 1478, 1–15. doi:10.1016/j.brainres.2012.08.038

Wardlaw, J.M., Smith, C., Dichgans, M., 2013. Mechanisms of sporadic cerebral small vessel disease: insights from neuroimaging. Lancet Neurol. 12, 483–497. doi:10.1016/S1474-4422(13)70060-7

Westlye, L.T., Walhovd, K.B., Dale, A.M., Bjørnerud, A., Due-Tønnessen, P., Engvig, A., Grydeland, H., Tamnes, C.K., Ostby, Y., Fjell, A.M., 2010. Life-span changes of the human brain white matter: diffusion tensor imaging (DTI) and volumetry. Cereb. Cortex 20, 2055–2068. doi:10.1093/cercor/bhp280

Whalley, L.J., Dick, F.D., McNeill, G., 2006. A life-course approach to the aetiology of late-onset dementias. Lancet Neurol. 5, 87–96. doi:10.1016/S1474-4422(05)70286-6

White, T., El Marroun, H., Nijs, I., Schmidt, M., van der Lugt, A., Wielopolki, P.A., Jaddoe, V.W.V., Hofman, A., Krestin, G.P., Tiemeier, H., Verhulst, F.C., 2013. Pediatric population-based neuroimaging and the Generation R Study: the intersection of developmental neuroscience and epidemiology. Eur. J. Epidemiol. 28, 99–111. doi:10.1007/s10654-013-9768-0

Whitmer, R.A., Gunderson, E.P., Quesenberry, C.P., Zhou, J., Yaffe, K., 2007. Body mass index in midlife and risk of Alzheimer disease and vascular dementia. Curr. Alzheimer Res. 4, 103–109. doi:10.2174/156720507780362047

Wierenga, L.M., Langen, M., Oranje, B., Durston, S., 2014. Unique developmental trajectories of cortical thickness and surface area. Neuroimage 87, 120–126. doi:10.1016/j.neuroimage.2013.11.010

Winkler, A.M., Sabuncu, M.R., Yeo, B.T.T., Fischl, B., Greve, D.N., Kochunov, P., Nichols, T.E., Blangero, J., Glahn, D.C., 2012. Measuring and comparing brain cortical surface area and other areal quantities. Neuroimage 61, 1428–1443. doi:10.1016/j.neuroimage.2012.03.026

Yang, H., Long, X.-Y., Yang, Y., Yan, H., Zhu, C.-Z., Zhou, X.-P., Zang, Y.-F., Gong, Q.-Y., 2007. Amplitude of low frequency fluctuation within visual areas revealed by resting-state functional MRI. Neuroimage 36, 144–152. doi:10.1016/j.neuroimage.2007.01.054

Zang, Y., Jiang, T., Lu, Y., He, Y., Tian, L., 2004. Regional homogeneity approach to fMRI data analysis. Neuroimage 22, 394–400. doi:10.1016/j.neuroimage.2003.12.030

Zhang, H., Schneider, T., Wheeler-Kingshott, C.A., Alexander, D.C., 2012. NODDI: practical in vivo neurite orientation dispersion and density imaging of the human brain. Neuroimage 61, 1000–1016. doi:10.1016/j.neuroimage.2012.03.072

Zou, Q.-H., Zhu, C.-Z., Yang, Y., Zuo, X.-N., Long, X.-Y., Cao, Q.-J., Wang, Y.-F., Zang, Y.-F., 2008. An improved approach to detection of amplitude of low-frequency fluctuation (ALFF) for resting-state fMRI: fractional ALFF. J. Neurosci. Methods 172, 137–141. doi:10.1016/j.jneumeth.2008.04.012

