## Supplementary material for "The MRi-Share database: brain imaging in a cross-sectional cohort of 1,870 university students": MRi-Share_supplemental

### Description of the ABACI pipelines

Schematic figures for each pipeline follow the general scheme used for the description of UKB pipelines in (Alfaro-Almagro et al., 2018) to facilitate the comparison.

#### T1 and T2-FLAIR structural pipeline

Supplemental Figure 1 shows a flow-chart of the structural pipeline.

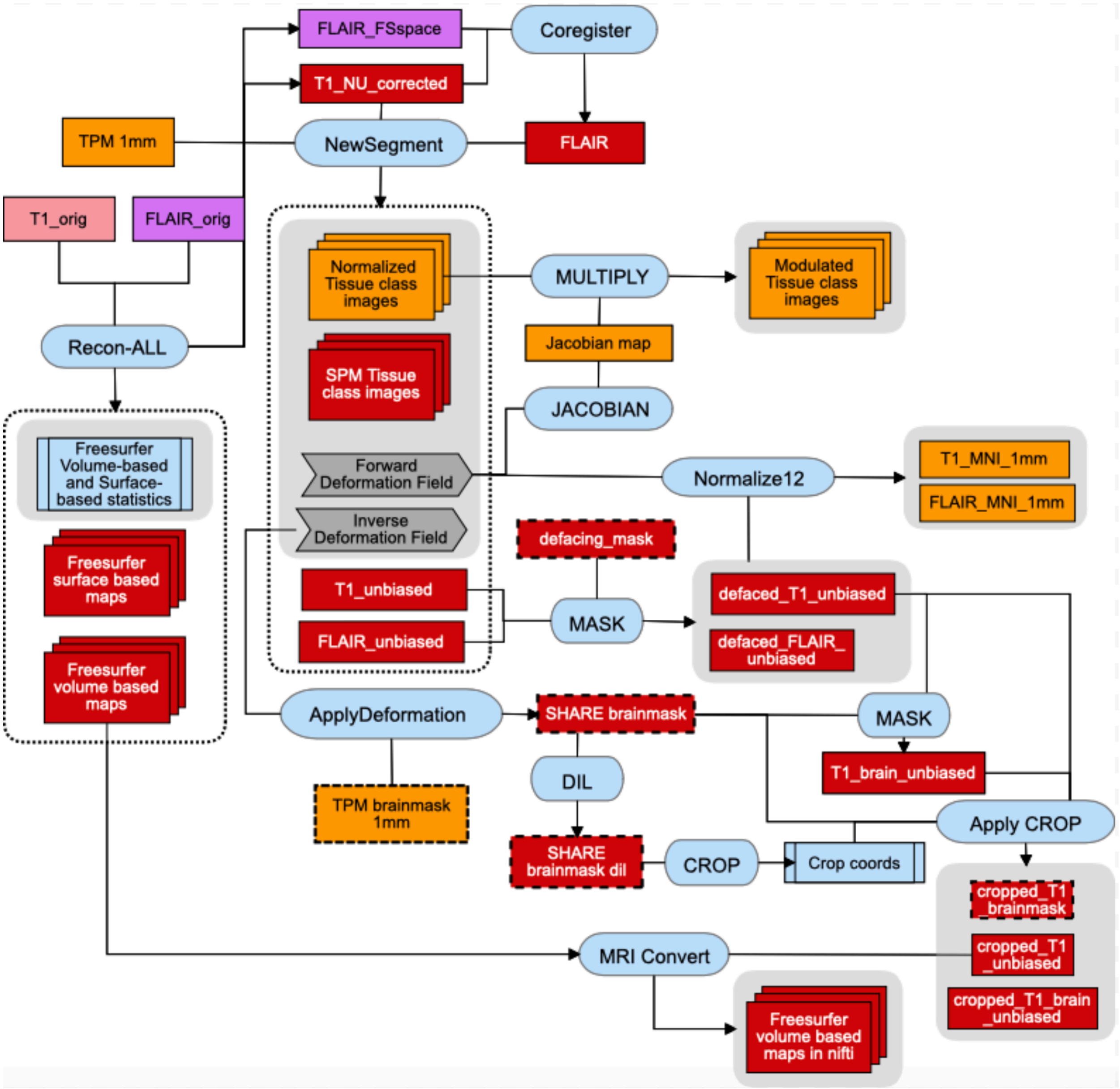

**Supplemental Figure 1: Schematic representation of the structural pipeline.**

In the surface-based processing branch of the structural pipeline, we used the 'recon-all' command of Freesurfer to reconstruct pial and white surfaces of the brain and to obtain both surface- and volume-based metrics from the T1 scan. Briefly, this involves non-parametric non-uniform intensity normalization (N3: Sled et al., 1998), linear transformation to the MNI305 template for estimation of intracranial volume (eTIV), intensity normalization that scales the mean intensity of the white matter to 110, skull-stripping, volume-based linear and non-linear registration to a probabilistic brain atlas for labelling of subcortical structures (Fischl et al., 2004, 2002), and surface-based registration for automatic cortical parcellation (Dale et al., 1999). We used the 'recon-all' command with '-FLAIRpial' multi-channel option to use FLAIR for refining the pial surface, and '-3T' option to use optimized parameters for the N3 correction for data acquired at 3T (based on Zheng et al., 2009) as well as the use of a 3T-based MNI305 template. We also used '-brainstem-structures' to obtain segmentations of 4 brainstem structures (medulla oblongata, pons, midbrain and superior cerebellar peduncle; Iglesias et al., 2015b), and '-hippocampal-subfields-T1T2' flag with FLAIR image input to obtain multi-channel segmentations of hippocampal subfields (Iglesias et al., 2015a).

In a semi-independent branch of the pipeline for volume-based processing, we used the N3-corrected T1 generated by Freesurfer and the FLAIR coregistered to the T1 space as the inputs for multi-channel tissue segmentation using the 'Segment' function of SPM12. This is an extension of the 'Unified Segmentation' framework described in (Ashburner and Friston, 2005) which includes an improved registration method, the option for multi-channel segmentation, as well as the new default tissue probability map (TPM) based on T2-weighted and proton density imaging data from 549 healthy adults (Ashburner et al., 2014). Our initial analysis pipeline used the raw, rather than N3-corrected T1, as the main input, with FLAIR as the secondary input, and with the default parameter setting of 'Segment'. The output images were then used both to generate tissue class maps for creation of the cohort-specific tissue probability map (TPM) template and to compute the final tissue class maps for the voxel-based morphometry analysis. However, the detailed quality control (QC) of individual tissue maps revealed that using the cohort-specific TPM this way somehow greatly biased the GM estimation on frontal regions (Supplemental Figure 2 and 3). After testing in a small subset of subjects (N = 50) as described in the section "Optimization of SPM12 segmentation" below, we 1) used Freesurfer N3-corrected T1 as the main and FLAIR coregistered to the T1 as secondary input, 2) optimized smoothing and regularization parameters for the bias field correction within 'Segment' function, and 3) kept the default TPM packaged with SPM12, rather than the cohort-specific TPM template for the unified tissue segmentation and normalization.

Using the inverse deformation field output from this step, a standard-space brain mask created from the default TPM template was transformed into the native T1 space, and this mask was applied to the bias field corrected T1 to generate a brain-extracted T1. We also used a dilated brain mask in native T1 space to compute the cropping coordinates to reduce image dimension, removing some voxels outside the brain. We then applied the cropping to both the brain-extracted and non-extracted (but defaced) T1 images. These cropped T1 images serve as the reference anatomical space for aligning other image modalities, as well as for converting Freesurfer volume-based maps (e.g. cortical and subcortical parcellation maps) to nifti format.

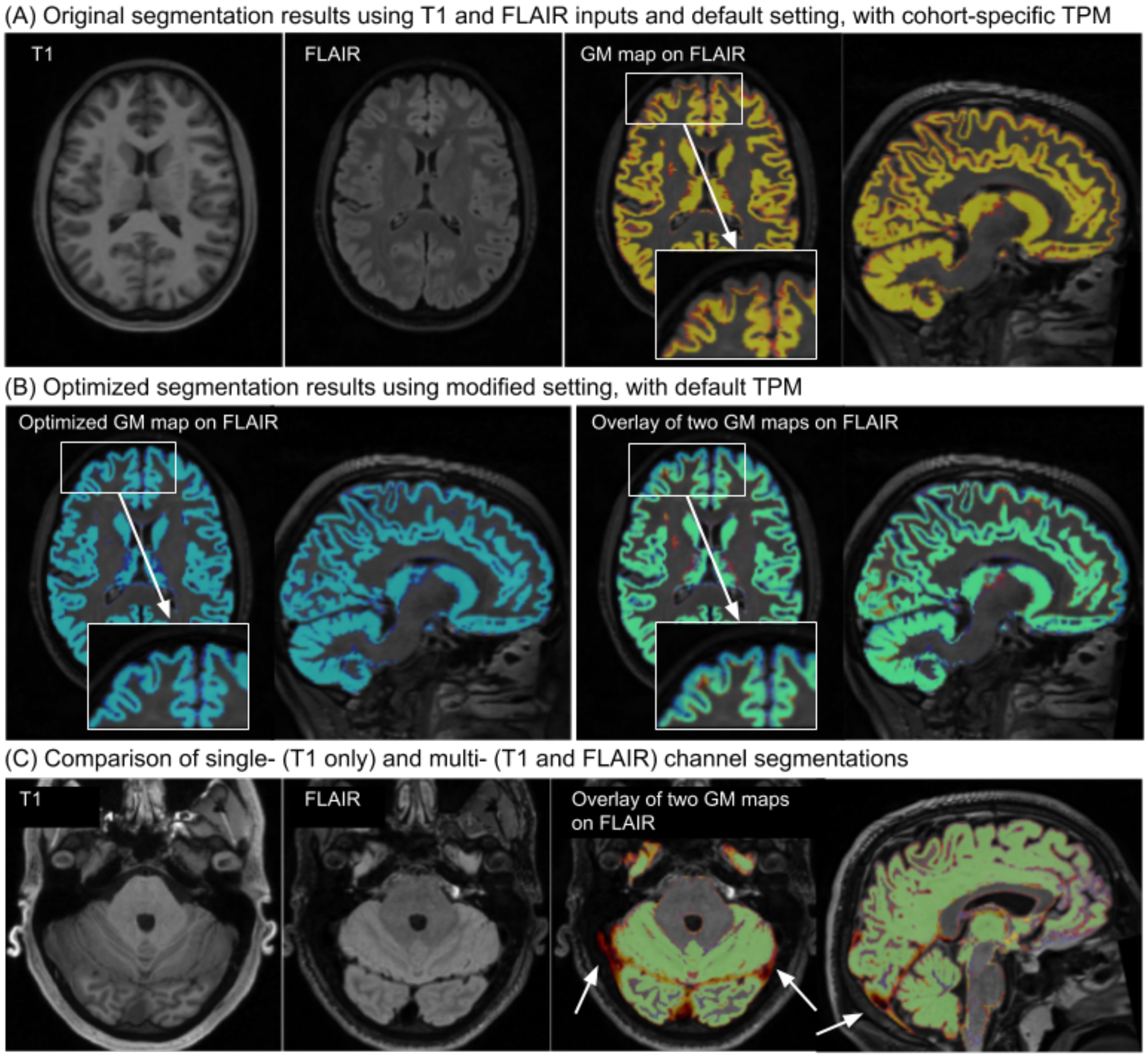

**Supplemental Figure 2. Examples of segmentation failures in single- and multi-channel segmentation procedure in SPM12 and improvement after the optimization.**

*(A) original multi-channel segmentation using T1 and FLAIR inputs, with cohort-specific TPM and using default parameter settings of SPM12, (B) when using only T1 image, with default TPM and default parameter settings, and (C) when using the optimized procedure, with multi-channel segmentation with N3-corrected T1 and par2 setting. (A) and (B) show the same axial and sagittal slices from a representative subject, with T1 and FLAIR axial slices shown in the two left panels of (A). GM map from original segmentation is shown using warm color scale in (A) right, while that from optimized segmentation is shown using cold color scale in (B) left. (B) right shows the overlay of the original (warm) and optimized (color) GM maps. In (C) T1, FLAIR images of another subject (2 left panels), along with the overlay of the single- (warm color scale) and multi-channel (cold color scale) segmentation of GM maps (right panel) are shown. Arrows point to mis-segmentations of meninges as GM in single-channel, T1-only segmentation*.

We computed the Jacobian modulation using the 'Deformations' utility in SPM12, and used this image to scale the spatially normalized tissue class maps for the final morphometric analysis. We used this approach rather than outputting the "modulated" normalized tissue class maps directly from 'Segment', due to prominent artifacts (see Supplemental Figure 4 for an example) possibly related to 'aliasing artifacts' mentioned by the creators of SPM12 (Ashburner et al., 2014).

For Freesurfer-based IDPs, we extracted surface- (CT, inner CSA, and volume for each region of Desikan-Killiany, DKT, and Destrieux cortical atlas regions, as well as pial CSA for Desikan-Killany regions) and volume-based metrics by extracting values from lh/rh.aparc.stats and aseg.stats tables generated automatically with 'recon-all' function. For pial CSA, we used 'mris_anatomical_stats' function to generate additional summary tables for each region in DKT and Destrieux atlases, and extracted values from these tables.

For SPM-based IDPs, we computed total tissue volumes from Jacobian-modulated tissue class maps for GM, WM, and CSF. Additionally, Harvard-Oxford cortical and subcortical atlases (Desikan et al., 2006; Frazier et al., 2005; Goldstein et al., 2007; Makris et al., 2006) and Diedrichsen probabilistic cerebellar atlas (Diedrichsen et al., 2009) were used to obtain total tissue volumes within each region defined by these atlases. The cortical atlas was split by hemisphere to obtain tissue volume estimates separately for the left and right hemispheres.

#### Optimization of SPM12 segmentation

Initially, we created a cohort-specific TPM from the first 500 participants from the MRi-Share sample, and used this TPM for the unified segmentation and normalization procedure as implemented in SPM12, using all the default parameters and with multi-channel setting that used T1 and FLAIR images as inputs. However, the visual QC of the resulting GM segmentation image (Supplemental Figure 3A) showed underestimation of GM that was clearly visible across the majority of subjects and most prominent in frontal pole regions. The degree of underestimation was striking when compared against Freesurfer-based cortical segmentation in the same subject.

As the Freesurfer-based cortical segmentations were noticeably more accurate in many subjects, we decided to use the information about the discrepancy between Freesurfer- and SPM12-based segmentation to compare the effects of modifications on our SPM12 pipeline in a subset of the MRi-Share data (N = 50). More specifically, we computed and compared the mean of SPM12-based GM tissue probability (in native T1 space) inside the Freesurfer-based cortical parcellations after performing SPM12 segmentation procedure with the combinations of the following: 1) TPM (default TPM vs cohort-specific TPM), 2) input channel (T1 only vs multi-channel using T1 and FLAIR), 3) bias-field correction parameter setting of the SPM12 (*regularization* and the *bias field smoothing* settings: default vs two set of parameters as described below), and 4) bias-field correction on T1 *prior* to the correction applied internally within the SPM12 unified segmentation framework (none vs N3 vs N4 correction).

We compared the default and cohort-specific TPM, because it appeared that the use of default TPM seemed to mitigate the underestimation of the GM in frontal regions in our preliminary investigations. We also compared multi-channel (T1 + FLAIR) against T1-only segmentation since most published studies use only T1, although in theory multimodal segmentation should improve performance, and at least one study found empirically this to be the case (Lindig et al., 2018). We tested two different sets of bias-field correction parameter settings against the default since it has been reported that default correction parameters were suboptimal both on simulated (Ganzetti et al., 2016a) and actual (Ganzetti et al., 2016b) MR images. While the default parameters for bias-field correction in SPM12 use $10^{-4}$ and 60 for the *regularization* and *bias-field smoothing*, respectively, Ganzetti *et al.* (2016a) reported $10^{-5}$ and 140 performed the best on their simulated 3T bias-field data, and in their second study (Ganzetti et al., 2016b) reported $10^{-2}$ and 30 to be the most optimal on the actual 3T dataset taken from the KIRBY21 database (https://www.nitrc.org/frs/shownotes.php?release_id=2178). We tested these three sets of settings (*def*: $10^{-4}$ and 60, *par2*: $10^{-5}$ and 140, *par3*: $10^{-2}$ and 30 for the *regularization* and *bias-field smoothing*, respectively). Even though this internal bias-field correction integrated with brain segmentation should be sufficient, and may even be superior to correction methods that are performed independently from segmentation (Ganzetti et al., 2016a), we tested the effects of applying two other commonly used correction methods, namely the N3 (Sled et al., 1998) and N4 (Tustison et al., 2010) prior to SPM12 segmentation.

Supplemental Figure 3 compares the degree of discrepancy between Freesurfer- and SPM12-based GM segmentation for different settings of the SPM12 as described above. They show that the use of cohort-specific TPM generally increases the discrepancy between the two algorithms (i.e. lower overall mean SPM12-based GM tissue inside Freesurfer-defined cortical GM), and also that the level of discrepancy becomes more variable across subjects, as indicated by larger standard deviations. Although single-channel segmentation using T1 image generally increases the amount of SPM12-based GM tissue inside Freesurfer-defined cortical GM, it also increases the amount of GM segmented *outside,* mis-segmenting the meninges around the cortex as GM (Supplemental Figure 2B). This is the case regardless of other parameter settings (e.g. Lindig et al., 2018), since meninges in T1 have similar contrast as GM. Within multi-channel segmentations using SPM12 default TPM, the amount of mean GM content discrepancy across cortical GM is similar across different T1 preprocessing options when using *def* or *par2* settings, but slightly worsens when using *par3* setting (Supplemental Figure 3A). However, region-by-region summary of GM content discrepancy shows that the discrepancy in the frontal pole region is most improved when using N3-corrected T1 image with *par2* setting (red arrows in Supplemental Figure 3B), without noticeably affecting the discrepancy in other regions. Visual inspection of the segmentation in a small number of subjects (Supplemental Figure 2C) corroborated this observation. Thus, for our pipeline we decided to 1) use SPM12 default TPM rather than cohort-specific template, 2) keep multi-channel with both T1 and FLAIR, but 3) use N3-corrected T1 from Freesurfer stream, and 4) adjust the *regularization* and *bias-field smoothing* settings in 'Segment' to $10^{-5}$ and 140, respectively.

(A) Comparison of mean SPM GM content in Freesurfer cortical GM

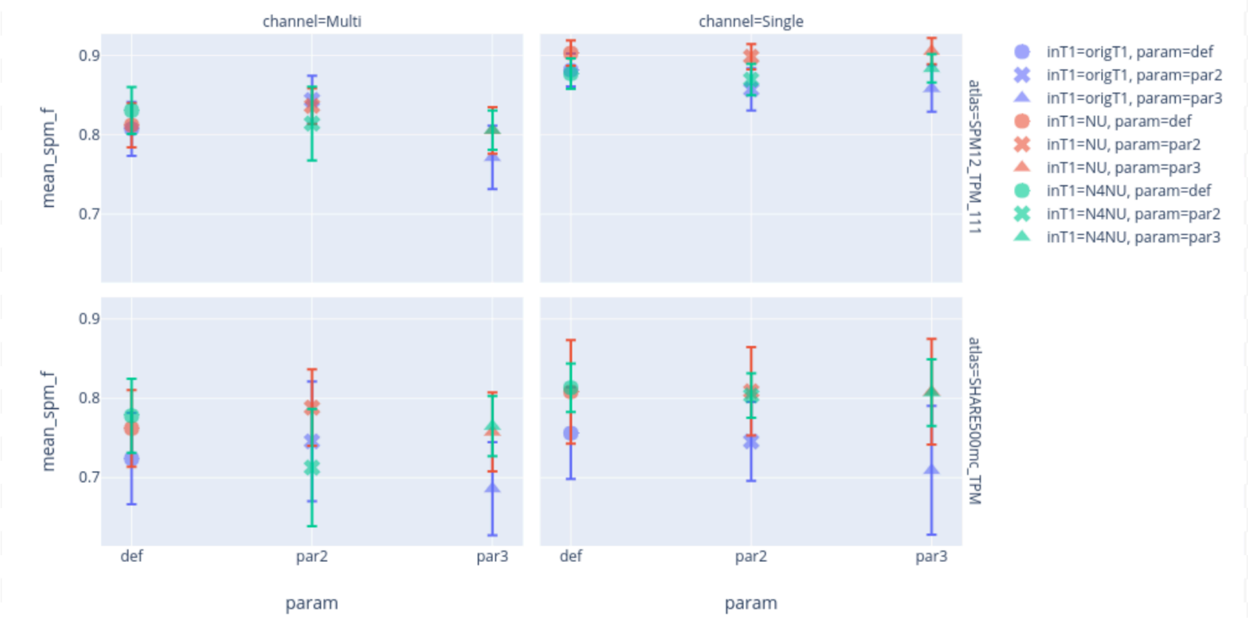

(B) Comparison of mean SPM GM content in Freesurfer Desikan cortical parcellation

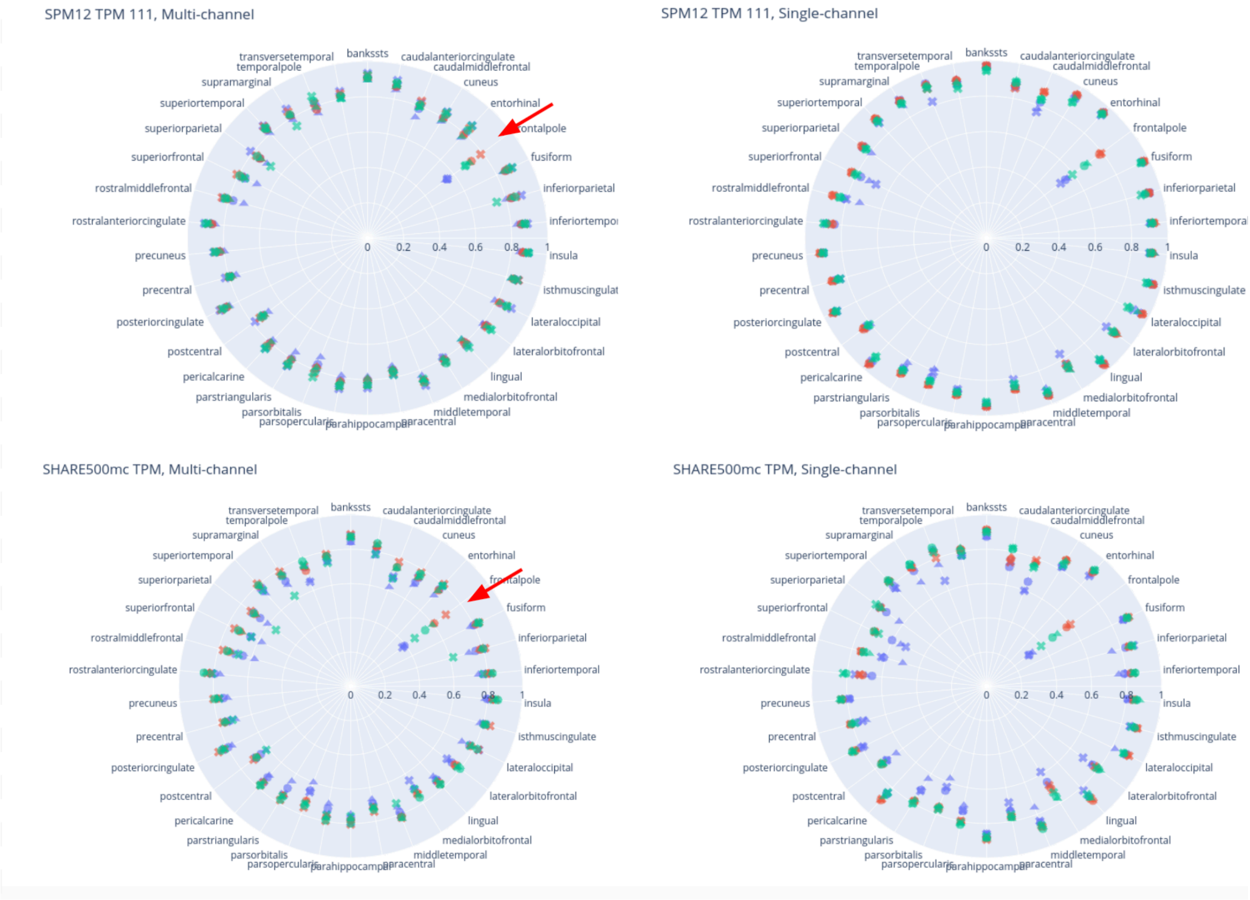

**Supplemental Figure 3. Results of tests for optimizing SPM12 segmentation.**

*Plots show the amount of discrepancy between Freesurfer-based segmentation of the cortical GM and SPM12-based GM probability when using different inputs and parameter settings in SPM12 'Segment' as described in the text. They summarize the mean SPM-based GM content within (A) cortical GM and (B) each region of the Desikan cortical parcellation, as segmented by Freesurfer. The mean values are calculated across the subset of 50 MRiShare subjects, and the error bars indicate the standard deviation. In both (A) and (B), the colors of each data point indicate any bias field correction applied prior to performing 'Segment' (blue: no correction, pink: N3 correction based on Freesurfer, green: N4 correction using ANTS), and the symbols represent parameter settings in 'Segment' (circle: default setting, cross: par2 setting, triangle: par3 setting). Columns on the left use both T1 and FLAIR images as input (i.e. multi-channel segmentation), while the right columns only use T1 (i.e. single-channel segmentation). Upper rows in each (A) and (B) use the default TPM template upsampled to 1mm isotropic resolution, and lower rows use the cohort-specific template derived from 500 MRiShare subjects. Red arrows in (B) indicate the reduced discrepancy in the frontal pole region when using the N3-corrected T1 image with par2 settings*.

#### Artefact in Jacobian-modulated maps produced by SPM12 'Segment' function

Jacobian-modulated maps of tissue class images in normalized space can be outputted directly in the 'Segment' function of SPM12 (denoted with prefix *mwc* for modulated, warped, class image). However, early visual inspection of this image revealed stripe-shaped artefacts in all images examined. These artefacts were not present if the modulated maps were produced "traditional" way by using the 'Deformations' utility in SPM12 to obtain the Jacobian modulation and then using it to scale the warped (i.e. spatially normalized) tissue class images (Supplemental Figure 4). Therefore, in our pipeline we included these extra steps to compute modulated tissue class images in normalized space rather than using the same map outputted directly from the 'Segment' function.

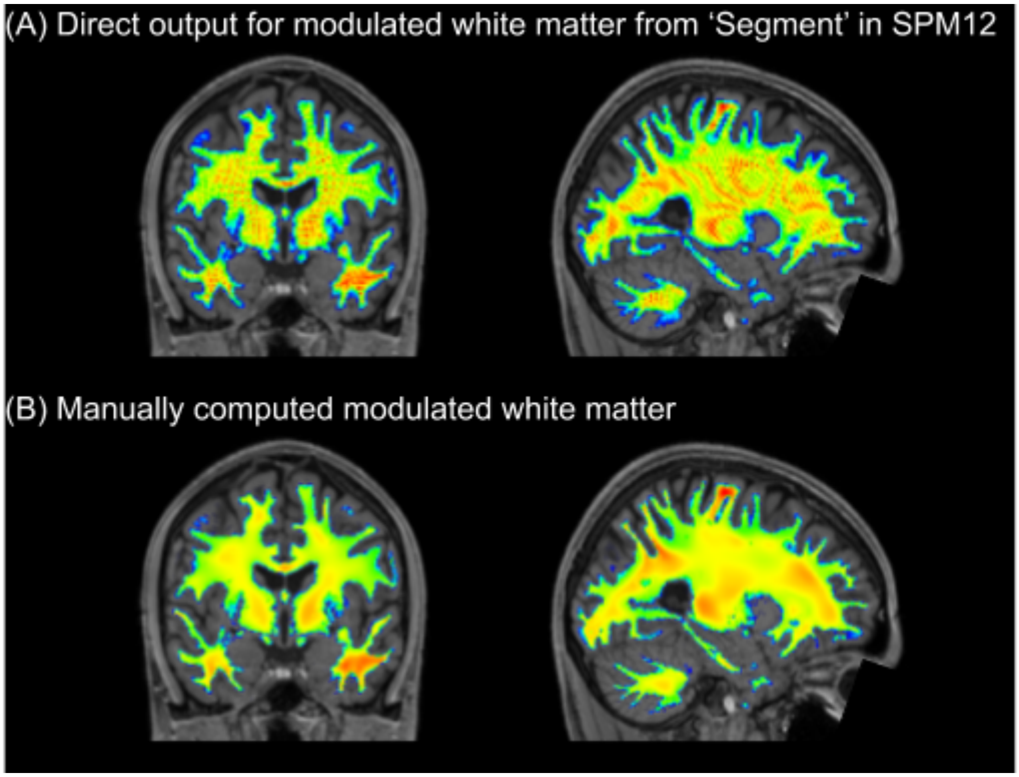

**Supplemental Figure 4. Artefact in modulated maps produced by SPM12.**

*Images show (A) an example of striped artefacts observed in the modulated maps produced with the 'Segment' function in SPM12, and (B) the modulated map computed in the "traditional" way for the same subject. Selected coronal (left panel) and sagittal (right panel) views of the modulated white matter map, overlaid on T1 image warped to standard template space from a representative subject are shown.*

#### Fieldmap generation pipeline

In the fieldmap generation pipeline (see Supplemental Figure 5), eight pairs of b=0 images with the opposing phase-encoding directions are extracted from DWI, merged, and fed into FSL TOPUP tool (Andersson et al., 2003). The resulting field coefficient map and movement parameter text file are kept for the later DWI processing pipeline. The TOPUP tool also produces unwarped, or distortion-corrected input b=0 images. The unwarped AP/PA b=0 images are averaged to create a fieldmap magnitude image, which is then skull-stripped with FSL BET tool (Smith, 2002), and linearly aligned to the reference T1 brain image using FSL FLIRT (Jenkinson et al., 2002; Jenkinson and Smith, 2001). The same linear transformation is also applied to the fieldmap phase image. Both the fieldmap phase and magnitude images in T1 structural space are kept for EPI unwarping in the later rs-fMRI pipeline.

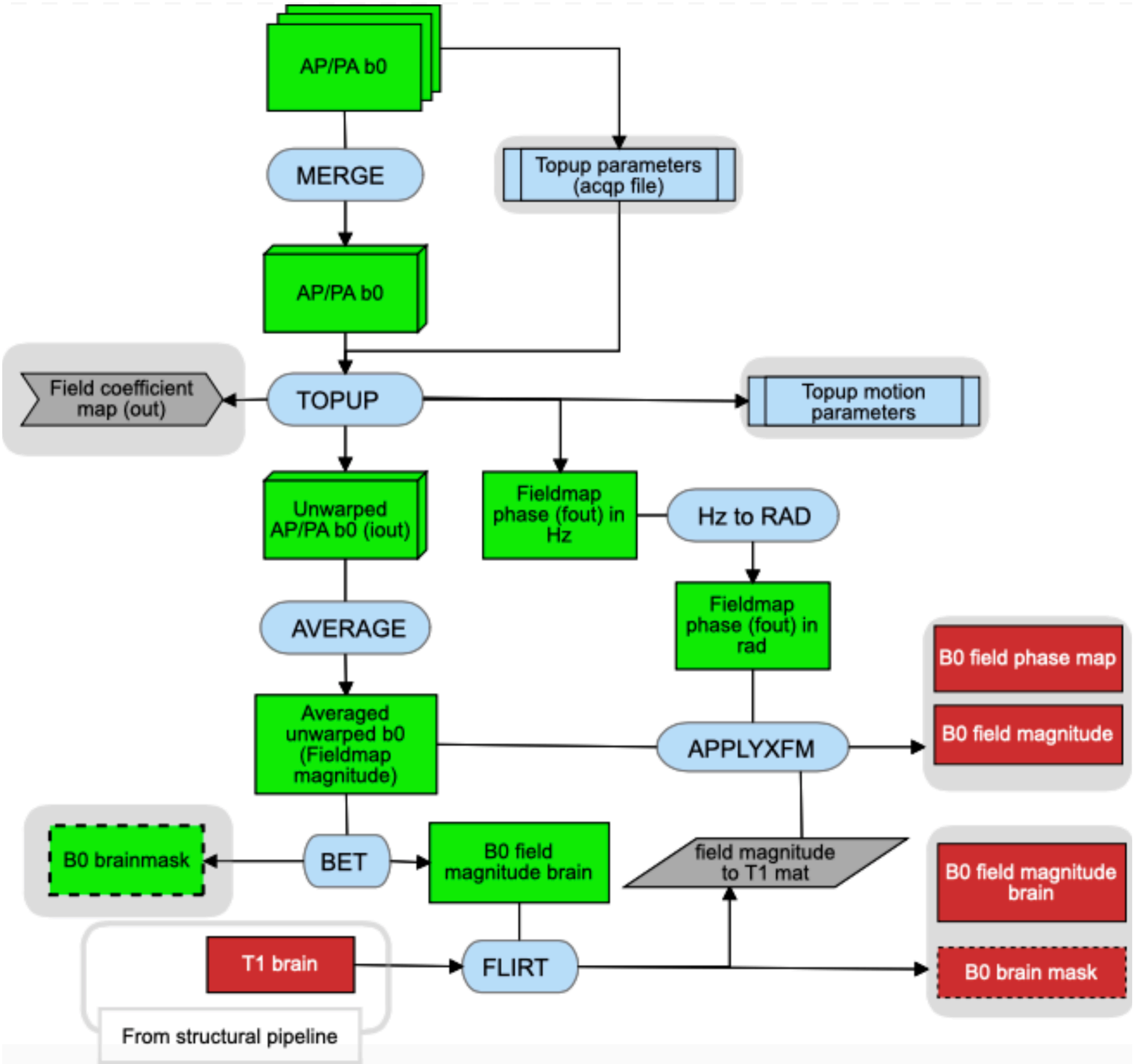

**Supplemental Figure 5: Schematic representation of fieldmap generation pipeline**

#### Diffusion MRI pipeline

Supplemental Figure 6 summarizes the DWI processing pipeline. In this pipeline, individual raw DWI data are first corrected for eddy current (EC) and top-up distortion using the FSL Eddy tool, with replacement of outlier slices (eddy_openmp as implemented in FSL v5.0.10 patch; (Andersson et al., 2016; Andersson and Sotiropoulos, 2016). After cropping the data to reduce non-brain tissue volumes, we apply non-local means filter to denoise and boost the SNR, using the 'nlmeans' denoising tool (Coupe et al., 2011, 2008) as implemented in the Dipy package (0.12.0; Garyfallidis et al., 2014). The resulting image is then used to estimate 1) DTI (Diffusion-Tensor Imaging; Basser et al., 1994) model parameters and 2) microstructural NODDI (Neurite Orientation Dispersion and Density Imaging; Zhang et al., 2012) model parameters. For DTI modelling, the volumes with high b-value (b=2000 s/mm$^2$) are removed from the denoised data before fitting the data with the dipy tools (Jensen and Helpern, 2010) to compute DTI maps, namely the maps of maps of fractional anisotropy (FA), mean, axial, and radial diffusivity (MD, AD, and RD, respectively). The diffusivity maps were further cleaned by removing diffusivity value outliers using Random Sample Consensus (RANSAC) approach (Choi et al., 2009), as implemented in the scikit-learn package (0.19.1; https://scikit-learn.org/stable/index.html). The denoising, DTI computation, and the RANSAC outlier removal were performed by wrapping Scilpy scripts, developed by Sherbrooke Connectivity Imaging Lab (https://scilpy.readthedocs.io/en/latest/).

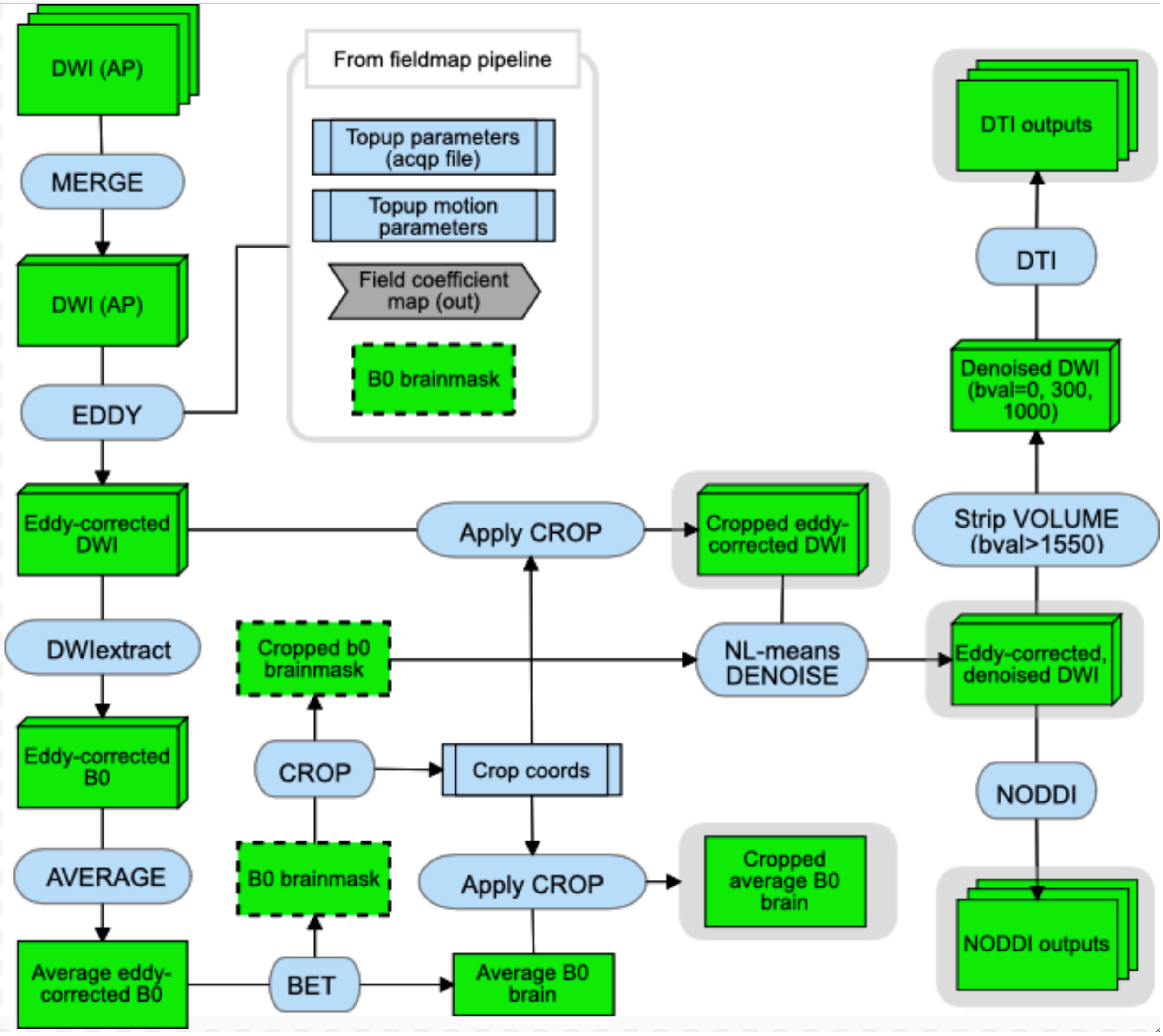

**Supplemental Figure 6: Schematic representation of diffusion MRI pipeline**

Before fitting denoised data for NODDI, we computed empirical values of cohort-specific isotropic and parallel diffusivity by computing the mean MD within lateral ventricles and mean AD within the corpus callosum in individual T1 space for each subject. The mean of these values across subjects were then used as dPar ($1.5 \times 10^{-3}$) and dIso ($2.4 \times 10^{-3}$) parameters in the AMICO (Accelerated Microstructure Imaging via Convex Optimization) tool (Daducci et al., 2015) when fitting the denoised DWI data to obtain the following NODDI metric maps: isotropic volume fraction (IsoVF), which indicates the proportion of free water volume of each voxel, neurite density index (NDI), which represents the proportion of intracellular volume in the remaining fraction, and orientation dispersion index (ODI), a measure of within-voxel fiber dispersion.

In order to obtain mean DTI/NODDI metrics within cerebral WM, we used sub-workflows that first upsample and coregister DTI and NODDI maps in the native DWI space (1.7 mm isotropic) to the individual native T1 space (1mm isotropic), then compute mean values within a cerebral WM mask defined for each subject in their native T1 space. The coregistration sub-workflow used the '*antsRegistrationSyNQuick'* script in the ANTS package to compute a linear (rigid + affine) transform and symmetric diffeomorphic image normalization (deformable SyN; Avants et al., 2008) warp to align the eddy- and motion-corrected average b0 image, upsampled to match T1 resolution, to the individual T1 image, then applied the computed transform to the upsampled DTI and NODDI maps. The subject-specific cerebral WM mask was derived from the Freesurfer segmentation, but it was also refined by masking it with the SPM12-derived native WM tissue probability map thresholded at 0.5. This ensured that the mean DTI/NODDI values were computed within the cerebral WM regions with limited partial volume effects. Other IDPs using Freesurfer labels were generated in a similar fashion, with mean values calculated within each of the subject-specific labels, and then refined by masking it with either the SPM12-based WM (for WM regional labels) or GM (GM regional labels) mask.

To generate IDPs based on JHU ICBM-DTI-81 white matter labels atlas (Mori et al., 2008; Oishi et al., 2008), we used SPM12 'Coregister', followed by 'Normalize' function that used the deformation field generated in the structural pipeline to transform DTI/NODDI maps in the native DWI space to the standard template space (1mm isotropic) in one step. Here, subject-specific, spatially normalized WM class images (also 1mm isotropic) from the structural pipeline were thresholded at 0.5 and used as the mask when computing mean DTI/NODDI values within each region in the atlas.

Finally, IDPs based on spatially normalized WM skeleton were generated using a script based on Baykara et al. (http://www.psmd-marker.com, Baykara et al., 2016), and described in (Beaudet et al., 2020). Briefly, it used FSL TBSS (Smith et al., 2006) to obtain spatially normalized, skeletonized WM maps based on the FA maps for each individual. After removing voxels near the ventricles with a custom mask developed by Baykara et al. (2016), the voxel value distribution for each of the DTI metric was analyzed within the skeletonized WM mask, to obtain mean, standard deviation, and the value between the 95th and 5th percentile values.

#### Resting-state fMRI pipeline

The pipeline used for preprocessing the resting state fMRI (rs-fMRI) is sketched in the Supplemental Figure 7.

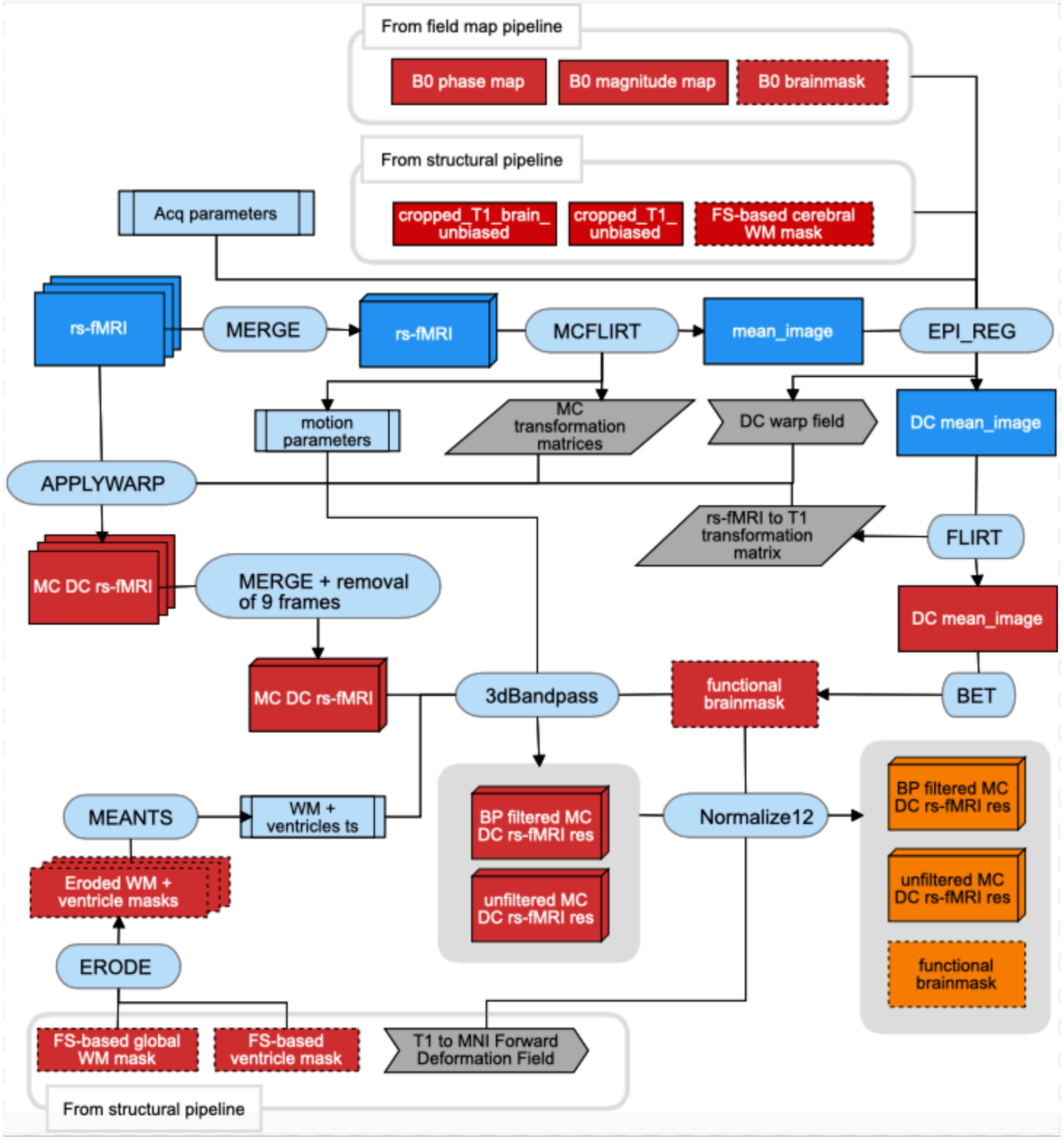

**Supplemental Figure 7. Schematic representation of resting-state fMRI pipeline**

Processing of the 15 minutes long rs-fMRI dataset requires additional data that have been processed with the structural and fieldmap generation pipelines (see above) including the b0-fieldmap, the T1-weighted volume, the nonlinear deformation field for stereotaxic transformation of the T1-weighted volume into the reference brain in the MNI standard space, and four masks based on FreeSurfer segmentation: 1) cerebral WM mask (not including corpus callosum), 2) global WM mask (cerebral WM with corpus callosum and cerebellar WM), 3) ventricle mask, and 4) brain mask.

The pipeline begins with the spatial alignment of the rs-fMRI data into the T1-weighted individual reference space using a 3-step procedure similar to the one described in Alfaro-Almagro et al. 2018. It includes motion correction with the FSL MCFLIRT (Jenkinson et al., 2002), followed by the combined echo-planar imaging (EPI) distortion correction and WM boundary-based registration (BBR) with the FSL 'epi_reg' script packaged with FSL FLIRT, and the final realignment to the T1 space with the FSL FLIRT (Jenkinson et al., 2002; Jenkinson and Smith, 2001). For the BBR, the Freesurfer-based WM mask was used, and the fieldmap phase and magnitude images were used for the EPI distortion correction. Both the BBR/distortion correction and post-BBR coregistrations were computed on the motion-corrected rs-fMRI mean image. For each volume of the rs-fMRI data, the spatial transformations for the three steps (motion correction, BBR/distortion correction, and post-BBR coregistration) were combined and applied together using the FSL 'applywarp' command, in order to minimize the number of interpolations. Note that the 9 first EPI volumes are removed from the analysis in order to take in account initial imperfect stabilisation of the signal. The FD that indexes the subject motion in the rs-fMRI, as described in (Power et al., 2014, 2012) was computed using the BRAMILA tools, developed by Brain and Mind Lab at Aalto University (https://users.aalto.fi/~eglerean/bramila.html).

Once in the individual T1 space, a brain mask was computed from rs-fMRI data with the FSL BET (Smith, 2002). This mask was applied when band-pass filtering the data using the AFNI 3dBandpass (Sforazzini et al., 2016). This step also included despiking and nuisance regression, using the times series of the Friston 24 motion parameters (Friston et al., 1996) and those extracted from the subject-specific, Freesurfer-based WM and ventricle masks. The WM and ventricle masks were eroded 3 and 2 times, respectively, but if the resulting mask contained less than 50 voxels, the mask with one less erosion was used to extract the nuisance time series. We used a frequency window of 0.01 to 0.1 Hz for the band-pass filtered data, and 0 to 99999 Hz to create unfiltered data with the identical despiking and nuisance regression. These images were then warped into the stereotaxic space using the SPM12 'Normalise' function with the deformation field from T1 space to the reference MNI brain generated in the structural pipeline, at a voxel sampling size of 2 mm isotropic.

For computing regional intrinsic connectivity (IC) matrices with and without global signal regression (GSR), the brain mask produced from the rs-fMRI data in T1 space was warped into the stereotaxic space using the same deformation field to transform the rs-fMRI data to the standard MNI space, and mean time series of the band-pass filtered data within this mask was extracted with FSL 'fslmeants'. This time series was then used as a regressor in FSL 'fsl_glm' function to obtain the residual of the band-pass filtered data. The band-pass filtered data with and without GSR were further cleaned to remove the effects of motion by scrubbing, in which we removed any volumes that exceeded the FD of 0.5mm (Power et al., 2014, 2012). We then extracted the average time series within each of the 384 regions (192 homotopic region pairs) of the AICHA atlas (Joliot et al., 2015) from these data, and computed the Pearson correlation coefficients (*r*) between every pair of regions to produce the two versions of regional IC matrices. When computing the group average IC matrices, any subjects with less than 75 % of a given AICHA region overlapping with the functional brain mask were excluded from the computation of the mean value for *r* involving that region (i.e. one row/column for the region).

The band-pass filtered data (but without any scrubbing or GSR) in the standard space were also used to compute three other properties of spontaneous brain activities at rest at a voxel-level: 1) regional homogeneity (ReHo; Zang et al., 2004) 2) amplitude of low frequency fluctuation (ALFF; Yang et al., 2007), and 3) fractional ALFF (fALFF; Zou et al., 2008). ReHo measures the degree of similarity in spontaneous fluctuations in the neighboring voxels, and thus shows how coherent the regional intrinsic neuronal activities are. It is computed as the average of the Kendal's coefficients of concordance between a given voxel time series and those of each of its 26 neighbors that are inside the brain mask. ALFF, in contrast, captures the total power of the regional intrinsic neuronal activity, and is computed as the integral of the square-root of the power spectrum of low-frequency (0.01 to 0.1 Hz) fluctuations in each voxel. Lastly, fALFF describes this low-frequency power as the fraction of total fluctuations in the entire range of frequency, which reportedly increases the sensitivity and specificity to the neuronal activity (Zou et al., 2008). To compute fALFF, the amplitude maps were created for both band-pass filtered and unfiltered data, and the ratio between the two were calculated for each voxel. All three maps were computed in the voxels inside the intersection between the functional brain mask and the Freesurfer-based brain mask, both in the spatially normalized space.

We also performed subject-level independent component analyses (ICA) using FSL MELODIC (Beckmann and Smith, 2004), after spatially smoothing the band-pass filtered data at full width at half maximum of 5 mm with AFNI 3dBlurInMask. The number of dimensions was estimated using the Laplace approximation to the Bayesian evidence of the model order in each subject (Beckmann and Smith, 2004; Minka, 2000)

#### Description of subject-level visual QC and quantitative QC metrics

Each pipeline in the ABACI generated a set of subject-level qualitative, visual QC images as well as quantitative QC metrics. As shown in Supplemental Figure 8, these were organized as linked HTML web pages, where a main page for each modality showed interactive distribution histograms for the main IDPs and/or QC metrics, with links to individual web pages for each participant through columns listing batch numbers and subject numbers per batch (each batch containing 50 subjects to avoid listing entire subjects in a single column). When a viewer hovers over the point in the scatterplot below the histogram (Supplemental Figure 8A), it shows the subject ID and the value of the metric for that subject, and when clicked, it opens the individual web page for that subject (Supplemental Figure 8B).

This way, it facilitates the detection and inspection of outliers for any given metrics within each pipeline. The interactive graphs embedded in the web pages were generated with custom python tools, using *bokeh* package (0.12.16, https://bokeh.org/). In sections below, we describe the QC procedures followed for each modality, as well as the details of qualitative and quantitative QC in each pipeline.

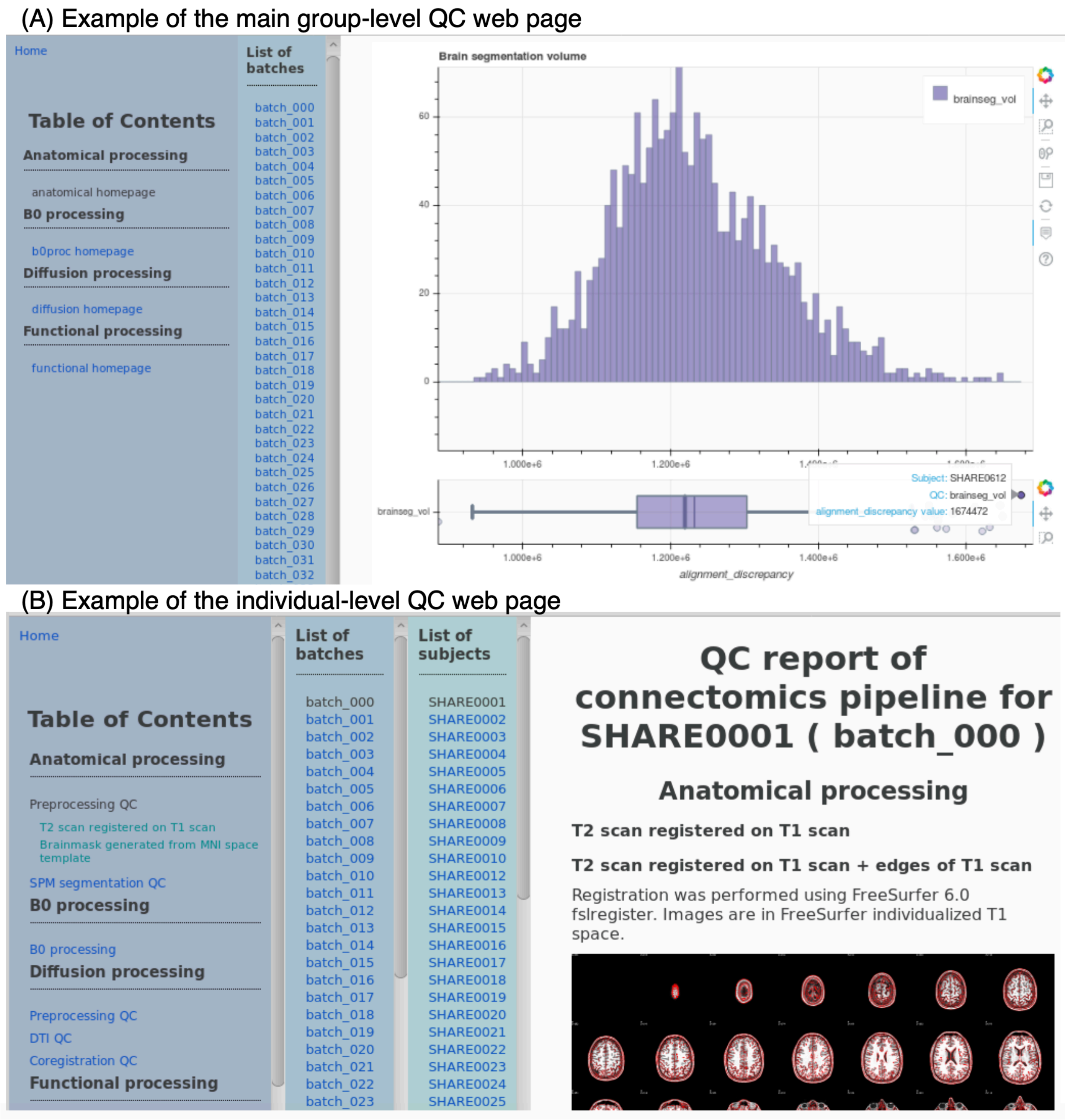

**Supplemental Figure 8. Example of the QC web page produced by the ABACI.**

*The examples of linked QC HTML web pages generated by the ABACI pipelines are shown for the structural pipeline. (A) The main page for the structural pipeline containing the distributions of selected IDPs as interactive histogram and scatter plot. The snapshot was taken when hovering over the subject with highest Freesurfer-segmented brain volume. Clicking on it would open the linked individual-level structural QC page for that subject. (B) An example of the individual-level QC page for the structural pipeline.*

#### Structural MRI processing QC

As a first step, the same three qualified MD investigators (B.M, E.M, and N.T-M) who reviewed raw T1 and FLAIR images for any incidental findings and non-incidental anomalies also flagged any images with visible artefacts, such as ringing and reduced contrast in the raw images. A trained rater (A.T) then rated the flagged images on four categories according to the rating system proposed by Backhausen et al. (2016); 1) Image sharpness, 2) Ringing, 3) Contrast to noise ratio (CNR) of subcortical structures, and 4) CNR of GM and WM. For each category, scores were given such that 0, 1, and 2 represented 'good', 'moderate', and 'bad' quality as described by Backhausen et al. (2016). Supplemental Figure 9 shows the example raw T1 and FLAIR images in a subject who received the worst score for T1 as well as combined T1 and FLAIR, along with the two representative output images from the two streams of structural pipeline. They show that despite the visible artefacts in both T1 and FLAIR images in this subject, there are no major failures in the tissue segmentation by SPM12 or in the surface- and volume-based segmentations in Freesurfer.

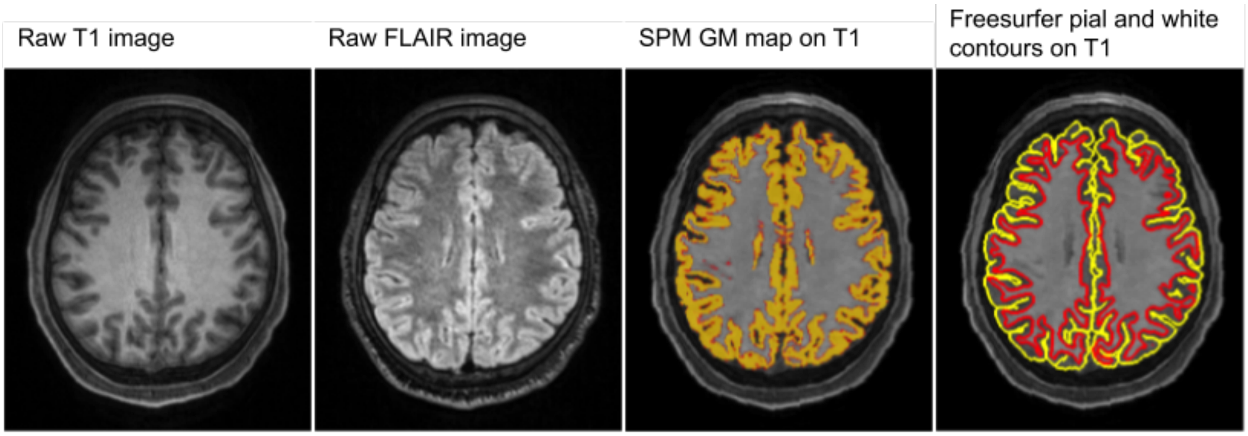

**Supplemental Figure 9. Examples of visible artefacts flagged at the initial check of raw T1 and FLAIR images.**

*Images show raw T1 (left most panel) and FLAIR (second left) images of a subject flagged by the initial review by an MD investigator and later received the worst score of 8 (4 for T1 and FLAIR respectively, using a method suggested by Backhausen et al., 2016). A faint ringing is visible on T1 image, which is also present and more visible on FLAIR image as well. Noise within the white matter is also visible on the FLAIR image. Despite receiving the worst rating for the combined quality of T1 and FLAIR images, both SPM12-based tissue segmentation (third left) and Freesurfer-based surface reconstruction (right most panel) do not present any major problems.*

Independently of the initial flagging of the artefacts, another trained rater (N.B) inspected individual qualitative QC images produced by our pipeline for all scanned participants. These images included the following (see Supplemental Figure 10 for examples of each):

- Multiple axial slices of T1 image with contours of coregistered FLAIR image for checking FLAIR-to-T1 coregistration
- Multi-axial slices of T1 image with brain mask overlay to check the quality of brain mask
- Multiple axial, sagittal, coronal slices of gray and white matter segmented with SPM12, overlaid on FLAIR image to inspect the quality of tissue segmentation

- Jacobian modulated gray and white matter overlaid on the T1 image warped to the template space to check for any unusual deformation.

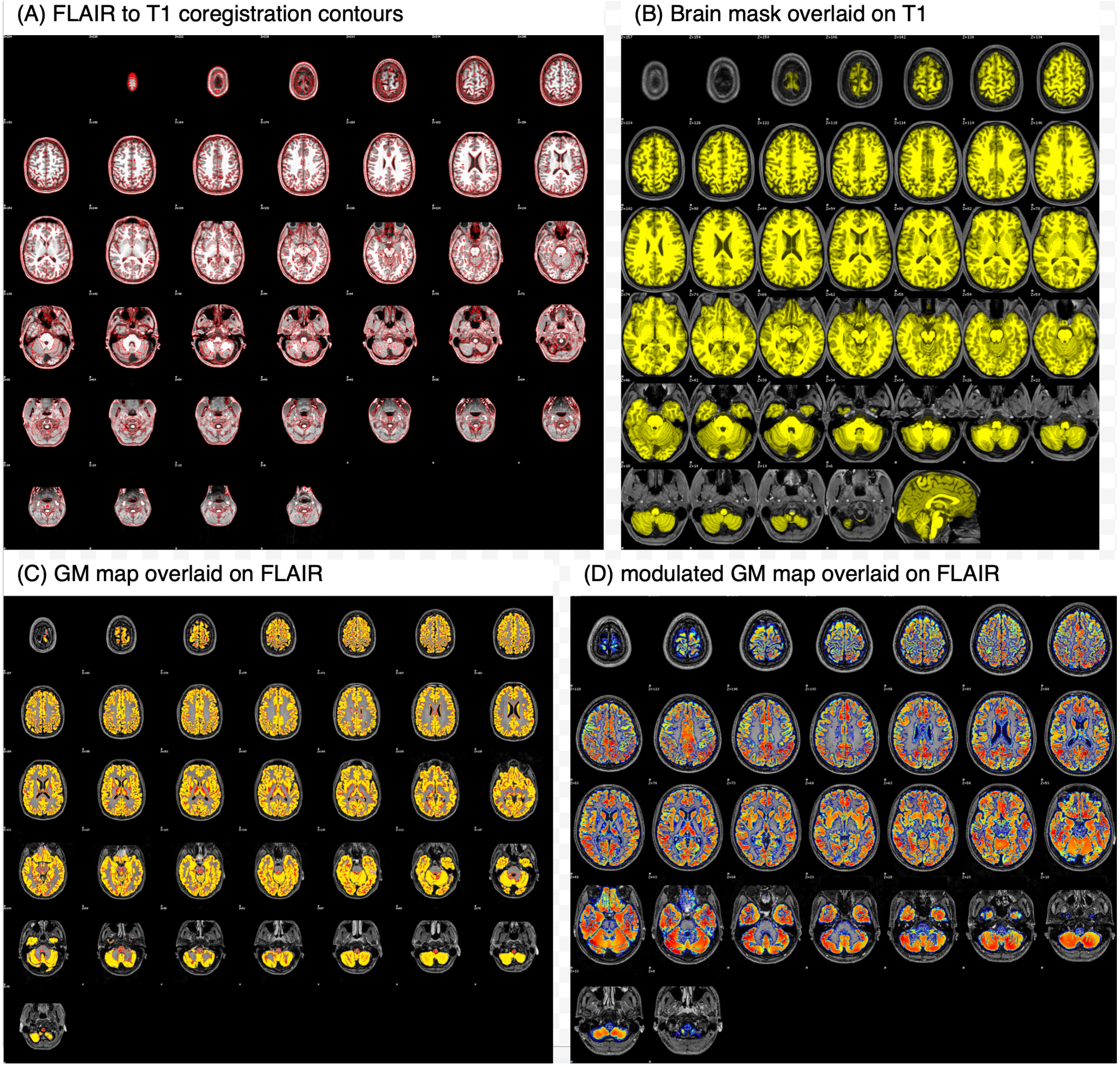

**Supplemental Figure 10. Examples of subject-level visual QC images for T1 and FLAIR structural pipeline.**

*(A) Multiple axial slices of T1 image with contours of coregistered FLAIR image for checking FLAIR-to-T1 coregistration, (B) multi-axial slices of T1 image with brain mask overlaid on them to check the quality of the mask, (C) GM map in native T1 space created by SPM12 overlaid on FLAIR image coregistered to T1 to check the quality of SPM12 tissue segmentation, and (D) modulated GM map overlaid on FLAIR image in the stereotaxic space to check any abnormal deformation. QC images similar to (C) and (D) were created for WM maps as well, and images showing multi-coronal as well as multi-sagittal slices were created for both GM and WM maps.*

The same rater also inspected another set of images and movies produced by a modified version of ENIGMA Cortical QC scripts (package 2.0, April 2017: http://enigma.ini.usc.edu/protocols/imaging-protocols/) that generated multiple views of Freesurfer surface reconstructions and cortical and subcortical parcellations to spot any gross failures in surface reconstruction. The rater followed the ENIGMA Cortical Quality Control Guide 2.0 April 2017 to check for any problems at the regional level in the Desikan cortical parcellation of each subject. While the inspection at the regional level revealed segmentation problems known to be relatively common, no major failures were found at the global surface reconstructions in 1,832 subjects examined. There were also no extreme outliers in global surface-based measures (mean CT, inner and pial CSA) or in volumetric variables (total GM and WM volumes; see Supplemental Figure 16 below in "Additional statistical analysis")

Supplemental Figure 11A shows the distributions of QC metrics computed as part of the structural pipeline and an additional sub-workflow that used the main segmentation output images to compute tissue signal-to-noise (SNR) and contrast-to-noise (CNR). More specifically, we computed:

- Cost function values for FLAIR to T1 coregistration
- Within-tissue SNR in GM and WM (defined as mean divided by standard deviation of the intensity values within these tissues)
- GM-to-WM or GM-to-CSF CNR (defined as the ratio of GM to WM or GM to CSF mean intensity values) for both T1 and FLAIR images, using SPM12-based tissue segmentations.

Similarly to UKB QC IDPs (Alfaro-Almagro et al., 2018), inverse values were computed for both SNR and CNR so as to make higher values of all the QC metrics represent worse quality.

We also computed the following QC metrics to check the distribution and outliers (Supplemental Figure 11B), and to explore their impact on morphometric measures of interest in our future investigation:

- Mean Euler number from the Freesurfer processing, which represents the topological complexity of the reconstructed cortical surface, and which was proposed to be a better index of data quality in structural scan (Rosen et al., 2018)
- Freesurfer total CNR based on *mri_cnr* tool packaged with Freesurfer
- tAverage edge strength (AES) measure, a recently proposed method for retrospectively quantifying the head motion from the structural scan (Zacà et al., 2018).
- Qoala-T score, a new metric for Freesurfer segmented MRI data, which has been proposed as a new quality metric that can be used to compare the quality of data across different datasets with differing acquisition parameters (Klapwijk et al., 2019).

The mean Euler number and AES have been shown in prior studies to affect the age effect estimate on cortical thickness (Madan, 2018; Rosen et al., 2018), even when the most problematic of the images are removed from the analyses. Of note, the mean framewise displacement (FD) from functional MRI (Power et al., 2012) has also been demonstrated to affect the structural morphometry (Madan, 2018; Reuter et al., 2015; Rosen et al., 2018; Savalia et al., 2017). We note that Qoala-T scores in our MRi-Share dataset is particularly high (mean ±

standard deviation [range] = 74.9 ± 9.7 [49.7 - 96.4] %, the maximum score is 100 %), with no images classified below the score of 30%, which is a suggested cut-off for exclusion without further visual QC (Klapwijk et al., 2019). Note that the direction of these extra QC metrics were not manipulated, in order to facilitate the comparison with prior studies using these metrics. The lower values in AES score, Freesurfer Euler number and total CNR, and Qoala-T score represent worse quality.

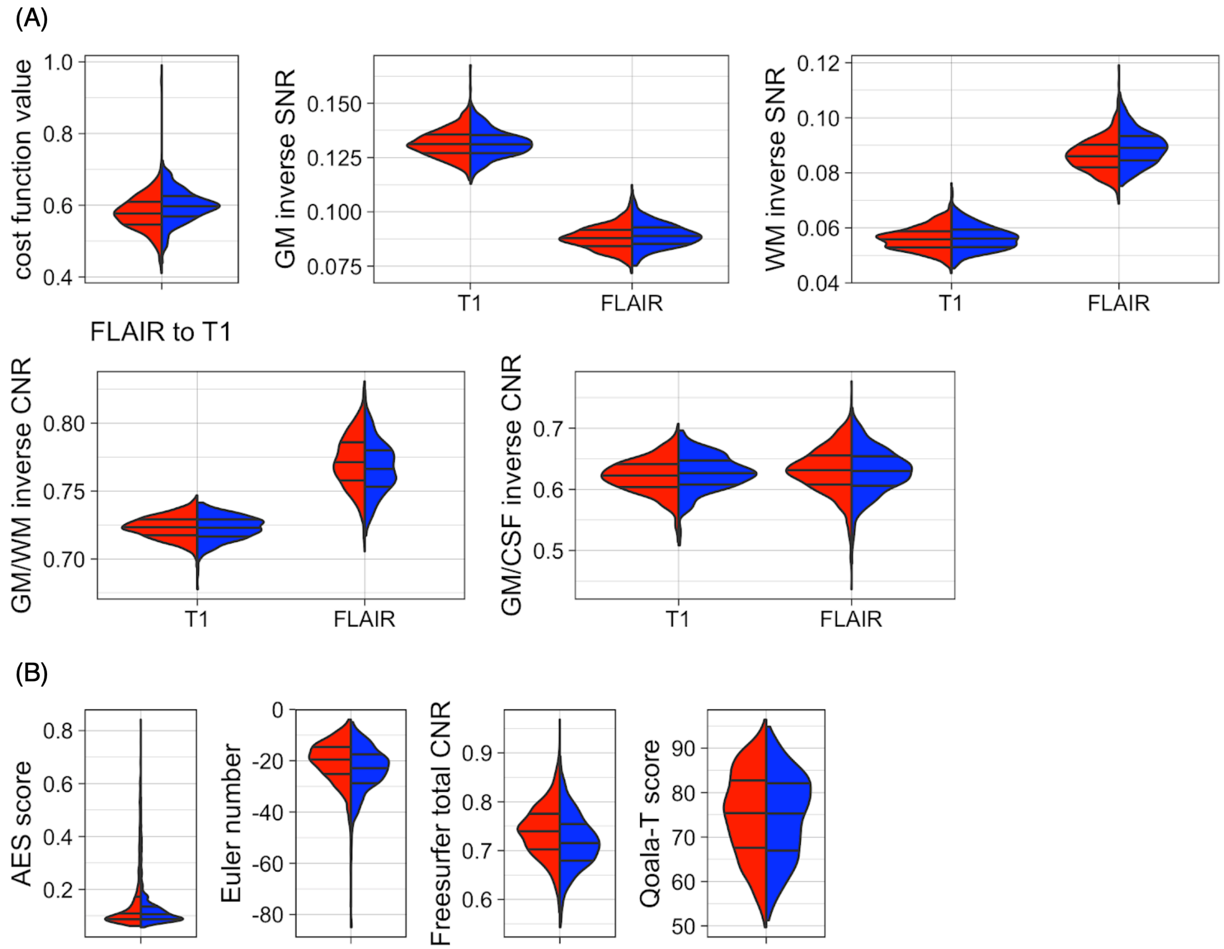

**Supplemental Figure 11. Distributions of quantitative QC metrics for structural pipeline.**

*Distributions of (A) QC IDPs computed in the structural pipeline, and (B) additional IDPs computed outside of our in-house pipeline are presented for male (blue) and female (red) participants. Details of these IDPs are described in the text.*

#### Fieldmap generation processing QC

During the fieldmap generation processing, the following qualitative QC images were produced for each subject:

- Multi-axial plot of B0 brain mask produced by FSL BET overlaid on the mean B0 image to check the quality of brain extraction,
- Multi-axial plot of the mean B0 image coregistered to the native T1 space with the contours of the T1 image to check the quality of B0-to-T1 coregistration.

These images were checked when QC metrics in either DWI or rs-fMRI processing suggested potential problems in the processing of these data, to make sure that they are not caused by the processing error at the fieldmap generation stage.

#### Diffusion MRI processing QC

During the diffusion processing, we generated both qualitative QC images and quantitative QC metrics for each subject. The qualitative QC images included the following (see Supplemental Figure 12 for examples), which were produced based on the QC metric output of FSL EDDY (Bastiani et al., 2019) and other in-house custom tools inspired by QC described in Tournier et al. (2011) and Roalf et al. (2016):

- Plots of motion root mean square (RMS) from FSL EDDY over the entire volumes, or time points, in DWI data. The RNS describes the displacement of each voxel, and therefore the amount of motion from one volume to the next. Both 'total' and 'restricted' displacement are plotted. The former includes distortion due to eddy current distortion, and the latter estimates the amount of displacement caused by the subject motion.
- Plots of slice-level outliers as well as mean and max outlier 'stdev' and 'sqr_stdev' values over the entire volumes in DWI data, also based on FSL EDDY output. These outliers can be caused by the signal dropout in a specific slice (or group of slices) due to subject motion. The 'stdev' value represents the number of standard deviations off the mean difference between the observed and predicted value computed for each slice. The 'sqr_dev' represents the number of standard deviations off the square root of the mean squared difference between the observation and prediction.
- Plots of voxel-level outliers over the entire volumes in DWI data, based on AFNI 3dToutcount, before and after EC correction and denoising
- Mid-sagittal plots of every volume for each b-value after EC correction and denoising
- Maps and intensity histograms of voxel-level CNR for each non-zero-b-value, produced by FSL EDDY
- Multi-axial maps and intensity histograms of temporal SNR (tSNR) for each b-value
- Multi-axial plot of DTI residual map with a mask of physically implausible voxels overlaid on top. The latter mask represents where the mean intensity level of the b = 0 images is below one or more non-zero-b-value images.

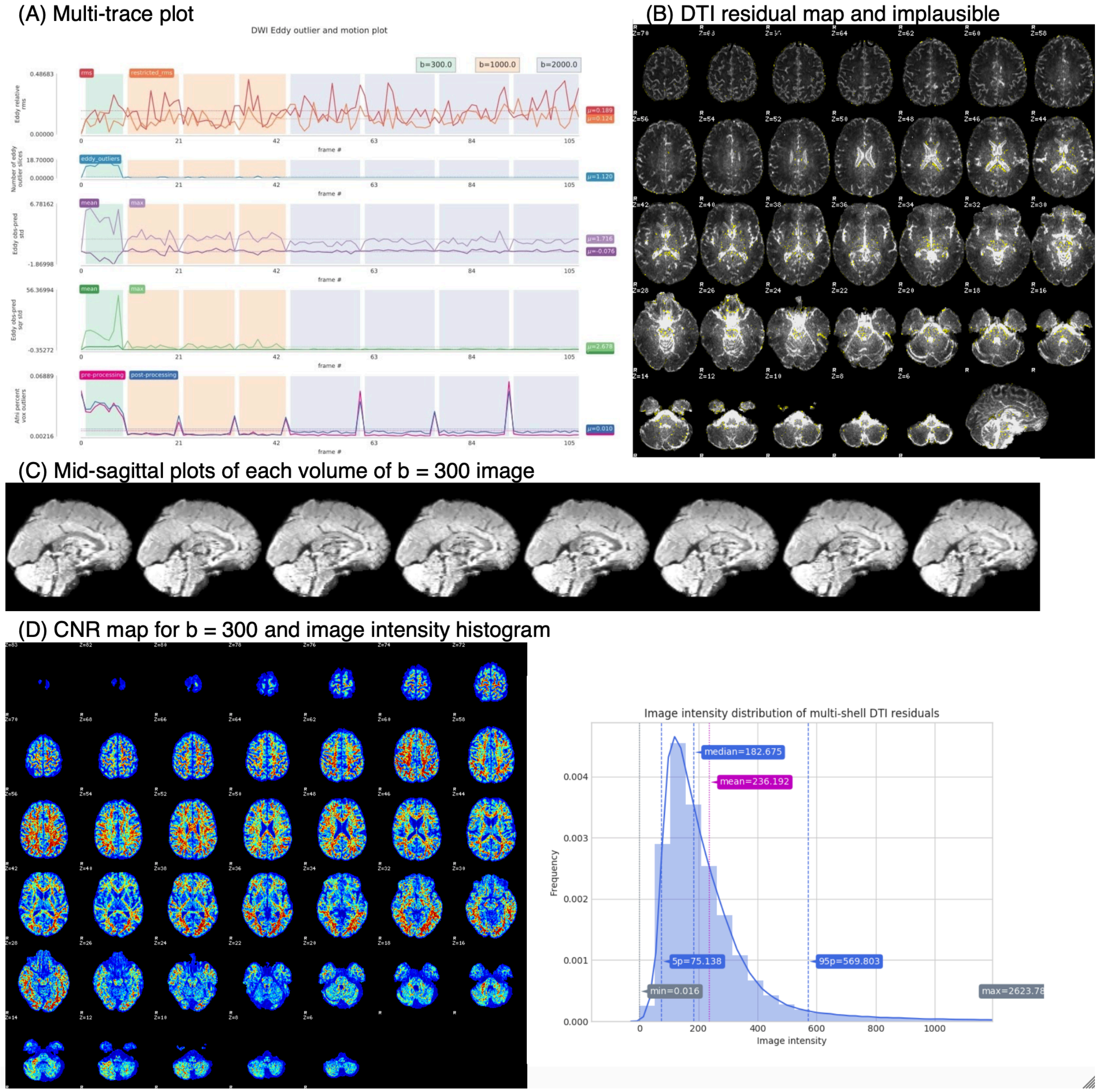

**Supplemental Figure 12. Examples of subject-level visual QC images for diffusion pipeline.**

*(A) Multi-trace plots showing, from the top, 1) 'total' and 'restricted' RMS, 2) number of EDDY outlier slices, 3) mean and max EDDY outlier 'stdev' values, 4) mean and max EDDY outlier 'sqr_stdev' values, and 5) mean fraction of intensity outlier voxels before and after EC correction and denoising. (B) Multi-axial plot of the DTI residual map, with a mask of implausible voxels overlaid on top. (C) Mid-sagittal plots of each volume for each b-value, here showing an example from b = 300 images. (D) Multi-axial plot of CNR and tSNR maps, together with image intensity histograms for these maps, for each non-zero-b-values and b-values for CNR and tSNR, respectively. Here showing the CNR map and histogram for b = 300 as an example.*

Rather than reviewing all web pages for each subject, however, we focused on identifying the outliers in any given quantitative QC metrics, then reviewing the web page for those subjects. Although there are increasing interests in the automated QC in DWI (Bastiani et al., 2019; Haddad et al., 2019; Liu et al., 2015, 2010; Oguz et al., 2014; Roalf et al., 2016; Tournier et al., 2011), there has been less consensus with regard to how and when a given data should be removed from the analysis (Liu et al., 2015), compared to the structural or functional data, where some standards have been suggested, albeit somewhat arbitrarily (e.g. Backhausen et al., 2016 for structural data; Power et al., 2015 for rs-fMRI data). As a result, such decisions depend on within- and across-study comparison of any given QC metrics to spot outlier participants or assess the overall quality of the acquired data. The quantitative QC metrics we computed were mainly derived from the same QC outputs used to generate the visual QC images described above, but represent numerical summary of these outputs. Here we present the distributions of these metrics in our dataset to be compared with other studies (Supplemental Figure 13):

- Mean ‘total’ and ‘restricted’ motion RMS
- Mean number of slices classified as outliers per volume during FSL EDDY
- Mean fraction of voxel intensity outliers in each volume after EC correction and denoising, as determined by AFNI 3dToutcount
- Inverse of the mean CNR inside the brain mask for each of the non-zero-b-values, computed using the voxelwise CNR map produced by FSL EDDY
- Inverse of the tSNR inside the brain mask computed for b-value
- Mean residuals inside the brain mask when fitting DTI model
- Fraction of physically implausible voxels inside the brain mask

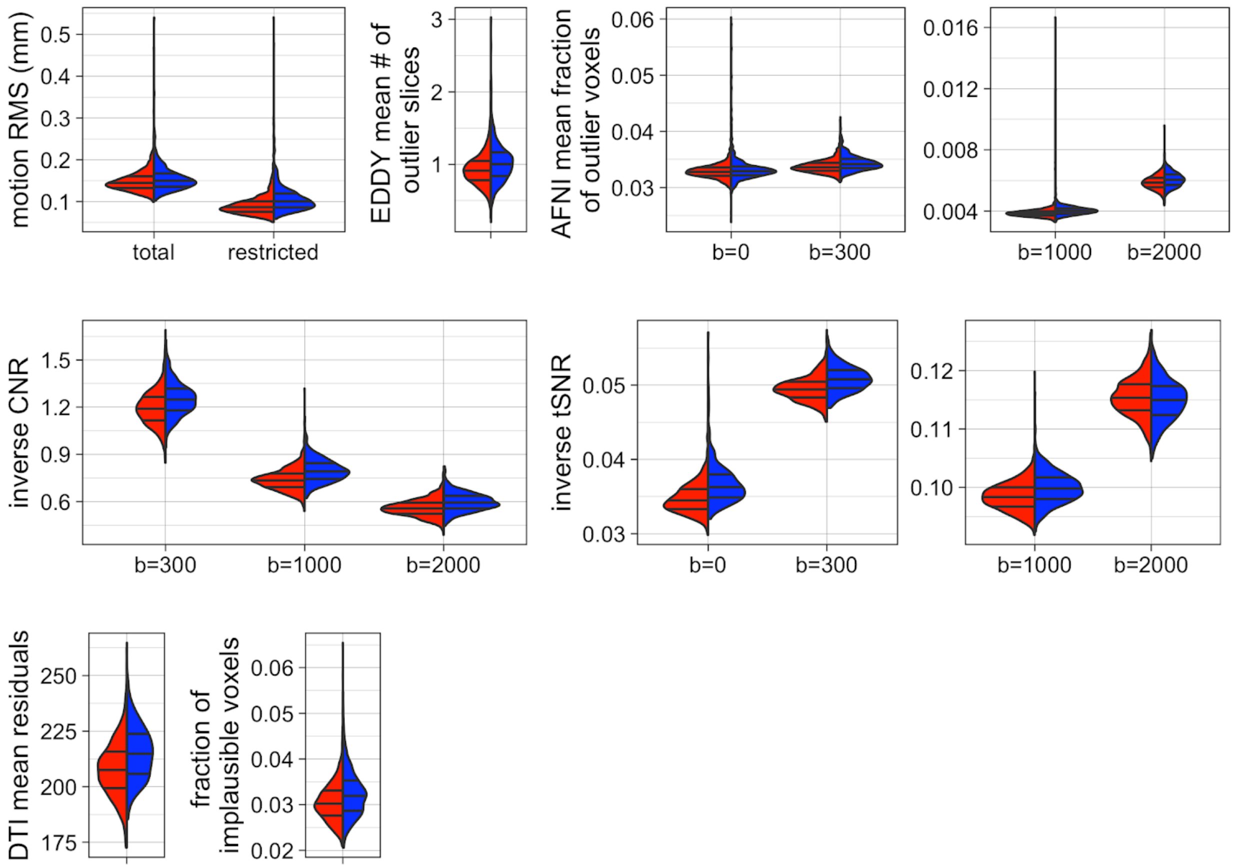

**Supplemental Figure 13. Distributions of quantitative QC metrics for diffusion pipeline.**

*Distributions of QC IDPs from the diffusion pipeline are shown for male (blue) and female (red) participants. See text for details of individual IDPs.*

#### Resting-state fMRI processing QC

As in DWI pipeline, we computed a number of both quantitative QC metrics and qualitative QC images during the pipeline execution. The qualitative QC images included the following (examples shown in Supplemental Figure 14):

- Plots of motion parameters from FSL MCFLIRT and FD over the entire time points of rs-fMRI data
- Plots of global mean signal, as well as mean signals within GM, WM, and ventricles over the entire time points of rs-fMRI data
- Carpet plots of preprocessed rs-fMRI data (Power, 2017) with and without GSR
- Multi-axial plot of the mean rs-fMRI data after motion and distortion correction, with contours of T1 after alignment
- Multi-axial plot of AICHA regional occupancy map, showing the percentage of the functional brain mask within each of the AICHA region in the stereotaxic space
- Similar multi-axial plot of AICHA regional occupancy map, but showing the percentage of GM/WM/ventricles within each AICHA region, using the Freesurfer-based masks in the stereotaxic space.

(A) Multi-trace motion and mean signal plots and carpet plots with and without GSR

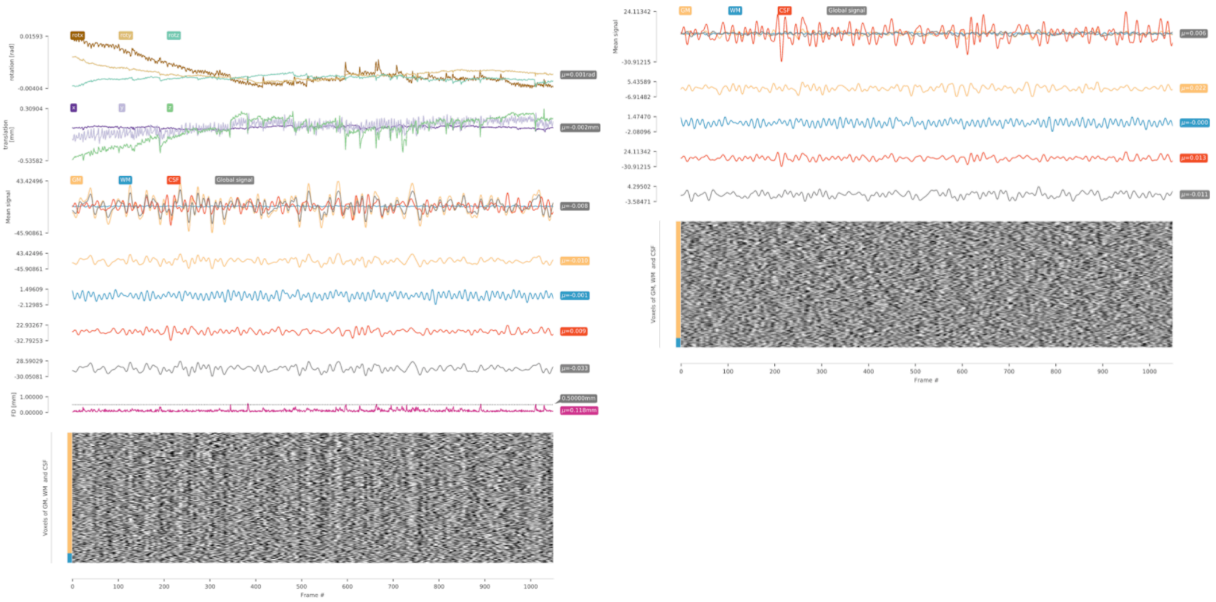

(B) EPI to T1 coregistration contours

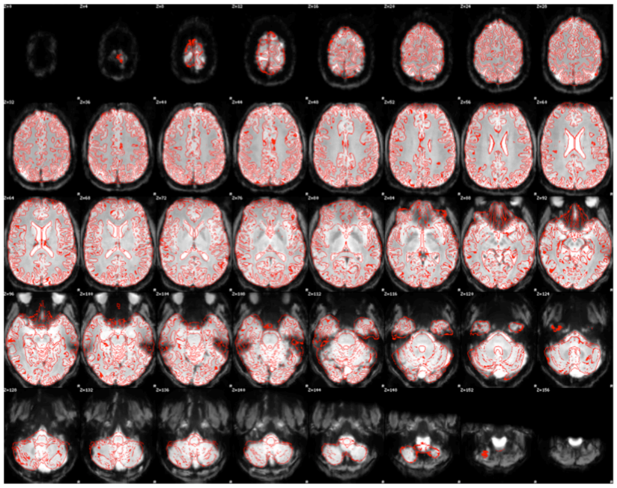

(C) AICHA regional occupancy map

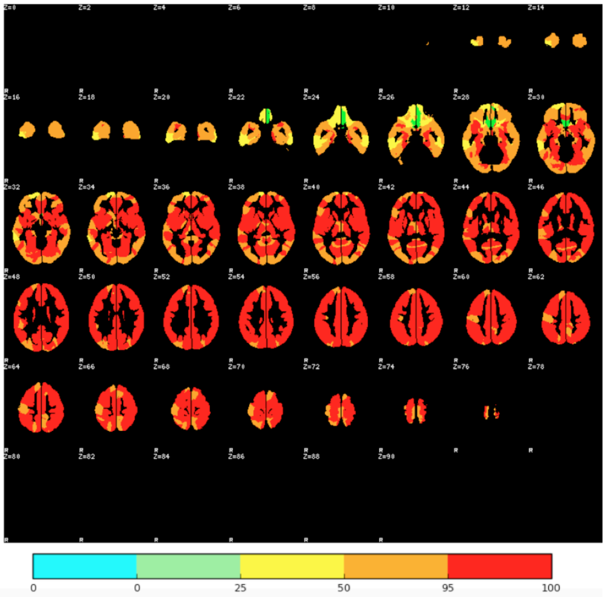

**Supplemental Figure 14. Example of subject-level visual QC images for resting-state fMRI pipeline.**

*(A) Multi-trace plots showing, from the top of the left panel, 1) absolute rotation, 2) absolute translation, 3) global mean signal and signals within GM/WM/ventricles, 4) FD, and 5) carpet plot of the preprocessed (motion- and distortion-corrected, band-passed) rs-fMRI data. The mean signal time courses and the carpet plot in the left and right panel were derived from data without and with GSR, respectively. (B) Multi-axial plot of the average pre-processed data aligned to T1, together with the contour of the reference T1 image. (C) Multi-axial plot of AICHA regional occupancy map, showing the percentage of the functional brain mask within each AICHA region. Similar AICHA occupancy maps were created that show the percentages of GM/WM/ventricles within each AICHA region.*

Also as in DWI pipeline, we focused on identifying the outliers in the quantitative QC metrics and checking their visual QC web pages rather than reviewing every single subject-level QC page. However, we note that any excessive motion was censured through 'scrubbing' described above when computing IC matrices. Also, some subjects were excluded on a regional basis when computing the average regions IC matrices. Here we present the distributions of the following quantitative QC metrics (Supplemental Figure 15) to be compared with other studies:

- Cost function values for EPI to T1 after the BBR alignment to T1, as well as after BBR + FLIRT alignment to T1
- Mean relative RMS displacement, as well as mean and max absolute translation and rotation (mean over each x, y, and z directions), over the entire time course of the rs-fMRI data estimated by MCFLIRT
- Inverse tSNR before and after band-pass filtering that also included despiking and nuisance regressor removal
- Mean FD (Power et al., 2012) and the proportion of volumes above FD above 0.2 and 0.5 (Power et al., 2015) over the entire time course of the rs-fMRI data

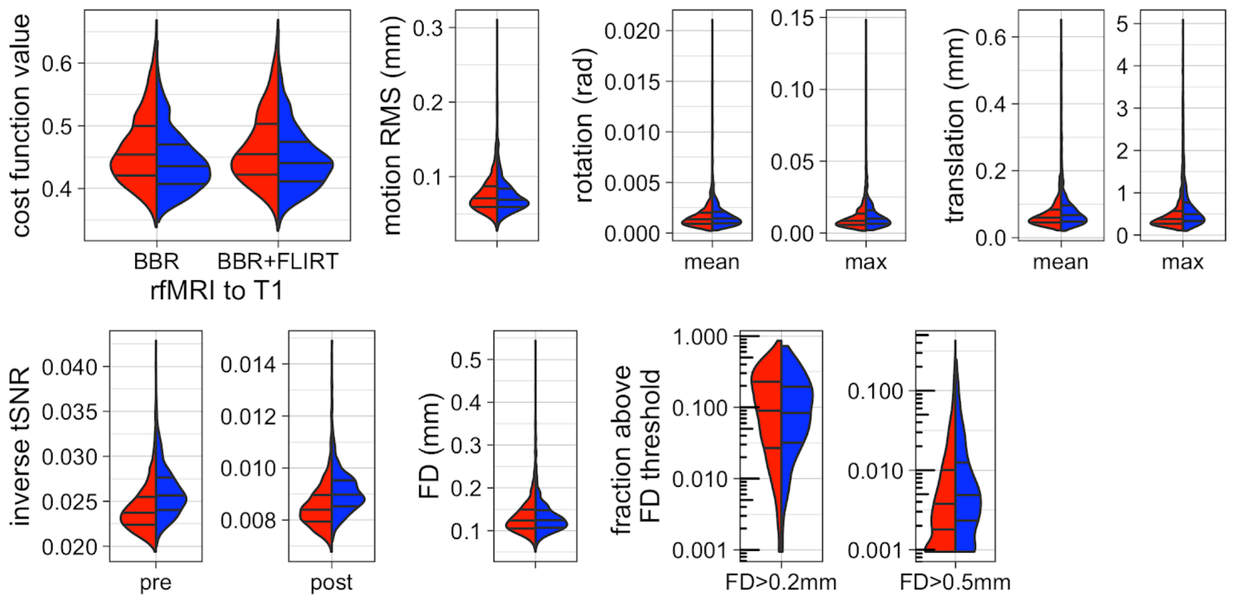

**Supplemental Figure 15. Distributions of quantitative QC metrics for resting-state fMRI pipeline.**

*Distributions of QC IDPs from the rs-fMRI pipeline are shown for male (blue) and female (red) participants. See text for details of individual IDPs. Note that log-scale is used for the fraction of rs-fMRI volumes above the given FD threshold, since the distributions for both thresholds are highly skewed*

### Additional statistical analyses

#### Sample distribution of some global IDP's

Supplemental Figure 16 shows the distributions of global IDPs presented in the current paper, namely mean CT, total inner and pial CSA, and total GM and WM volumes (Supplemental Figure 16A) from the structural pipeline, and mean values of four DTI metrics (FA, MD, AD, and RD) and three NODDI metrics (NDI, ODI, and IsoVF) within the cerebral WM (Supplemental Figure 16B).

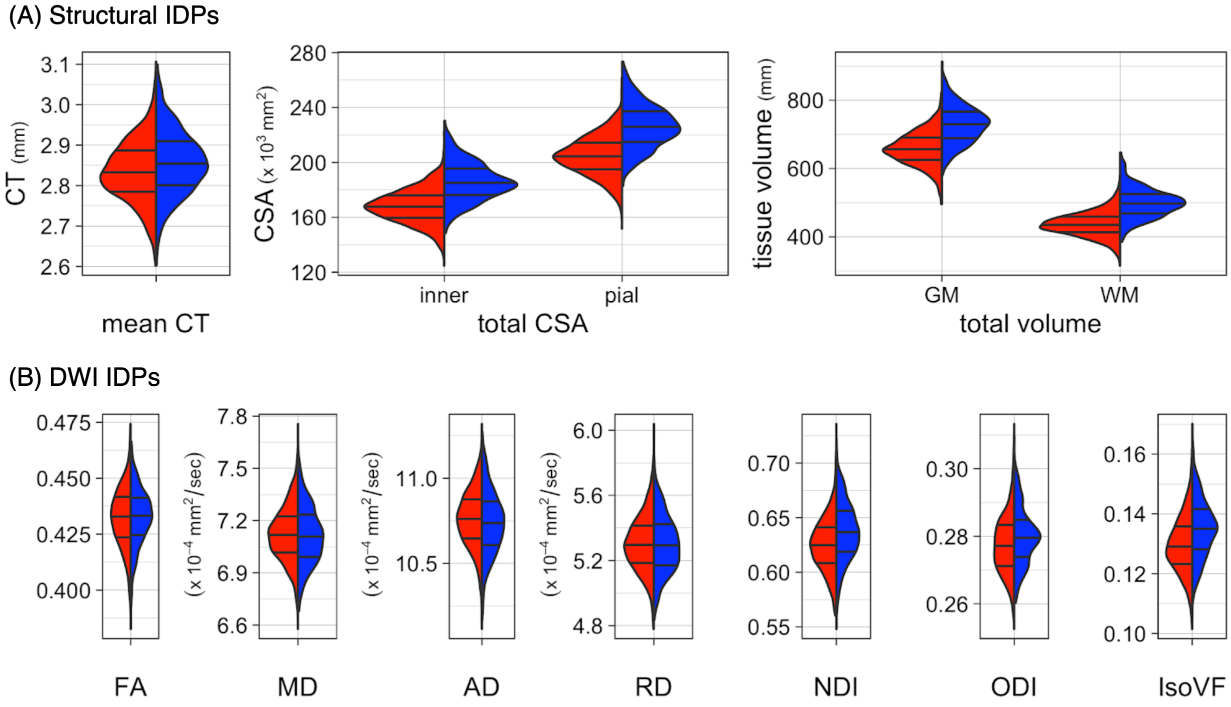

**Supplemental Figure 16. Distributions of global IDPs from the structural and diffusion pipelines.**

*Distributions of (A) five IDPs produced by the structural pipeline and (B) seven IDPs from the diffusion pipeline are presented for male (blue) and female (red) participants.*

#### Correlation of total inner and pial CSA

While most studies report the inner CSA defined by GM/WM interface, and this is the default cortical areal measure in Freesurfer, it is possible to compute the pial CSA defined by GM/CSF interface. Not surprisingly, the total inner and pial CSA were highly correlated in our data, with pial CSA being larger than inner CSA. Supplemental Figure 17 shows the relationship between the two variables in 1,722 subjects of the MRi-Share data.

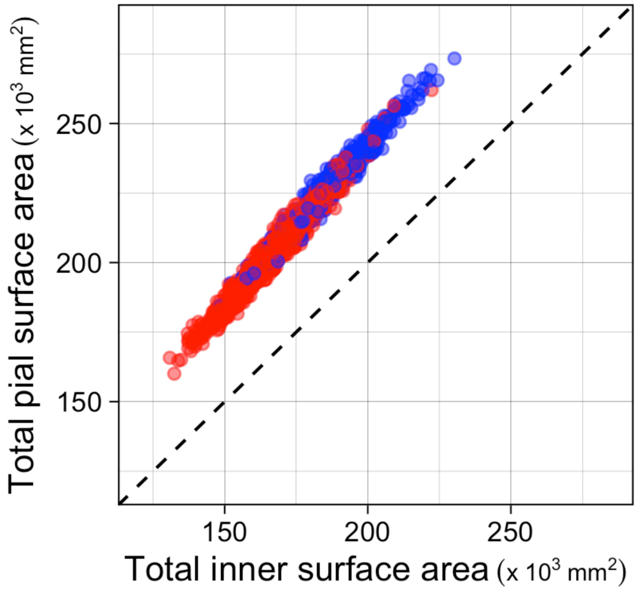

**Supplemental Figure 17. The relationship between CSA and CSA pial in 1,722 subjects.** The plot shows the total inner and pial CSA for each subject, represented as a dot in the scatter plot. The color represents the sex of the subject (blue for males and red for females).

#### The interaction of eTIV and sex in WM volume data

The combined sex analysis showed the significant interaction between eTIV and sex (*p* = 0.0004), indicating that the slope for males is steeper than females. Supplemental Figure 18 shows the small difference in the slopes of the WM volume-to-eTIV in males and females.

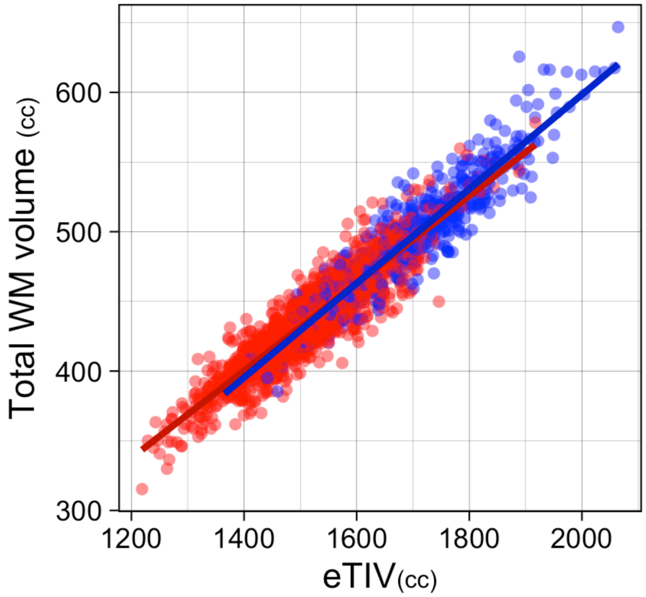

**Supplemental Figure 18. The interaction of eTIV and sex in WM volume data.**

*The plot shows the relationship between total WM volume and eTIV in both sexes. Males show slightly steeper slope than for females.*

#### Fit results of alternative models in the global GM, WM morphometry and WM property analyses

In the main manuscript, we primarily reported the results of linear age effect models with or without eTIV (or WM mask volume in the case of WM properties) that best describe the data according to BIC. However, here we report the results the alternative model fit results to assess how the inclusion or non-inclusion of global volume effects in the model affects the observed age or sex effects, and to allow comparison with any prior studies that did or did not control for such global volume effects.

##### Global gray matter morphometry

The BIC indicated that inclusion of eTIV did not improve overall fit for modeling CT, while it did for inner CSA, pial CSA, and GM volume data. Here we report the model fit results with eTIV for CT, and without eTIV for inner CSA, pial CSA, and GM in Supplemental Table 1. The estimates of age or sex effects are not much affected by the inclusion of eTIV. In contrast, excluding eTIV slightly reduces the estimates of age-related variance for both inner and pial CSA, and GM, decreasing the level of significance for the age effects in these metrics, although the estimated slope for the age tends to increase, in particular for inner and pial CSA data in males. Not surprisingly, the exclusion of eTIV also significantly impacts the observed sex effects on these metrics, as males generally have larger total eTIV and, as a result, larger raw values for inner and pial CSA as well as GM, than females. These absolute differences in the sexes disappear and in some cases reverse the effects when eTIV is accounted for in the model.

**Supplemental Table 1. The effects of inclusion or exclusion of eTIV on the age and sex effects on global GM morphometry.**

*The results of alternative linear model fit for mean CT, total inner CSA, and total pial CSA from Freesurfer v6.0, as well as total GM volume from SPM12, are shown. Parameter estimates (β) for age and sex are shown, together with 95% confidence intervals (CI) for each β, as well as uncorrected p values and partial eta squared as the effect sizes for each variable. The Model column indicates the alternative model (see text) with the corresponding model number as described in Methods section. The interaction between sex and age was tested in the combined group but was not significant in any of the metrics listed here, and thus is not included in the table. Overall model fit is indicated as adjusted squared R values.*

| | | | Age | | | Sex (M>F) | | | |
|---|---|---|---|---|---|---|---|---|---|
| | ***N*** | **Model** | β | *p* | partial $\eta^2$ | β | *p* | partial $\eta^2$ | **adj $R^2$** |
| **CT** | | | ($x10^{-3}$ mm/yr) | | | ($x10^{-3}$ mm) | | | |
| Male | 470 | (2) | -2.47 [-6.46, 1.53] | 0.23 | 0.003 | -- | | -- | -0.001 |
| Female | 1252 | (2) | -4.56 [-6.95, -2.18] | $1.8x10^{-4}$ | 0.011 | -- | | -- | -0.001 |
| Combined | 1722 | (4) | -3.52 [-5.76, -1.27] | 0.0022 | 0.008 | 21.8 [10.1, 33.5] | $<2.0x10^{-16}$ | 0.009 | 0.024 |
| **inner CSA** | | | ($x10^2$ $mm^2$/yr) | | | ($x10^2$ $mm^2$) | | | |
| Male | 470 | (1) | -7.51 [-14.3, -0.68] | 0.031 | 0.010 | -- | | -- | 0.008 |
| Female | 1252 | (1) | -2.26 [-6.20, 1.67] | 0.26 | 0.001 | -- | | -- | 0.000 |
| Combined | 1722 | (3) | -4.89 [-8.63, -1.14] | 0.011 | 0.003 | 180.4 [166.8, 193.9] | $<2.0x10^{-16}$ | 0.284 | 0.028 |
| **pial CSA** | | | ($x10^2$ $mm^2$/yr) | | | ($x10^2$ $mm^2$) | | | |
| Male | 470 | (1) | -11.3 [-19.1, -3.44] | 0.0049 | 0.017 | -- | | -- | 0.015 |
| Female | 1252 | (1) | -5.06 [-9.66, -0.47] | 0.031 | 0.004 | -- | | -- | 0.003 |
| Combined | 1722 | (3) | -3.10 [-12.5, -3.81] | $2.4x10^{-4}$ | 0.007 | 217.7 [202.0, 233.3] | $<2.0x10^{-16}$ | 0.301 | 0.301 |
| **GM** | | | (cc/yr) | | | (cc) | | | |
| Male | 470 | (1) | -3.47 [-6.17, -0.78] | 0.012 | 0.014 | -- | | -- | 0.011 |
| Female | 1252 | (1) | -2.63 [-4.19, -1.07] | $9.9x10^{-4}$ | 0.009 | -- | | -- | 0.008 |
| Combined | 1722 | (3) | -3.05 [-4.54, -1.57] | $5.8x10^{-5}$ | 0.01 | 72.2 [66.8, 77.5] | $<2.0x10^{-16}$ | 0.010 | 0.291 |

#### Global white matter morphometry and properties

Like GM, the total WM volume was also significantly affected by eTIV. For the mean DTI metrics within the cerebral WM mask, BIC indicated that the inclusion of mask volume in the model improved the model sufficiently for MD and AD in male, but not in female, data. Here we report the alternative model results in Supplemental Table 2.

**Supplemental Table 2. The effects of inclusion or exclusion of eTIV or the WM mask volume on the age and sex effects on WM volume and DTI data.**

*The results of alternative linear model fit for total white matter volume (WM) from SPM12, as well as mean DTI metrics within subject-specific cerebral WM masks are shown. Parameter estimates (β) for age and sex are shown, together with 95% confidence intervals (CI) for each β, as well as uncorrected p values and partial eta squared as the effect sizes for each variable. The Model column indicates the alternative model (see text), with the corresponding model number as described in Methods section. The interaction between sex and age was tested in the combined group but was not significant in any of the metrics listed here, and thus is not included in the table. Overall model fit is indicated as adjusted squared R values*.

| | | | Age | | | Sex (M>F) | | | |
|---|---|---|---|---|---|---|---|---|---|
| | ***N*** | **Model** | β | *p* | partial $\eta^2$ | β | *p* | partial $\eta^2$ | **adj $R^2$** |
| **CT** | | | (x$10^{-3}$ mm/yr) | | | (x$10^{-3}$ mm) | | | |
| Male | 470 | (2) | -2.47 [-6.46, 1.53] | 0.23 | 0.003 | -- | | -- | -0.001 |
| Female | 1252 | (2) | -4.56 [-6.95, -2.18] | 1.8x$10^{-4}$ | 0.011 | -- | | -- | -0.001 |
| Combined | 1722 | (4) | -3.52 [-5.76, -1.27] | 0.0022 | 0.008 | 21.8 [10.1, 33.5] | <2.0x$10^{-16}$ | 0.009 | 0.024 |
| **inner CSA** | | | (x$10^2$ mm²/yr) | | | (x$10^2$ mm²) | | | |
| Male | 470 | (1) | -7.51 [-14.3, -0.68] | 0.031 | 0.010 | -- | | -- | 0.008 |
| Female | 1252 | (1) | -2.26 [-6.20, 1.67] | 0.26 | 0.001 | -- | | -- | 0.000 |
| Combined | 1722 | (3) | -4.89 [-8.63, -1.14] | 0.011 | 0.003 | 180.4 [166.8, 193.9] | <2.0x$10^{-16}$ | 0.284 | 0.028 |
| **pial CSA** | | | (x$10^2$ mm²/yr) | | | (x$10^2$ mm²) | | | |
| Male | 470 | (1) | -11.3 [-19.1, -3.44] | 0.0049 | 0.017 | -- | | -- | 0.015 |
| Female | 1252 | (1) | -5.06 [-9.66, -0.47] | 0.031 | 0.004 | -- | | -- | 0.003 |
| Combined | 1722 | (3) | -3.10 [-12.5, -3.81] | 2.4x$10^{-4}$ | 0.007 | 217.7 [202.0, 233.3] | <2.0x$10^{-16}$ | 0.301 | 0.301 |
| **GM** | | | (cc/yr) | | | (cc) | | | |
| Male | 470 | (1) | -3.47 [-6.17, -0.78] | 0.012 | 0.014 | -- | | -- | 0.011 |
| Female | 1252 | (1) | -2.63 [-4.19, -1.07] | 9.9x$10^{-4}$ | 0.009 | -- | | -- | 0.008 |
| Combined | 1722 | (3) | -3.05 [-4.54, -1.57] | 5.8x$10^{-5}$ | 0.01 | 72.2 [66.8, 77.5] | <2.0x$10^{-16}$ | 0.010 | 0.291 |

When not taking eTIV into account, the total WM volume no longer showed any significant positive age-related variations, and the age-related variance in the data was much reduced. As in GM, it also affected the observed sex effects, indicating a significantly larger raw WM values in males than in females, which disappears once eTIV is taken into account.

Inclusion or exclusion of the WM mask volume did not impact the estimates of age effects in any of the mean DTI metrics. It did slightly affect the observed sex effects in the diffusivity measures, with a tendency for higher diffusivity in females than in males when controlling for the mask volume but no difference when not controlling for the volume.

For the mean NODDI metrics, BIC indicated the inclusion of WM mask volume in the model to improve the fit for IsoVF only, and not for NDI or ODI. The Supplemental Table 3 reports the alternative model results, showing the results with the mask volume in the model for NDI and ODI, and without the mask volume for IsoVF. The inclusion or exclusion of the mask volume did not noticeably affect either age or sex effects in any of the NODDI metrics, although the sex difference in the mean IsoVF (higher in males than in females) seems to be attenuated slightly when not taking into account the mask volume.

**Supplemental Table 3. The effects of inclusion or exclusion of the WM mask volume on the age and sex effects on NODDI data.**

*The results of alternative linear model fit for mean NODDI metrics within subject-specific cerebral WM masks are shown. Parameter estimates (β) for age and sex are shown, together with 95% confidence intervals (CI) for each β, as well as uncorrected p values and partial eta squared as the effect sizes for each variable. The Model column indicates the alternative model (see text) with the corresponding model number as described in Methods section. The interaction between sex and age was tested in the combined group but was not significant in any of the metrics listed here, and thus is not included in the table. Overall model fit is indicated as adjusted squared R values.*

| | | | Age | | | Sex (M>F) | | | |
|---|---|---|---|---|---|---|---|---|---|
| | *N* | **Model** | β [95% CI] | *p* | partial $\eta^2$ | β [95% CI] | *p* | partial $\eta^2$ | **adj $R^2$** |
| **NDI** | | | ($\times 10^{-3}$ /yr) | | | ($\times 10^{-3}$) | | | |
| Male | 468 | (2) | 3.65 [2.37, 4.94] | $4.1\times 10^{-8}$ | 0.063 | -- | -- | -- | 0.060 |
| Female | 1246 | (2) | 2.51 [1.74, 3.27] | $1.7\times 10^{-10}$ | 0.032 | -- | -- | -- | 0.034 |
| Combined | 1714 | (4) | 3.08 [2.36, 3.80] | $<2.0\times 10^{-16}$ | 0.041 | 11.0 [6.95, 13.7] | $2.5\times 10^{-9}$ | 0.023 | 0.083 |
| **ODI** | | | ($\times 10^{-4}$/yr) | | | ($\times 10^{-4}$) | | | |
| Male | 468 | (2) | 6.69 [2.61, 10.8] | 0.0014 | 0.022 | -- | -- | -- | 0.018 |
| Female | 1246 | (2) | 6.45 [3.66, 9.25] | $6.5\times 10^{-6}$ | 0.016 | -- | -- | -- | 0.015 |
| Combined | 1714 | (4) | 6.57 [4.04, 9.10] | $4.0\times 10^{-7}$ | 0.018 | 22.1 [10.2, 34.0] | $2.7\times 10^{-4}$ | 0.009 | 0.026 |
| **IsoVF** | | | ($\times 10^{-4}$/yr) | | | ($\times 10^{-4}$) | | | |
| Male | 468 | (1) | 3.16 [-1.76, 8.08] | 0.21 | 0.003 | -- | -- | -- | 0.001 |
| Female | 1246 | (1) | 3.65 [0.66, 6.64] | 0.017 | 0.005 | -- | -- | -- | 0.004 |
| Combined | 1714 | (3) | 3.41 [0.61, 6.21] | 0.017 | 0.004 | 55.6 [45.4, 65.7] | $<2.0\times 10^{-16}$ | 0.064 | 0.067 |

### References

Alfaro-Almagro, F., Jenkinson, M., Bangerter, N.K., Andersson, J.L.R., Griffanti, L., Douaud, G., Sotiropoulos, S.N., Jbabdi, S., Hernandez-Fernandez, M., Vallee, E., Vidaurre, D., Webster, M., McCarthy, P., Rorden, C., Daducci, A., Alexander, D.C., Zhang, H., Dragonu, I., Matthews, P.M., Miller, K.L., Smith, S.M., 2018. Image processing and Quality Control for the first 10,000 brain imaging datasets from UK Biobank. Neuroimage 166, 400–424. doi:10.1016/j.neuroimage.2017.10.034

Andersson, J.L.R., Graham, M.S., Zsoldos, E., Sotiropoulos, S.N., 2016. Incorporating outlier detection and replacement into a non-parametric framework for movement and distortion correction of diffusion MR images. Neuroimage 141, 556–572. doi:10.1016/j.neuroimage.2016.06.058

Andersson, J.L.R., Skare, S., Ashburner, J., 2003. How to correct susceptibility distortions in spin-echo echo-planar images: application to diffusion tensor imaging. Neuroimage 20, 870–888. doi:10.1016/S1053-8119(03)00336-7

Andersson, J.L.R., Sotiropoulos, S.N., 2016. An integrated approach to correction for off-resonance effects and subject movement in diffusion MR imaging. Neuroimage 125, 1063–1078. doi:10.1016/j.neuroimage.2015.10.019

Ashburner, J., Barnes, G., Chen, C.-C., Daunizeau, J., Flandin, G., Friston, K., Kiebel, S., Kilner, J., Litvak, V., Moran, R., Penny, W., Razi, A., Stephan, K., Tak, S., Zeidman, P., Gitelman, D., Henson, R., Hutton, C., Glauche, V., Mattout, J., Phillips, C., 2014. SPM12 Manual.

Ashburner, J., Friston, K.J., 2005. Unified segmentation. Neuroimage 26, 839–851. doi:10.1016/j.neuroimage.2005.02.018

Avants, B.B., Epstein, C.L., Grossman, M., Gee, J.C., 2008. Symmetric diffeomorphic image registration with cross-correlation: evaluating automated labeling of elderly and neurodegenerative brain. Med. Image Anal. 12, 26–41. doi:10.1016/j.media.2007.06.004

Backhausen, L.L., Herting, M.M., Buse, J., Roessner, V., Smolka, M.N., Vetter, N.C., 2016. Quality Control of Structural MRI Images Applied Using FreeSurfer-A Hands-On Workflow to Rate Motion Artifacts. Front. Neurosci. 10, 558. doi:10.3389/fnins.2016.00558

Basser, P.J., Mattiello, J., LeBihan, D., 1994. MR diffusion tensor spectroscopy and imaging. Biophys. J. 66, 259–267. doi:10.1016/S0006-3495(94)80775-1

Bastiani, M., Cottaar, M., Fitzgibbon, S.P., Suri, S., Alfaro-Almagro, F., Sotiropoulos, S.N., Jbabdi, S., Andersson, J.L.R., 2019. Automated quality control for within and between studies diffusion MRI data using a non-parametric framework for movement and distortion correction. Neuroimage 184, 801–812. doi:10.1016/j.neuroimage.2018.09.073

Baykara, E., Gesierich, B., Adam, R., Tuladhar, A.M., Biesbroek, J.M., Koek, H.L., Ropele, S., Jouvent, E., Alzheimer's Disease Neuroimaging Initiative, Chabriat, H., Ertl-Wagner, B., Ewers, M., Schmidt, R., de Leeuw, F.-E., Biessels, G.J., Dichgans, M., Duering, M., 2016. A novel imaging marker for small vessel disease based on skeletonization of white matter tracts and diffusion histograms. Ann. Neurol. 80, 581–592. doi:10.1002/ana.24758

Beaudet, G., Tsuchida, A., Petit, L., Tzourio, C., Caspers, S., Schreiber, J., Pausova, Z., Patel, Y., Paus, T., Schmidt, R., Pirpamer, L., Sachdev, P.S., Brodaty, H., Kochan, N., Trollor, J., Wen, W., Armstrong, N.J., Deary, I.J., Bastin, M.E., Wardlaw, J.M., Mazoyer, B., 2020. Age-Related Changes of Peak Width Skeletonized Mean Diffusivity (PSMD) Across the Adult Lifespan: A Multi-Cohort Study. Front. Psychiatry 11, 342. doi:10.3389/fpsyt.2020.00342

Beckmann, C.F., Smith, S.M., 2004. Probabilistic independent component analysis for functional magnetic resonance imaging. IEEE Trans. Med. Imaging 23, 137–152. doi:10.1109/TMI.2003.822821

Choi, S., Kim, T., Yu, W., 2009. Performance evaluation of RANSAC family, in: Procedings of the British Machine Vision Conference 2009. Presented at the British Machine Vision

Conference 2009, British Machine Vision Association, pp. 81.1-81.12. doi:10.5244/C.23.81
Coupe, P., Manjon, J., Robles, M., Collins, L.D., 2011. Adaptive Multiresolution Non-Local Means Filter for 3D MR Image Denoising. IET Image Processing.
Coupe, P., Yger, P., Prima, S., Hellier, P., Kervrann, C., Barillot, C., 2008. An optimized blockwise nonlocal means denoising filter for 3-D magnetic resonance images. IEEE Trans. Med. Imaging 27, 425–441. doi:10.1109/TMI.2007.906087
Daducci, A., Canales-Rodríguez, E.J., Zhang, H., Dyrby, T.B., Alexander, D.C., Thiran, J.-P., 2015. Accelerated Microstructure Imaging via Convex Optimization (AMICO) from diffusion MRI data. Neuroimage 105, 32–44. doi:10.1016/j.neuroimage.2014.10.026
Dale, A.M., Fischl, B., Sereno, M.I., 1999. Cortical surface-based analysis. I. Segmentation and surface reconstruction. Neuroimage 9, 179–194. doi:10.1006/nimg.1998.0395
Desikan, R.S., Ségonne, F., Fischl, B., Quinn, B.T., Dickerson, B.C., Blacker, D., Buckner, R.L., Dale, A.M., Maguire, R.P., Hyman, B.T., Albert, M.S., Killiany, R.J., 2006. An automated labeling system for subdividing the human cerebral cortex on MRI scans into gyral based regions of interest. Neuroimage 31, 968–980. doi:10.1016/j.neuroimage.2006.01.021
Diedrichsen, J., Balsters, J.H., Flavell, J., Cussans, E., Ramnani, N., 2009. A probabilistic MR atlas of the human cerebellum. Neuroimage 46, 39–46. doi:10.1016/j.neuroimage.2009.01.045
Fischl, B., Salat, D.H., Busa, E., Albert, M., Dieterich, M., Haselgrove, C., van der Kouwe, A., Killiany, R., Kennedy, D., Klaveness, S., Montillo, A., Makris, N., Rosen, B., Dale, A.M., 2002. Whole Brain Segmentation. Neuron 33, 341–355. doi:10.1016/S0896-6273(02)00569-X
Fischl, B., Salat, D.H., van der Kouwe, A.J.W., Makris, N., Ségonne, F., Quinn, B.T., Dale, A.M., 2004. Sequence-independent segmentation of magnetic resonance images. Neuroimage 23 Suppl 1, S69-84. doi:10.1016/j.neuroimage.2004.07.016
Frazier, J.A., Chiu, S., Breeze, J.L., Makris, N., Lange, N., Kennedy, D.N., Herbert, M.R., Bent, E.K., Koneru, V.K., Dieterich, M.E., Hodge, S.M., Rauch, S.L., Grant, P.E., Cohen, B.M., Seidman, L.J., Caviness, V.S., Biederman, J., 2005. Structural brain magnetic resonance imaging of limbic and thalamic volumes in pediatric bipolar disorder. Am. J. Psychiatry 162, 1256–1265. doi:10.1176/appi.ajp.162.7.1256
Friston, K.J., Williams, S., Howard, R., Frackowiak, R.S., Turner, R., 1996. Movement-related effects in fMRI time-series. Magn. Reson. Med. 35, 346–355. doi:10.1002/mrm.1910350312
Ganzetti, M., Wenderoth, N., Mantini, D., 2016a. Quantitative evaluation of intensity inhomogeneity correction methods for structural MR brain images. Neuroinformatics 14, 5–21. doi:10.1007/s12021-015-9277-2
Ganzetti, M., Wenderoth, N., Mantini, D., 2016b. Intensity Inhomogeneity Correction of Structural MR Images: A Data-Driven Approach to Define Input Algorithm Parameters. Front. Neuroinformatics 10, 10. doi:10.3389/fninf.2016.00010
Garyfallidis, E., Brett, M., Amirbekian, B., Rokem, A., van der Walt, S., Descoteaux, M., Nimmo-Smith, I., Dipy Contributors, 2014. Dipy, a library for the analysis of diffusion MRI data. Front. Neuroinformatics 8, 8. doi:10.3389/fninf.2014.00008
Goldstein, J.M., Seidman, L.J., Makris, N., Ahern, T., O'Brien, L.M., Caviness, V.S., Kennedy, D.N., Faraone, S.V., Tsuang, M.T., 2007. Hypothalamic abnormalities in schizophrenia: sex

effects and genetic vulnerability. Biol. Psychiatry 61, 935–945. doi:10.1016/j.biopsych.2006.06.027
Haddad, S.M.H., Scott, C.J.M., Ozzoude, M., Holmes, M.F., Arnott, S.R., Nanayakkara, N.D., Ramirez, J., Black, S.E., Dowlatshahi, D., Strother, S.C., Swartz, R.H., Symons, S., Montero-Odasso, M., ONDRI Investigators, Bartha, R., 2019. Comparison of quality control methods for automated diffusion tensor imaging analysis pipelines. PLoS ONE 14, e0226715. doi:10.1371/journal.pone.0226715
Iglesias, J.E., Augustinack, J.C., Nguyen, K., Player, C.M., Player, A., Wright, M., Roy, N., Frosch, M.P., McKee, A.C., Wald, L.L., Fischl, B., Van Leemput, K., Alzheimer's Disease Neuroimaging Initiative, 2015a. A computational atlas of the hippocampal formation using ex vivo, ultra-high resolution MRI: Application to adaptive segmentation of in vivo MRI. Neuroimage 115, 117–137. doi:10.1016/j.neuroimage.2015.04.042
Iglesias, J.E., Van Leemput, K., Bhatt, P., Casillas, C., Dutt, S., Schuff, N., Truran-Sacrey, D., Boxer, A., Fischl, B., Alzheimer's Disease Neuroimaging Initiative, 2015b. Bayesian segmentation of brainstem structures in MRI. Neuroimage 113, 184–195. doi:10.1016/j.neuroimage.2015.02.065
Jenkinson, M., Bannister, P., Brady, M., Smith, S., 2002. Improved optimization for the robust and accurate linear registration and motion correction of brain images. Neuroimage 17, 825–841. doi:10.1016/s1053-8119(02)91132-8
Jenkinson, M., Smith, S., 2001. A global optimisation method for robust affine registration of brain images. Med. Image Anal. 5, 143–156. doi:10.1016/s1361-8415(01)00036-6
Jensen, J.H., Helpern, J.A., 2010. MRI quantification of non-Gaussian water diffusion by kurtosis analysis. NMR Biomed. 23, 698–710. doi:10.1002/nbm.1518
Joliot, M., Jobard, G., Naveau, M., Delcroix, N., Petit, L., Zago, L., Crivello, F., Mellet, E., Mazoyer, B., Tzourio-Mazoyer, N., 2015. AICHA: An atlas of intrinsic connectivity of homotopic areas. J. Neurosci. Methods 254, 46–59. doi:10.1016/j.jneumeth.2015.07.013
Klapwijk, E.T., van de Kamp, F., van der Meulen, M., Peters, S., Wierenga, L.M., 2019. Qoala-T: A supervised-learning tool for quality control of FreeSurfer segmented MRI data. Neuroimage 189, 116–129. doi:10.1016/j.neuroimage.2019.01.014
Lindig, T., Kotikalapudi, R., Schweikardt, D., Martin, P., Bender, F., Klose, U., Ernemann, U., Focke, N.K., Bender, B., 2018. Evaluation of multimodal segmentation based on 3D T1-, T2- and FLAIR-weighted images - the difficulty of choosing. Neuroimage 170, 210–221. doi:10.1016/j.neuroimage.2017.02.016
Liu, B., Zhu, T., Zhong, J., 2015. Comparison of quality control software tools for diffusion tensor imaging. Magn. Reson. Imaging 33, 276–285. doi:10.1016/j.mri.2014.10.011
Liu, Z., Wang, Y., Gerig, G., Gouttard, S., Tao, R., Fletcher, T., Styner, M., 2010. Quality control of diffusion weighted images. Proc. SPIE 7628. doi:10.1117/12.844748
Madan, C.R., 2018. Age differences in head motion and estimates of cortical morphology. PeerJ 6, e5176. doi:10.7717/peerj.5176
Makris, N., Goldstein, J.M., Kennedy, D., Hodge, S.M., Caviness, V.S., Faraone, S.V., Tsuang, M.T., Seidman, L.J., 2006. Decreased volume of left and total anterior insular lobule in schizophrenia. Schizophr. Res. 83, 155–171. doi:10.1016/j.schres.2005.11.020
Minka, T., 2000. Automatic choice of dimensionality for PCA. (Technical Report No. 514). MIT Media Lab Vision and Modeling Group.

Mori, S., Oishi, K., Jiang, H., Jiang, L., Li, X., Akhter, K., Hua, K., Faria, A.V., Mahmood, A., Woods, R., Toga, A.W., Pike, G.B., Neto, P.R., Evans, A., Zhang, J., Huang, H., Miller, M.I., van Zijl, P., Mazziotta, J., 2008. Stereotaxic white matter atlas based on diffusion tensor imaging in an ICBM template. Neuroimage 40, 570–582. doi:10.1016/j.neuroimage.2007.12.035

Oguz, I., Farzinfar, M., Matsui, J., Budin, F., Liu, Z., Gerig, G., Johnson, H.J., Styner, M., 2014. DTIPrep: quality control of diffusion-weighted images. Front. Neuroinformatics 8, 4. doi:10.3389/fninf.2014.00004

Oishi, K., Zilles, K., Amunts, K., Faria, A., Jiang, H., Li, X., Akhter, K., Hua, K., Woods, R., Toga, A.W., Pike, G.B., Rosa-Neto, P., Evans, A., Zhang, J., Huang, H., Miller, M.I., van Zijl, P.C.M., Mazziotta, J., Mori, S., 2008. Human brain white matter atlas: identification and assignment of common anatomical structures in superficial white matter. Neuroimage 43, 447–457. doi:10.1016/j.neuroimage.2008.07.009

Power, J.D., 2017. A simple but useful way to assess fMRI scan qualities. Neuroimage 154, 150–158. doi:10.1016/j.neuroimage.2016.08.009

Power, J.D., Barnes, K.A., Snyder, A.Z., Schlaggar, B.L., Petersen, S.E., 2012. Spurious but systematic correlations in functional connectivity MRI networks arise from subject motion. Neuroimage 59, 2142–2154. doi:10.1016/j.neuroimage.2011.10.018

Power, J.D., Mitra, A., Laumann, T.O., Snyder, A.Z., Schlaggar, B.L., Petersen, S.E., 2014. Methods to detect, characterize, and remove motion artifact in resting state fMRI. Neuroimage 84, 320–341. doi:10.1016/j.neuroimage.2013.08.048

Power, J.D., Schlaggar, B.L., Petersen, S.E., 2015. Recent progress and outstanding issues in motion correction in resting state fMRI. Neuroimage 105, 536–551. doi:10.1016/j.neuroimage.2014.10.044

Reuter, M., Tisdall, M.D., Qureshi, A., Buckner, R.L., van der Kouwe, A.J.W., Fischl, B., 2015. Head motion during MRI acquisition reduces gray matter volume and thickness estimates. Neuroimage 107, 107–115. doi:10.1016/j.neuroimage.2014.12.006

Roalf, D.R., Quarmley, M., Elliott, M.A., Satterthwaite, T.D., Vandekar, S.N., Ruparel, K., Gennatas, E.D., Calkins, M.E., Moore, T.M., Hopson, R., Prabhakaran, K., Jackson, C.T., Verma, R., Hakonarson, H., Gur, R.C., Gur, R.E., 2016. The impact of quality assurance assessment on diffusion tensor imaging outcomes in a large-scale population-based cohort. Neuroimage 125, 903–919. doi:10.1016/j.neuroimage.2015.10.068

Rosen, A.F.G., Roalf, D.R., Ruparel, K., Blake, J., Seelaus, K., Villa, L.P., Ciric, R., Cook, P.A., Davatzikos, C., Elliott, M.A., Garcia de La Garza, A., Gennatas, E.D., Quarmley, M., Schmitt, J.E., Shinohara, R.T., Tisdall, M.D., Craddock, R.C., Gur, R.E., Gur, R.C., Satterthwaite, T.D., 2018. Quantitative assessment of structural image quality. Neuroimage 169, 407–418. doi:10.1016/j.neuroimage.2017.12.059

Savalia, N.K., Agres, P.F., Chan, M.Y., Feczko, E.J., Kennedy, K.M., Wig, G.S., 2017. Motion-related artifacts in structural brain images revealed with independent estimates of in-scanner head motion. Hum. Brain Mapp. 38, 472–492. doi:10.1002/hbm.23397

Sforazzini, F., Bertero, A., Dodero, L., David, G., Galbusera, A., Scattoni, M.L., Pasqualetti, M., Gozzi, A., 2016. Altered functional connectivity networks in acallosal and socially impaired BTBR mice. Brain Struct. Funct. 221, 941–954. doi:10.1007/s00429-014-0948-9

Sled, J.G., Zijdenbos, A.P., Evans, A.C., 1998. A nonparametric method for automatic correction of intensity nonuniformity in MRI data. IEEE Trans. Med. Imaging 17, 87–97. doi:10.1109/42.668698

Smith, S.M., 2002. Fast robust automated brain extraction. Hum. Brain Mapp. 17, 143–155. doi:10.1002/hbm.10062

Smith, S.M., Jenkinson, M., Johansen-Berg, H., Rueckert, D., Nichols, T.E., Mackay, C.E., Watkins, K.E., Ciccarelli, O., Cader, M.Z., Matthews, P.M., Behrens, T.E.J., 2006. Tract-based spatial statistics: voxelwise analysis of multi-subject diffusion data. Neuroimage 31, 1487–1505. doi:10.1016/j.neuroimage.2006.02.024

Tournier, J.-D., Mori, S., Leemans, A., 2011. Diffusion tensor imaging and beyond. Magn. Reson. Med. 65, 1532–1556. doi:10.1002/mrm.22924

Tustison, N.J., Avants, B.B., Cook, P.A., Zheng, Y., Egan, A., Yushkevich, P.A., Gee, J.C., 2010. N4ITK: improved N3 bias correction. IEEE Trans. Med. Imaging 29, 1310–1320. doi:10.1109/TMI.2010.2046908

Yang, H., Long, X.-Y., Yang, Y., Yan, H., Zhu, C.-Z., Zhou, X.-P., Zang, Y.-F., Gong, Q.-Y., 2007. Amplitude of low frequency fluctuation within visual areas revealed by resting-state functional MRI. Neuroimage 36, 144–152. doi:10.1016/j.neuroimage.2007.01.054

Zacà, D., Hasson, U., Minati, L., Jovicich, J., 2018. Method for retrospective estimation of natural head movement during structural MRI. J. Magn. Reson. Imaging 48, 927–937. doi:10.1002/jmri.25959

Zang, Y., Jiang, T., Lu, Y., He, Y., Tian, L., 2004. Regional homogeneity approach to fMRI data analysis. Neuroimage 22, 394–400. doi:10.1016/j.neuroimage.2003.12.030

Zhang, H., Schneider, T., Wheeler-Kingshott, C.A., Alexander, D.C., 2012. NODDI: practical in vivo neurite orientation dispersion and density imaging of the human brain. Neuroimage 61, 1000–1016. doi:10.1016/j.neuroimage.2012.03.072

Zheng, W., Chee, M.W.L., Zagorodnov, V., 2009. Improvement of brain segmentation accuracy by optimizing non-uniformity correction using N3. Neuroimage 48, 73–83. doi:10.1016/j.neuroimage.2009.06.039

Zou, Q.-H., Zhu, C.-Z., Yang, Y., Zuo, X.-N., Long, X.-Y., Cao, Q.-J., Wang, Y.-F., Zang, Y.-F., 2008. An improved approach to detection of amplitude of low-frequency fluctuation (ALFF) for resting-state fMRI: fractional ALFF. J. Neurosci. Methods 172, 137–141. doi:10.1016/j.jneumeth.2008.04.012
